# Single-cell Multiome Analysis of Chromatin State and Transcriptome in the Human Basal Ganglia

**DOI:** 10.64898/2026.02.03.703645

**Authors:** Lei Chang, Kai Li, Yang Xie, Guojie Zhong, Jonathan A. Rink, Cindy Tatiana Báez-Becerra, Audrey Lie, Hannah S Indralingam, Keyi Dong, Timothy Loe, Zhaoning Wang, Songpeng Zu, Joseph Colin Kern, Zoey Zhao, Eric Boone, Jesus Flores, Alexander Monell, Jacqueline Olness, Cesar Barragan, Emma Osgood, William Owens, Natalie Schenker-Ahmed, Wenjin Zhang, Di Liu, Ariana S. Barcoma, Jackson K. Willier, Kyle W. Knutson, Kaitlyn G. Russo, Jiayi Liu, Silvia Cho, Jessica Arzavala, Carissa K. Young, Guha V. Sundaram, Austin C. Manning, Yareli Sanchez, Aleksandra Bikkina, Jillian Berry, Xiaomeng Gao, Carolyn O’Connor, Michelle Liem, Mikayla V. Marrin, Cynthia Rose, Shane N. Alt, Chenxu Zhu, Nathan R Zemke, Wubin Ding, Amit Klein, Yuanyuan Fu, Nelson Johansen, Trygve E. Bakken, Rebecca D. Hodge, C. Dirk Keene, Ed S. Lein, Daofeng Li, Quan Zhu, Ting Wang, Xiangmin Xu, Joseph R. Ecker, M. Margarita Behrens, Bing Ren

## Abstract

The basal ganglia play essential roles in motor control, emotion, learning and reward processing. Their dysfunction contributes to many neurological and psychiatric disorders. However, the gene regulatory programs defining basal ganglia cell-type identity and function remain poorly understood, limiting interpretation of disease-associated non-coding variants. Here, we present the first single-cell multiome atlas of histone modifications and transcriptomes across eight basal ganglia regions from neurotypical adult human donors. Joint profiling reveals cell-type-specific deployment of active and repressive *cis*-regulatory elements and gene regulatory networks, and suggests a combinatorial homeobox transcription factor code underlying cell identity. Integration with matched spatial transcriptomic MERFISH data uncovers regional heterogeneity of epigenomic landscapes. Comparative analysis between human and mouse medium spiny neurons uncovers conservation of core gene regulatory features. This atlas interprets non-coding risk variants of neuropsychiatric disorders and supports the development of a deep learning model to predict gene regulation and functional effects of disease-associated variants.

**HIGHLIGHTS:**

- Joint single-cell profiling of transcriptomes and three histone modifications across eight human basal ganglia regions characterizes active and repressive chromatin states at cell-type resolution.
- Cell-type-specific gene regulatory programs decode combinatorial homeobox TF grammar governing the identity and diversification of basal ganglia neurons.
- Intergrative analyses link noncoding neuropsychiatric risk variants to specific cell types, regulatory elements, and candidate target genes.
- A sequence-to-function deep-learning model predicts gene regulation from DNA sequence and prioritizes functional disease-associated variants.

## INTRODUCTION

The basal ganglia (BG) comprise a group of interconnected nuclei that orchestrate diverse neural circuits governing motor control, emotion, cognition, learning, and reward processing^1–4^. Dysfunction of BG circuits contributes to a broad spectrum of neurological and psychiatric disorders, including addiction^5,6^, Parkinson’s disease^7^, Huntington’s disease^8^, and schizophrenia^9^. The BG contain diverse neuronal and non-neuronal cell types, with particularly prominent contributions from GABAergic neurons^10,11^. Characterizing cell-type-specific features at the molecular and cellular levels is essential for understanding both BG function and the mechanisms underlying BG-related diseases. Single-cell transcriptomic studies have dissected the cell taxonomy of the BG in rodents^12–18^ and primates^19–25^, and have illuminated cell-type-specific gene expression signatures. However, the gene regulatory programs that establish and maintain BG cell identities remain poorly defined.

The distal *cis*-regulatory elements (CREs), together with the transcription factors (TFs) that bind to them comprise the gene regulatory networks that control cell-type-specific gene expression^26^. While tens of thousands of genetic variants have been associated with neurological and mental disorders^27^, the vast majority of these risk variants fall outside protein-coding regions^28,29^. These non-coding variants are hypothesized to contribute to disease pathogenesis by disrupting cell-type-specific CREs and altering target gene expression^30^; however, the regulatory mechanisms for most loci remain elusive due to a lack of comprehensive annotation of regulatory elements and their activity across human brain regions and cell types. Epigenomic profiling provides a vital bridge, allowing for the annotation of putative CREs, the elucidation of gene regulatory programs, and the functional interpretation of non-coding variants^28^.

Previously, candidate *cis*-regulatory elements (cCREs) in the human brain were identified using single-cell chromatin accessibility mapping (e.g., sci-ATAC-seq), providing a catalog of candidate promoters and enhancers for over a hundred brain cell types^31^. However, accessibility alone is insufficient to identify the repressive or poised regulatory elements. Incorporating histone modification profiles has the potential of overcoming this limitation, as different activity states of cCREs are characterized by distinct combination of histone marks: H3K27ac decorates active promoters and enhancers; H3K27me3 demarcates facultative heterochromatin and repressed states; and H3K9me3 marks constitutive heterochromatin and silenced states^32,33^. Previous studies have reported that histone modification dysregulation is a common theme in synaptic plasticity and cognition impairments associated with BG dysfunction^34^. Importantly, as these post-translational modifications are reversible, modulation of these epigenetic marks presents opportunities for therapeutic interventions^35,36^ aimed at resetting regulatory states to modify downstream molecular and behavioral phenotypes in diseases.

In this study, we present a comprehensive single-cell atlas of three histone modifications, including H3K27ac, H3K27me3, and H3K9me3, each jointly profiled with the transcriptome across eight BG regions from seven neurotypical adult human donors, using a recently developed single-cell multiomic approach known as Droplet Paired-Tag^37^. Integrative analysis of ∼600,000 nuclei resolved 61 cell groups and annotated chromatin states across 50% of the genome on average. All data are available for direct visualization and exploration in the BICAN Basal Ganglia Epigenome Portal (https://basalganglia.epigenomes.net)^38^.

This atlas enabled the annotation of 532,547 cCREs in total, with potential target genes defined for 356,380 distal cCREs. We also identified 179,396 repressive chromatin regions forming loops with target genes in a cell-type-specific manner. To map gene regulation in anatomical context, we generated a MERFISH-based spatial transcriptomic dataset and integrated it with the Droplet Paired-Tag dataset to reconstruct a high-resolution spatial epigenomic map that allowed us to investigate how gene expression patterns and epigenomic regulatory logic are spatially organized. Integration with mouse brain Paired-Tag data further enabled systematic analysis of evolutionary conservation and divergence in epigenome and gene regulation.

Analysis of cell-type-specific regulatory programs provides evidence supporting the hypothesis that combinatorial homeobox TFs code drives neuronal identity. Distinct regulatory programs are highlighted for D1 and D2 medium spiny neuron (MSN) subtypes, which exhibit highly similar transcriptome and epigenomic states relative to other BG cell types. We further utilized these regulatory networks to interpret likely causal variants of neuropsychiatric diseases in each cell type, and developed a deep-learning model to predict the functional impact of disease risk variants. Collectively, the single-cell atlas reported in this study enables the understanding of BG cell-type-specific gene regulatory programs, improves the interpretation of BG-relevant non-coding disease risk variants, and lays the groundwork for more effective investigations into the molecular underpinnings of neurological and psychiatric disorders.

## RESULTS

### A single-cell atlas of histone modifications and transcriptome for human basal ganglia

To comprehensively map the histone modification landscapes of the human BG, we performed dissections of eight anatomical BG subregions from seven neurotypical donors (5 male, 2 female; age range 29–67 years). These regions included head of caudate nucleus (CaH), body of caudate nucleus (CaB), tail of caudate nucleus (CaT), globus pallidus (GP) containing the external and internal segments of globus pallidus (GPe, GPi) and ventral pallidum (VeP), putamen (Pu), nucleus accumbens (NAC), subthalamic nucleus (STH), midbrain gray matter (MGM1) containing substantia nigra (SN), red nucleus (RN), and ventral tegmental region (VTR) (Figure 1A and S1.1A-C; Table S1.1; Methods). For each brain sample, we carried out Droplet Paired-Tag^37^ to simultaneously profile the transcriptome and the histone modification landscape at single-nucleus resolution. To capture the spatial context of individual cells, we performed MERFISH^39^ assays to monitor the expression of ∼1000 genes across the same BG regions at single molecular and sub-optical diffraction resolution in parallel (Figure 1A and S1.1A; Table S1.2). To increase the representation of neuronal diversity, we utilized fluorescence-activated nucleus sorting (FANS) to enrich the neuronal fraction to ∼50%. The nuclei pool was subsequently divided into three parallel reactions, each targeting a distinct histone mark, including H3K27ac, H3K27me3, or H3K9me3 (Figure 1B). A total of 793,276 nuclei passed the initial quality controls. After removing profiles that likely resulted from barcode collisions or doublets, 608,879 nuclei were retained: 96,051 from CaH, 102,705 from CaB, 37,702 from CaT, 57,829 from GP, 98,310 from Pu, 79,495 from NAC, 63,901 from STH, and 72,886 from MGM1 (Figure 1A).

**Figure 1.**
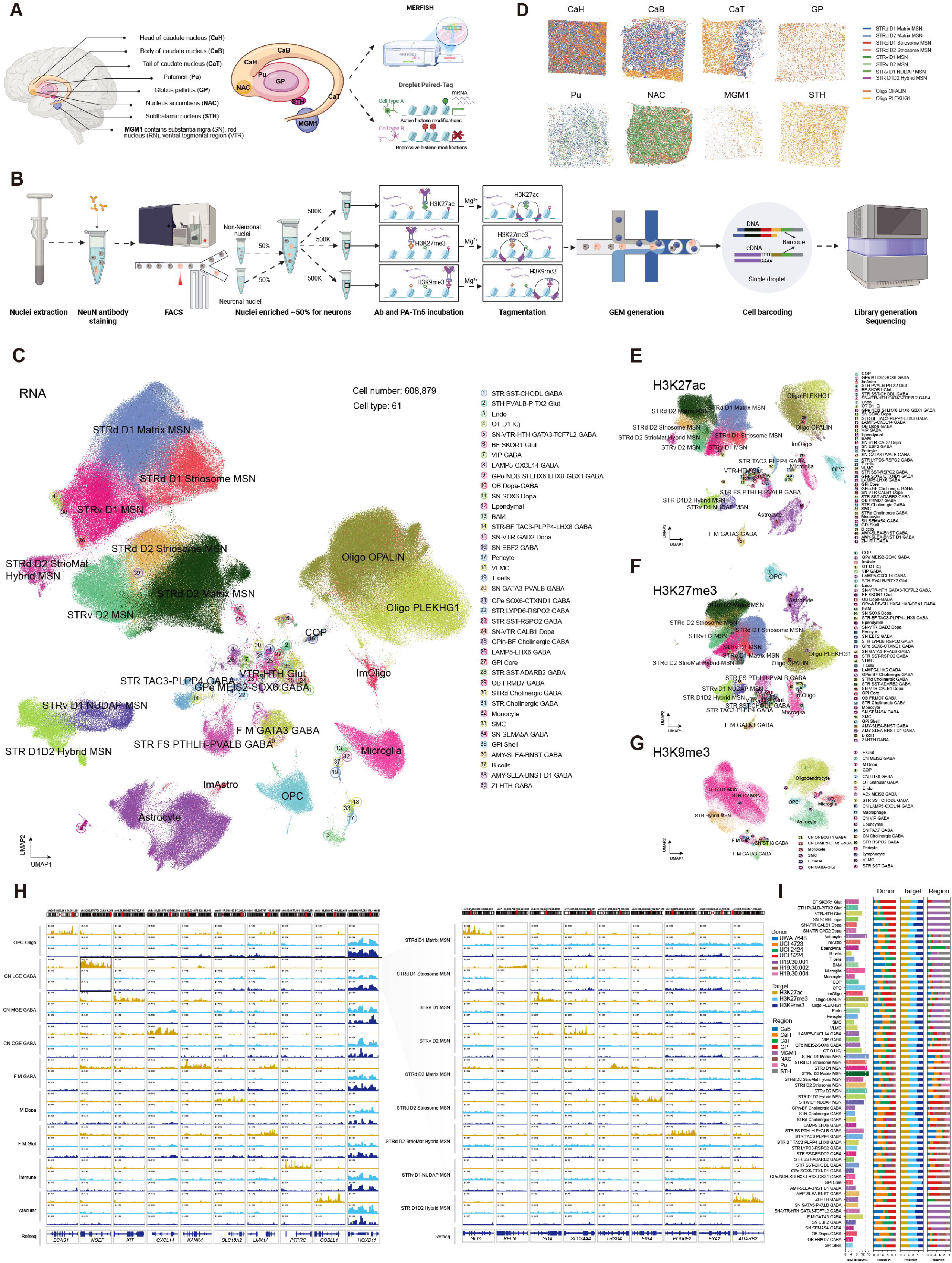
Single-cell joint profiling of histone modifications and transcriptome in the human basal ganglia. (A) Schematic of human basal ganglia dissections and multimodal profiling by Droplet Paired-Tag and MERFISH spatial transcriptomics. (B) Workflow of nuclei isolation and library preparation for Droplet Paired-Tag. (C) UMAP embedding and clustering analysis of transcriptome modality from the Droplet Paired-Tag datasets, colored by group types. (D) The spatial maps of cell groups in each basal ganglia region identified using MERFISH with the probe set targeting ∼1,000 genes. (E) UMAP embedding and clustering analysis of H3K27ac profiles from the Droplet Paired-Tag datasets, colored by group types defined by corresponding single-cell transcriptome. (F) UMAP embedding and clustering analysis of H3K27me3 profiles from the Droplet Paired-Tag datasets, colored by group types defined by corresponding single-cell transcriptome. (G) UMAP embedding and clustering analysis of H3K9me3 profiles from the Droplet Paired-Tag datasets, colored by subclass types defined by corresponding single-cell transcriptome. (H) Genome browser tracks showing aggregated histone modification profiles for each class and MSN group within CN LGE GABA class at selected marker gene loci that were used for cell cluster annotation. (I) Bar plots showing, from left to right, total number of nuclei, relative contribution of donors, three histone modification marks distribution, and brain region distribution for each of the 61 cell groups identified in the Droplet Paired-Tag dataset.

The Droplet Paired-Tag dataset generated in this study exhibits quality superior to previously published resources^37^. The transcriptomic libraries yielded a median number of 7,931 UMI and 3,131 genes per nucleus under 34.8-80.3% duplication rate (Figure S1.2A). The complexity of the DNA-dedicated libraries for histone modifications is higher than previous report^37^, with median fragment counts of 4,097 for H3K27ac, 6,005 for H3K27me3, and 3,751 for H3K9me3 (Figure S1.2B). Data reliability was further confirmed by reasonable fraction of reads in peaks scores (FRiP) (Figure S1.2C), enrichments of reads near transcription start sites (TSSs) (Figure S1.2D), and strong correlations across donors within the same region (Figure S1.3A-D).

We next applied MapMyCells^14^, a hierarchical correlation mapping algorithm, to 608,879 nuclei, to categorize each cell based on the Human and Mammalian Brain Atlas (HMBA) consensus basal ganglia taxonomy^11^ (Figure 1C and S1.4A). Nuclei were first assigned to four neighborhoods: neighborhood I enriched for glutamatergic (Glut, putatively excitatory) neurons (1.37%); neighborhood II enriched for non-neuronal cells (37.14%); neighborhood III enriched for GABAergic (GABA, putatively inhibitory) neurons (61.47%); and neighborhood IV enriched for Subpallium GABA-Glut (0.02%) (Figure S1.4B). Hierarchical mapping further assigned cells into 12 classes and 36 subclasses, including 2 subclasses of glutamatergic neurons, 1 subclass of dopaminergic neurons, 13 subclasses of non-neuronal cells, 16 subclasses of GABAergic neurons, 3 subclasses of MSNs, and 1 subclass of GABA-Glut (Figure S1.4B). Lastly, nuclei in each subclass were further classified into 1–3 unique groups, yielding 61 distinct groups in total (Figure S1.4B). To estimate the performance of MapMyCells, we calculated the overlap scores between our query nuclei annotation and the reference cell identity in major group-level clusters, which showed strong consistency (Figure S1.4C), indicating the accuracy of cell type annotation of Droplet Paired-tag dataset. Several group-level differences between our study and the HMBA basal ganglia consensus taxonomy^11^ were observed, reflecting the distinctions in tissue dissection boundaries and species scope. First, our data revealed an undefined GABAergic group under F M GATA3 GABA subclass, termed F M GATA3 GABA at group level, which was specific to MGM1. We speculated that this cell group originated from the RN, which was included in our MGM1 dissections but not in the HMBA dissections. Second, our data lacks SN GATA3-PAX8 GABA group, which is specific to marmoset and macaque, and not present in human in the HMBA consensus basal ganglia taxonomy for 3 primate species.

The single-cell spatial transcriptomic profiles were annotated using the Droplet Paired-Tag transcriptome as reference. After segmentation and removal of low-complexity cells, MERFISH profiles showed high reproducibility across replicates from two different labs and concordance with bulk RNA-seq (Figure S1.5A-D). Confident cell labels were assigned to 765,076 MERFISH cells with more than 100 transcripts per cell using canonical correlation analysis (CCA) integration^40^. The spatial distributions of BG cell groups differed across BG subregions. For example, MSNs and oligodendrocytes formed clustered compartments within the caudate nucleus (Figure 1D). The relative proportions were quantified at both subclass and group levels and showed various cellular compositions across BG regions (Figure S1.6A-B).

We also performed independent clustering for each histone modification atlas and transferred cell-type identities from the paired transcriptomic data. While all 36 subclasses were recovered across all three marks, the resolution of specific cell groups varied by histone modification types. H3K27ac and H3K27me3 successfully resolved all 61 groups (Figure 1E, F), and H3K9me3 profiles recovered 60 groups (Figure 1G). Specifically, STR D1 and STR D2 MSNs separated clearly in H3K27ac and H3K27me3 UMAPs at both subclass and group levels, but were intermixed in H3K9me3 UMAP (Figure 1E-G), likely reflecting a common cellular lineage of the two cell subclasses. Histone modification signals at marker gene loci recapitulated cell-type-specific expression patterns. Visualization of the three histone marks on canonical marker genes of BG classes and MSN groups showed that H3K27ac enrichment at TSS and gene body was tightly coupled with cell-type-specific high expression (Figure 1H and S1.7A). In contrast, these TSSs and gene bodies were marked by weak H3K27ac signals or repressive marks (H3K27me3/H3K9me3) in cell types where the gene expression were silenced (Figure 1H). We also found strong repressive H3K27me3 and H3K9me3 signals on the *HOXD11* gene across all BG cell types, in accordance with the silenced state of this gene (Figure 1H).

We further examined cell number, donor contribution, histone mark composition and region distribution for each of the 61 cell groups (Figure 1G and S1.7B-D). Most non-neuronal cell types were evenly distributed among different BG subregions (Figure 1G and S1.8), whereas some neuronal types displayed strong regional specificity (Figure 1G and S1.8). Specifically, the six groups under Glut Sero Dopa neighborhood had highly enriched subregional distributions within STH and MGM1 (Figure 1G and S1.8), while the nine MSN groups were preferentially distributed across striatal regions (Figure 1G and S1.8).

In summary, we generated the first high-quality single-cell atlas jointly profiling histone modifications and the transcriptome across eight BG subregions in neurotypical adult humans, resolving 61 distinct BG cell types.

### Annotating the gene regulatory elements and chromatin states in basal ganglia cell types

To systematically characterize chromatin regions modified by distinct histone marks, we annotated each histone mark-modified DNA sequence with standard genomic features. In the basal ganglia, the single-cell H3K27ac, H3K27me3, and H3K9me3 profiles spanned 34.8%, 48.1%, and 43.9% of the genome, respectively (Figure 2A). Collectively, these three histone marks covered 61.3% of the genome. By contrast, single-cell ATAC-seq profiles for the whole human brain covered 8.7% of the genome^31^, and bulk human brain ATAC-seq datasets from ENCODE covered 9.3%^32^. Therefore, chromatin-state annotation using histone modification profiles has the potential for annotating a substantially broader genomic landscape. The proportions of genomic features, including gene related features such as promoters/TSSs, untranslated regions (UTRs), exons, introns, intergenic regions and transposable elements (TEs), varied across histone marks. Consistent with its roles in active regulatory chromatin, H3K27ac was preferentially enriched at promoters, TSS-proximal regions, and gene bodies (exons and introns), in a manner that is positively correlated with gene expression (Figure 2B). More than half of H3K27me3- and H3K9me3-marked regions overlapped TE sequences, including long interspersed nuclear elements (LINEs), short interspersed nuclear elements (SINEs), long terminal repeats (LTRs), simple repeats and satellite sequences (Figure 2B). Relative to H3K27ac, H3K27me3- and H3K9me3-marked regions showed higher proportions in LINEs, LTRs, and satellite sequences, but lower proportions in SINEs (Figure 2C).

**Figure 2.**
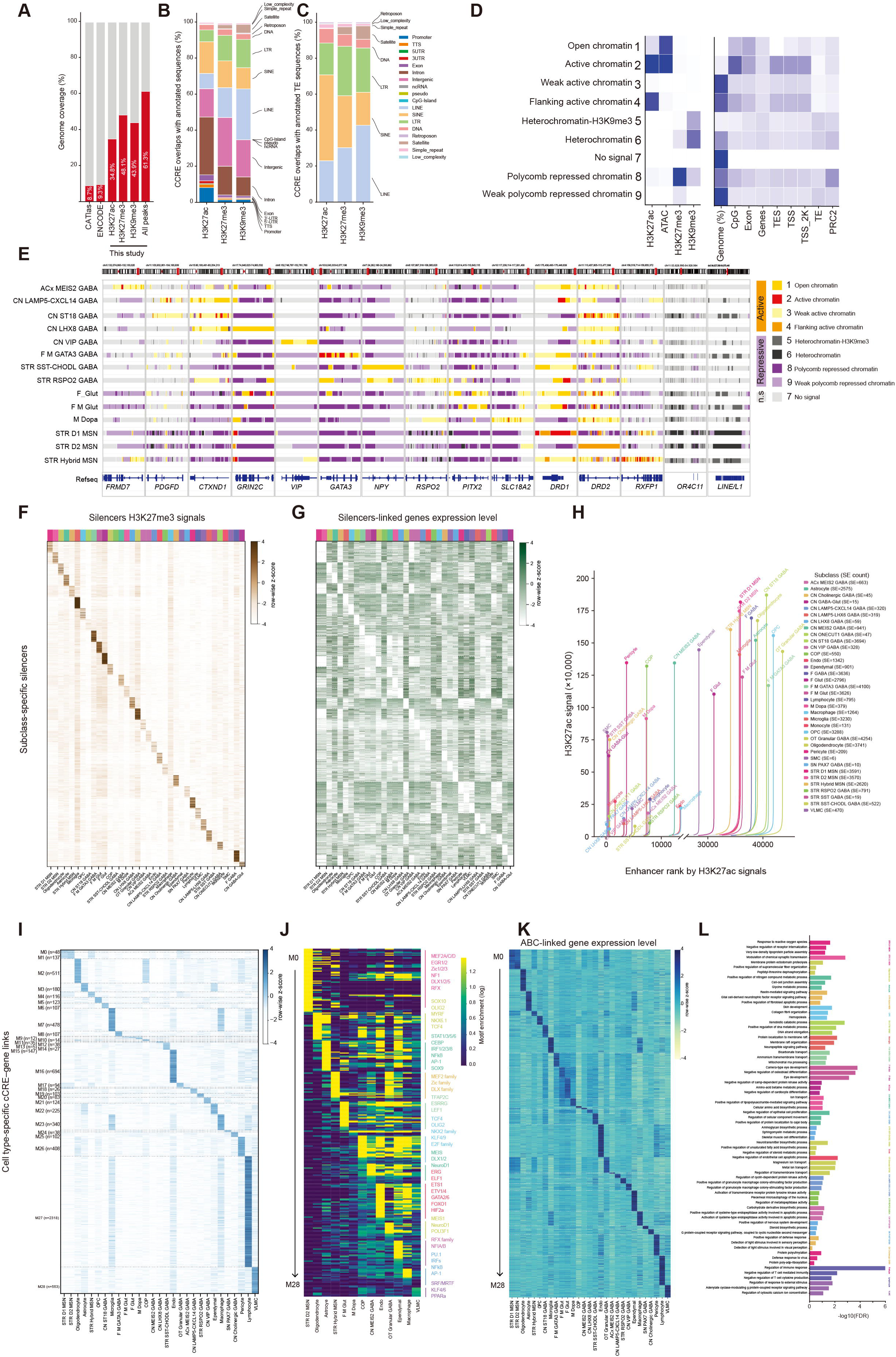
Droplet Paired-Tag defines chromatin states and identifies cell-type-specific gene regulatory elements. (A) Stacked bar plot showing the percentage of genome (red) covered by regulatory elements identified from previous single-cell ATAC-seq dataset in CATlas, bulk ATAC-seq datasets from ENCODE project, single-cell histone modification profiling datasets for three marks generated in this study, and the combined signal from all three marks in this study. (B) Bar plot showing the fraction of three histone modification mark profiles overlapping different classes of standard genomic features. LTR, long terminal repeat; TTS, transcription termination site; UTR, untranslated region. (C) Bar plot showing the fraction of three histone modification mark profiles overlapping different classes of TEs in the human genome. (D) Left, emission probabilities for three histone modifications and open accessibility in 9 ChromHMM states. Right, genomic feature enrichments for each ChromHMM state. (E) Chromatin state landscapes at subclass marker gene loci for major neuronal subclasses. (F) The H3K27me3 modification levels at 36 subclass-specific silencers modules identified from S-P loops. (G) The RNA expression levels of silencer linked genes within S-P loops across 36 subclasses. (H) Super enhancers identified using H3K27ac profiles across different subclasses. (I) Heatmap showing H3K27ac levels at putative enhancers linked to genes by the ABC model. (J) TF motif enrichment in ABC model-identified putative enhancers for distinct modules. (K) Heatmap showing expression levels of genes linked to putative enhancers by ABC model in each subclass. (L) Biological pathways enrichment in genes linked to putative enhancers by ABC model in each subclass.

Next, we inferred chromatin states using ChromHMM^41^ by jointly modeling three histone modifications together with chromatin accessibility data from human basal ganglia reported in the HMBA companion manuscript^11^. After evaluating models across multiple parameterizations, we selected a 9-state model that exhibited near-perfect concordance between two pseudo replicates and was broadly consistent with previously reported chromatin-state annotations (Figure S2.1A-E). State identities were defined by their emission profiles and assigned descriptive labels based on similarity to established chromatin signatures. The model resolved four active chromatin states: open chromatin with high level accessibility and weak H3K27ac, active chromatin with concomitantly high accessibility and H3K27ac, flanking active chromatin with strong H3K27ac but low accessibility, and weak active chromatin with only low H3K27ac (Figure 2D). The model also resolved four repressive chromatin states: H3K9me3-marked heterochromatin, mixed heterochromatin with high H3K9me3 and weak H3K27me3, H3K27me3-marked polycomb-repressed chromatin, and weak polycomb repressed chromatin with only low H3K27me3 (Figure 2D). An additional unknown state showed no detectable signal across any of the four profiles (Figure 2D). These chromatin states annotated 44.6-60.14% of the genome across BG cell types (Table S2.1). These chromatin-state maps enabled visualization of multiple functional predictions. Marker genes for each BG subclass adopted active states in the corresponding subclass but repressive states in other subclasses (Figure 2E). Conversely, genes with little or no expression in the basal ganglia were consistently annotated as repressive states across subclasses, including *OR4C11* and the gene-poor LINE1 elements (Figure 2E).

Regulatory elements, such as promoters, enhancers and silencers, bind TFs to control cell-type-specific transcriptional programs. To extract candidate regulatory elements from histone modification profiles and infer gene regulatory programs in each cell type, we first identified subclass-resolved cCREs from H3K27ac profiles, including both promoters and enhancers. We aggregated H3K27ac profiles from nuclei within each subclass into two pseudo-replicates, called peaks with MACS2 following the ENCODE ChIP-seq analysis pipeline, and retained only peaks reproducibly detected in both replicates. In total, we identified 532,547 H3K27ac peaks across 36 subclasses, 64.38% of which overlapped with HMBA basal ganglia snATAC-seq peaks^11^, including 311,977 shared by more than one subclasses and 220,570 that were unique to a single subclass. The cell-type-specific cCREs displayed unique H3K27ac patterns in most subclasses, whereas MSNs exhibited highly similar patterns between STR D1 and STR D2 subtypes. We next examined chromatin-state annotations for these H3K27ac cCREs across BG subclasses. State assignments were strongly cell-type dependent (Figure S2.2A). Consistent with subclass-restricted regulatory programs, cCREs were annotated as active states in the corresponding cell type, whereas the same elements were frequently assigned repressive states in other cell types where they were inactive (Figure S2.2B).

The cCRE catalog defined from H3K27ac peaks provided a broad set of cCREs. To nominate a higher-confidence set of functional enhancers, we leveraged cell-type-resolved chromatin loops from snm3C-seq (Ding *et al.*, co-submit), which shared the same donors and dissections with Droplet Paired-Tag, to identify enhancer–promoter (E-P) loops, in which one anchor overlapped a promoter and the other was marked by H3K27ac^42^ (Table S2.2). Genes whose promoters were located in these loops showed elevated expression in the corresponding cell types, whereas the paired distal anchors exhibited strong H3K27ac signals, supporting their roles as active enhancers (Figure S2.2C-D). These loop-defined, cell-type-specific enhancers displayed distinct TF binding motif signatures, including PKNOX, EGR1/2, and RFX1/2 in neuronal subclasses, lineage regulators such as MYRF in oligodendrocytes, and immune-associated factors PU.1 and ISRE in microglia and immune populations (Figure S2.2E). Consistently, gene ontology (GO) enrichment analysis of enhancer-linked target genes by loops was also cell-type coherent, with neuronal subclasses enriched for dendrite, axon and synaptic signaling terms, and immune populations enriched for immune and metabolic regulatory pathways (Figure S2.2F). Notably, the enhancer–promoter loops remained largely consistent between STR D1 and STR D2 MSNs.

Similarly, we identified silencer–promoter (S-P) loops using H3K27me3 profiles, requiring one anchor to overlap a promoter and the other to be marked by H3K27me3^42^ (Table S2.3). Target genes whose promoters participated in these loops showed reduced expression in the corresponding cell types, concordant with strong H3K27me3 signals at the paired distal silencer anchors (Figure 2F-G). Silencer landscapes were also subclass-specific and revealed distinct TF binding motif enrichments between STR D1 and STR D2 MSNs: STR D1 MSN silencers were enriched for regulatory factors of GABAergic lineages (GSX2 and LHX6) and glutamatergic neurogenesis programs (NEUROG2); STR D2 MSN silencers were enriched for nuclear receptors, immune and inflammatory regulators (RORα/γ, THR, PU.1, ETS1/ELF, NFκB-p65, and IRF). Silencers in F M Glut were enriched for ETS, GABPA and FOX motifs together with NEUROD/E-box elements, functioning in regulation of metabolic state and differentiation programs (Figure S2.2G). GO analysis of silencer-linked target genes highlighted enrichment for metabolic processes and broad receptor and enzyme activity terms (Figure S2.2H).

To identify super-enhancers (SE), we applied the ROSE algorithm^43,44^ to H3K27ac profiles and assigned putative target genes based on promoter proximity. We identified a total of 54,847 SEs from 36 subclasses (Figure 2H; Table S2.4). We detected broadly shared SEs at housekeeping gene loci, including *ACTB*, *RPLP0*, and *EEF2*; and cell-type-specific SEs, such as a MSN-enriched SE at *FOXP1* and a STR D1 MSN–specific SE at *DRD1*, the key marker gene distinguishing STR D1 MSNs from STR D2 MSNs (Figure S2.3A). GO analysis of SE-linked target genes recapitulated expected cell biology: neuronal SEs preferentially associated with ion transport, membrane depolarization/action, and synaptic transmission terms, whereas oligodendrocyte SEs were enriched for myelination-related pathways, and other non-neuronal subclasses showed distinct chemotaxis and cell signaling structural functions (Figure S2.3B).

To further resolve cell-type-specific transcriptional regulatory programs, we integrated subclass/group-aggregated H3K27ac, transcriptome, HMBA chromatin accessibility^11^, and chromatin contact profiles from snm3C-seq (Ding *et al.*, co-submit) into the activity-by-contact (ABC) framework^45,46^ to infer enhancer–promoter links for each cell type (Table S2.5). Each gene was assigned an average of 28.81 putative enhancers, and each enhancer candidate was linked to 2.41 genes on average. This strategy linked 7,547 cell-type-specific enhancers to 7,949 putative target genes. These enhancers exhibited strong H3K27ac enrichment in the corresponding cell type (Figure 2I), except for STR D1 and STR D2 MSNs. TF binding motif enrichment in cell-type-specific enhancers nominated key regulators of cell identity, including key regulators of GABAergic neurons DLX1/2 in CN MEIS2 GABA subgroup, OLIG2 in oligodendrocytes and OPCs, and SOX9 in astrocytes (Figure 2J). The 7,949 target genes showed corresponding cell-type-specific expression (Figure 2K). These target genes were enriched for processes coherent with cellular function and identity, including modulation of chemical synaptic transmission enriched in STR D2 MSN, neurotransmitter biosynthetic process enriched in STR SST-CHODL GABA neurons, and cell-cell junction assembly enriched in astrocytes (Figure 2L).

Together, these analyses expand the functional epigenomic landscape of the human basal ganglia and provide a comprehensive catalog of cell-type-specific regulatory elements and chromatin states.

### Cell-type-specific gene regulatory programs in basal ganglia

To delineate cell-type-specific gene expression and histone modification features in basal ganglia, we first identified variably modified genomic regions and differentially expressed genes in each cell type. Across major neuronal and non-neuronal subclasses, extensive cell-type-specific features (FDR< 0.10) in all four modalities were identified, spanning hundreds to thousands of genes and genomic regions per subclass (Figure 3A). GO enrichment analysis of differentially expressed genes showed that STR D1 and STR D2 MSNs shared highly similar biological process enrichments. Relative to other subclasses, MSNs showed significant enrichment for terms related to striatum development, together with relative depletion of programs associated with dopaminergic neuron differentiation, mesenchymal cell proliferation, neuroendocrine cell differentiation (Figure S3.1).

**Figure 3.**
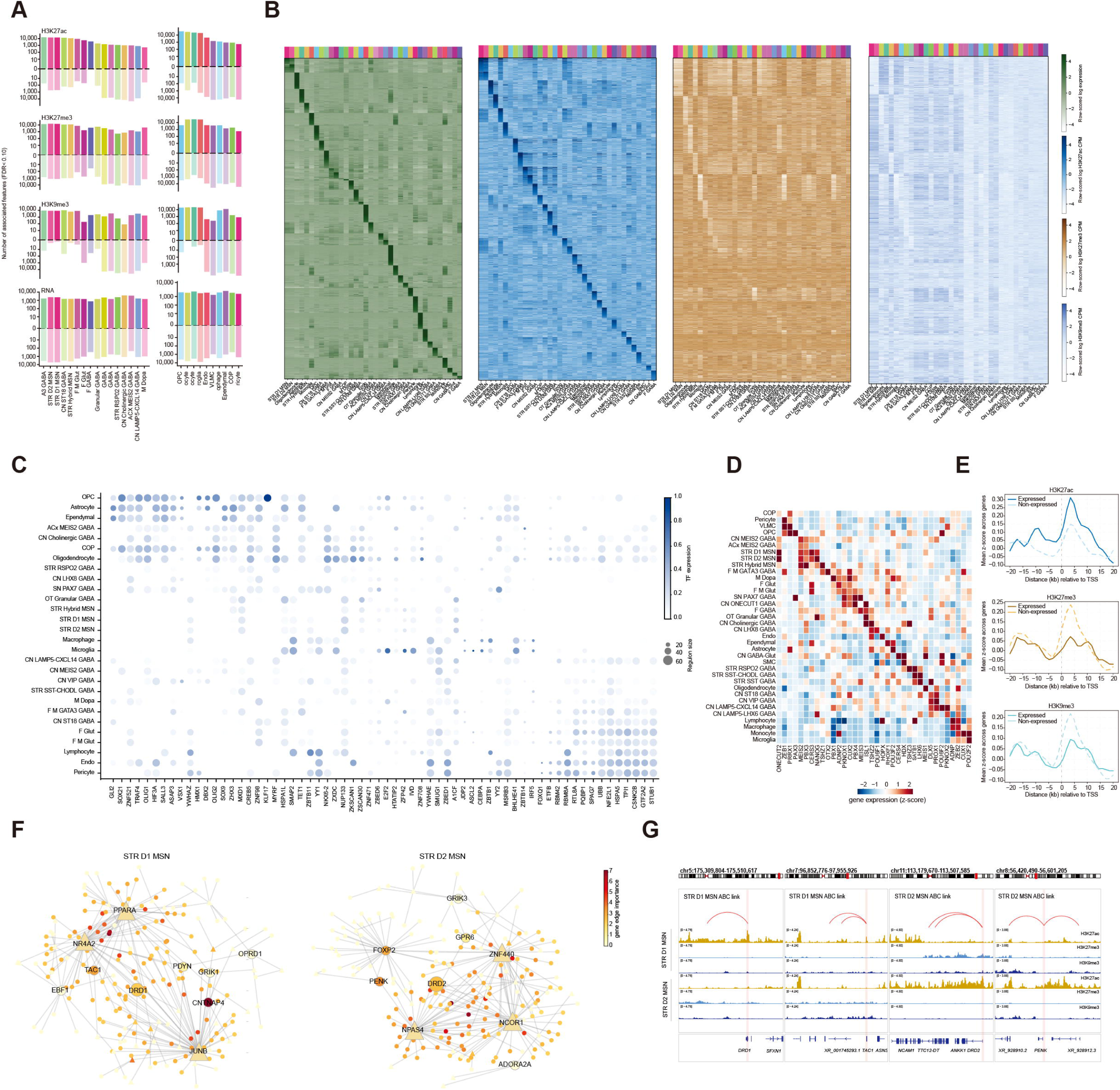
Cell type-specific epigenetic changes and gene regulatory networks. (A) Number of genes and genomic regions in each subclass for each modality with significant up and down changes. Features were quality control filtered by FDR<0.1. (B) Heatmaps showing expression levels of cell type-specific upregulated genes and three histone modification levels of their top putative enhancers from ABC model across different subclasses. (C) Clustermap of eRegulons from TF regulatory networks inferred by SCENIC+ and overlapped with enhancers inferred from the ABC model for each subclass of cells in the basal ganglia. Dot size indicates the number of genes within each TF eRegulon in each subclass, and dot color indicates TF expression levels in each subclass. (D) Heatmap showing the combinatorial expression pattern of selected homeobox transcription factor genes across different subclasses. (E) Average histone modification levels around the TSS of expressed versus non-expressed homeobox transcription factor genes. (F) Network plots illustrating representative GRNs with altered activity in STR D1 and STR D2 MSNs. Triangles represent TFs, and circles represent target genes. Both TFs and target genes are colored according to SCENIC+ derived regulatory edge importance scores. (G) Examples of gene regulatory networks with STR D1 MSN-specific and STR D2 MSN-specific putative enhancer and target gene links from ABC model and histone modification tracks in STR D1 and STR D2 MSNs.

To determine how histone modifications shape cell-type-specific gene expression, we quantified H3K27ac, H3K27me3, and H3K9me3 signals around TSSs (± 2 kb) of highly expressed subclass marker genes. Across cell types, highly expressed genes exhibited strong promoter-proximal H3K27ac levels accompanied by depletion of the repressive H3K27me3 and H3K9me3 modifications (Figure 3.2A). This close coupling between histone modification profiles and transcript levels suggested that the local epigenomic environment served as an indicator of transcriptional activity. We next examined H3K27ac levels at the top three ABC model-predicted cCREs linked to each subclass marker gene and observed higher modified levels in the corresponding subclass than in others (Figure 3B and S3.2B). We also found that a subset of cCREs linked to these marker genes by the ABC model exhibited elevated repressive histone marks in non-expressing subclasses and showed cell-type-specific enrichment of TF binding motifs (Figure 3B and S3.2C). Together, these results indicate that subclass-specific gene expression is associated with coordinated histone modification states at both promoters and distal enhancers.

Furthermore, we applied SCENIC+^47^ to the H3K27ac Droplet Paired-Tag dataset to infer gene regulatory networks (GRNs), including enhancer–TF–target gene relationships, across BG cell types (Table S3.1). SCENIC+ identified candidate enhancers from the single-cell H3K27ac dataset, enriched TF-binding motifs occurring within candidate enhancers, and linked TFs to candidate enhancers and target genes. We identified a subset of enhancer-driven regulons (eRegulons), each consisting of a TF, its associated regulatory enhancers, and target genes. To increase confidence, SCENIC+-defined enhancers were further intersected with enhancer–gene links inferred by the ABC model. We then quantified eRegulon activity across BG subclasses using TF expression and the number of target genes per eRegulon (Figure 3C). This integrative GRN analysis revealed that eRegulon TFs had different effect size and specificity across different cell types (Figure 3C). We identified BG subclass-specific eRegulons and highlighted broad enrichment for homeobox TFs (40/152), indicating they are the central components of human BG GRNs (Table S3.2).

The homeobox TF eRegulons displayed striking cell-type-specific patterns in both expression levels and eRegulon sizes (Figure S3.2D). Homeobox TFs are known to play a key role in neural development^48–50^. Previous studies have reported that each individual neuron class in the nematode Caenorhabditis elegans can be defined by the expression of a unique combination of homeobox genes^51–53^. To test whether a similar Homeobox TF grammar operates in the human basal ganglia, we first examined homeobox TF expression across BG subclasses and observed a subset of homeobox TFs with strong cell-type specificity and functional enrichment in neuronal differentiation and axial patterning (Figure 3D and S3.2E). As high-level regulators of developmental lineages, homeobox genes are subject to strict epigenetic regulation and have been reported to be controlled by PRC2 during development^54^. Profiling histone modifications around TSSs revealed that expressed homeobox TF genes displayed TSS-proximal H3K27ac enrichment, whereas non-expressed homeobox TF genes exhibited promoter-associated repression with elevated H3K27me3 and H3K9me3 (Figure 3E). These patterns support a model that homeobox TF gene expression is tightly controlled by chromatin states in a cell-type-restricted manner.

We next tested the hypothesis that homeobox TFs combinatorially achieve cell-type-specific downstream regulation beyond differences in expression. While the chromatin accessibility of homeobox TF binding motifs didn’t show strong cell-type specificity (Figure S3.2F), co-occurrence of other TFs, evident by specific combinations of homeobox TF motifs and co-TF motifs^55^, exhibited cell-type specificity (Figure S3.3A). For example, PBX and MEIS established co-factors of HOX family TFs, and showed selective co-accessibility with distinct HOX paralogs in a cell-type-specific manner. PBX/MEIS co-accessibility was preferentially associated with HOXA in CN LAMP5–LHX6 GABA neurons, HOXB in F M GATA3 GABA neurons, HOXC in CN ONECUT1 GABA neurons, and HOXD in CN cholinergic GABA neurons (Figure S3.3B). Together, these results suggest that neuronal identity is maintained through both histone modification-mediated control of homeobox TF expression and cell-type-specific accessibility of co-TF motifs that enable distinct combinatorial binding programs and downstream gene regulation.

MSN is the predominant neuronal type of the basal ganglia. In the classical model of BG circuitry, MSNs are broadly partitioned into two subtypes with opposing roles: STR D1 MSNs preferentially express the dopamine D1 receptor and promote movement via the direct pathway, whereas STR D2 MSNs express the dopamine D2 receptor and suppress movement via the indirect pathway^56,57^. Despite these well-established functional distinctions, the transcriptomic and active epigenomic regulatory features of STR D1 and STR D2 MSNs have modest differences in global analysis across all the BG subclasses, indicating a highly similar state between these two MSN subtypes. We therefore focused on utilizing inferred GRNs to illuminate regulatory programs that underlie distinct functions. We first identified differentially expressed genes (DEGs) between STR D1 and STR D2 MSNs and performed GO enrichment analysis on upregulated gene sets in each subclass (Figure S3.4A-B). Genes upregulated in STR D1 MSNs were enriched for terms related to interneuron migration and addiction-associated processes, whereas genes upregulated in STR D2 MSNs were preferentially associated with G protein–coupled receptor (GPCR) signaling pathways and locomotory behavior (Figure S3.4B). The chromatin state transitions between STR D1 and STR D2 MSNs indicated epigenomic regulations behind the observed DEGs (Figure S3.4C). The GRNs inferred for STR D1 and STR D2 MSNs shared many key TFs and target genes (Figure S3.4D). Focusing on eRegulons centered on the dopamine receptor D1 (*DRD1*) gene in STR D1 MSNs and the dopamine receptor D2 (*DRD2*) gene in STR D2 MSNs, we identified NR4A2, PPARA, and JUNB as prominent regulators associated with *DRD1*, whereas ZNF440, NPAS4, and NCOR1 were linked to *DRD2* (Figure 3F). Beyond *DRD1* and *DRD2*, tachykinin precursor 1 (*TAC1*) and proenkephalin (*PENK*) were also among the top DEGs between STR D1 and STR D2 MSNs and exhibited high edge importance within the GRNs (Figure 3F and S3.4A). We checked ABC model-predicted enhancers linked to these four marker genes and found concordant regulatory signatures: elevated H3K27ac at both promoters and linked enhancers in the subtype with higher expression, together with increased H3K27me3 and H3K9me3 in the lower-expressing subtype (Figure 3G). Homeobox TFs also showed subtype-associated differences in expression and histone modifications between STR D1 and STR D2 MSNs (Figure S3.4E). For instance, the *IRX6* gene showed higher expression and promoter-proximal H3K27ac in STR D2 MSNs (Figure S3.4E-F). Collectively, these results illustrated that GRNs derived from our multimodal epigenomic atlas provided a mechanistic framework for interpreting cell-type-specific transcriptional programs in the basal ganglia.

In summary, cell-type–resolved regulatory programs inferred from Droplet Paired-Tag dataset uncovered a combinatorial Homeobox TF code that specifies human neuronal cell-type identity and enabled mechanistic interpretation of cell-type-specific gene expression signatures.

### Spatial axes and developmental trajectories in basal ganglia

Heterogeneity within shared cell types across anatomical regions has been reported in human basal ganglia at the levels of DNA methylation and chromatin architecture^58^. Our Droplet Paired-Tag dataset provides an opportunity to further interpret the transcription and epigenetic regulatory mechanisms underlying BG spatial heterogeneity (Table S4.1). Group-level annotation of MSN subclasses incorporated regional distribution information: STRv MSNs labeled ventral MSNs mainly from the ventral striatum region (NAC) and STRd MSNs represent dorsal MSNs mainly from dorsal striatum regions (CaB and Pu). UMAP embeddings colored by regional labels revealed that groups within STR D1 MSN, STR D2 MSN, and STR Hybrid MSN subclasses exhibited regional biases along both the ventral–dorsal (V–D) and anterior–posterior (A–P) axes (Figure 4A-B). For groups distributed across multiple regions, cells from the same anatomical region clustered together, indicating shared region-associated molecular features (Figure 4A-B). We observed MSN group abundance shifted across regions: STRv D1 and D2 MSNs, STRv D1 NUDAP MSNs predominated in NAC, whereas STRd D1 and D2 Matrix MSNs, STRd D1 and D2 Striosome MSNs were enriched in CaB and Pu (Figure 4C). Along the A–P axis of caudate nucleus, STRd D1 Matrix MSNs increased from head (CaH) to tail (CaT), whereas STRd D1 and D2 Striosome MSNs decreased (Figure 4C).

**Figure 4.**
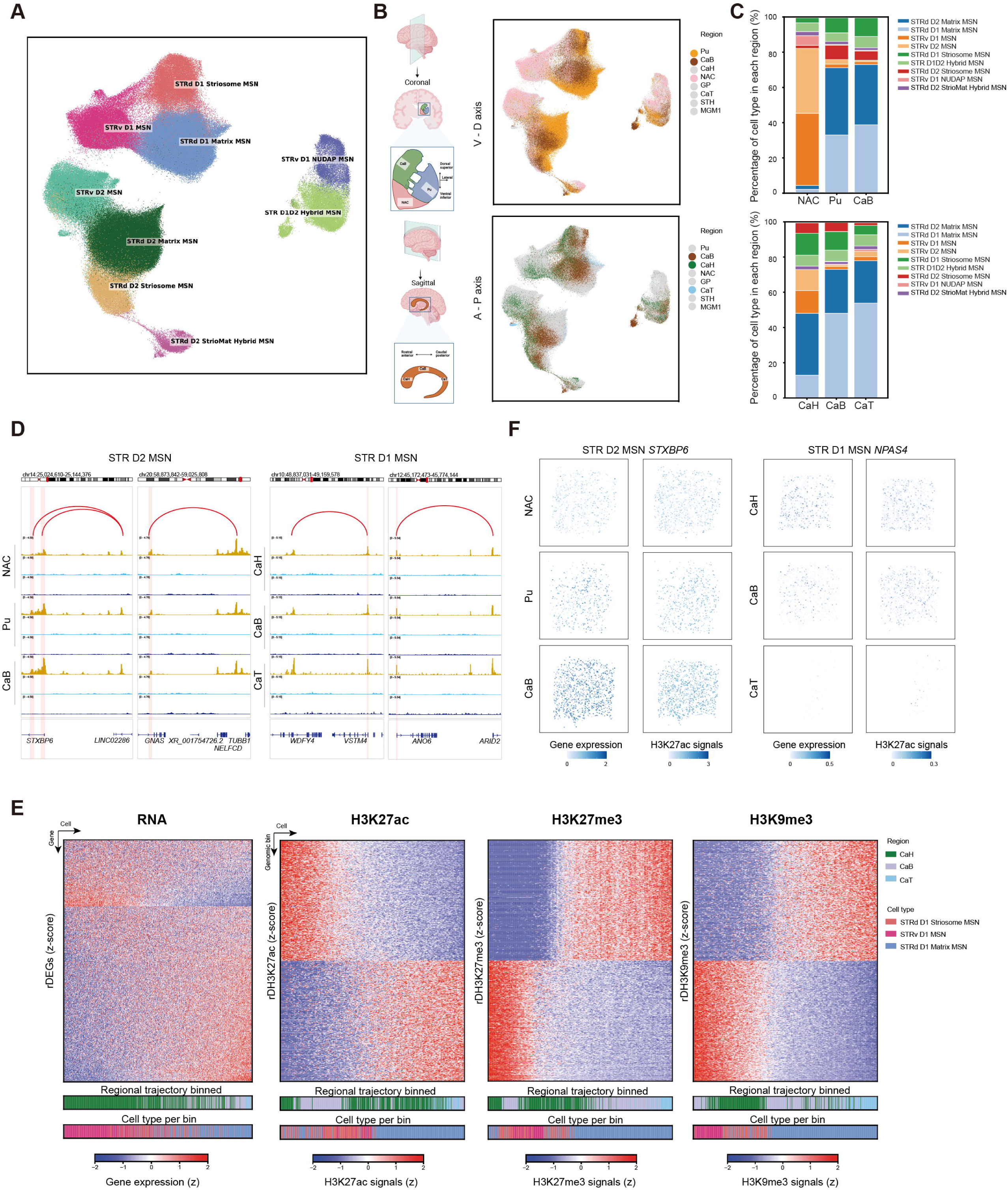
Epigenomic gradient changes along basal ganglia regional axes. (A) UMAP plot showing Droplet Paired-Tag transcriptome profiles of nine MSN groups. (B) UMAP plot showing MSNs profiles colored by brain region labels, highlighting regions along V–D axis (top) and A–P axis (bottom). (C) Stacked bars showing regional MSN group composition for each region along anatomical axes. (D) Genome browser tracks showing histone modification gradient changes along the V-D and A-P axis in both genes and their putative enhancers identified by the ABC model. (E) A–P axis gradients in expression of genes and modification–associated signal intensities of H3K27ac, H3K27me3 and H3K9me3 peaks for groups within STR D1 MSNs. (F) Droplet Paired-Tag gene expression and H3K27ac modification patterns of representative genes mapped to their predicted spatial location by integrative analysis with MERFISH. Left, the *STXBP6* gene shows the spatial gradient pattern among the V-D axis in STR D2 MSNs. Right: the *NPAS4* gene shows the spatial gradient pattern among the A-P axis in STR D1 MSNs. The left columns display the RNA expression from Droplet Paired-Tag. The right columns show the gene-body H3K27me3 signal density.

Previous studies have noted functional and connectivity distinctions between dorsal (CaB and Pu) and ventral (NAC) regions of striatum^1,58^, as well as anterior (CaH and CaB) and posterior (CaT) parts of the caudate nucleus^59,60^. To investigate the epigenetic foundations of these axial specializations, we analyzed gradient changes in transcriptome and histone modification landscapes in MSNs along anatomical V-D and A-P axes across different BG regions. Regional axes for each MSN subclass were constructed by ordering single cells along continuous transcriptional and epigenetic gradients. In STR D2 MSNs, this ordering revealed gene modules that progressively increased and decreased along a trajectory with ventral-to-dorsal organization from NAC to Pu and CaB (Figure S4.1A; Table S4.2). In parallel, we identified subsets of genomic bins with increased and decreased histone modification gradient changes along the trajectories showing V-D axial trends (Figure S4.1A; Table S4.2). For instance, moving from NAC through Pu to CaB, *GNAS* and *SEMA3D* genes decreased in expression and H3K27ac modification, whereas *STXBP6* and *NRG3* genes increased in both modalities (Figure 4D and S4.1B). Similarly, we also captured gene expression and epigenetic alterations in the STR D1 MSN subclass along the A-P axis of the caudate nucleus. (Figure 4E; Table S4.3). From CaH through CaB to CaT, *CDH10* and *NPAS4* genes showed decreasing expression and H3K27ac signals, while the *VSTM4* and *ANO6* genes increased in both modalities (Figure 4D and S4.1B). Consistent with ABC model-linked regulatory networks, *STXBP6*, *GNAS*, *VSTM4*, and *ANO6* genes also exhibited matched H3K27ac gradients at putative enhancers linked to their promoters. Systematic examination of regionally differential epigenetic features in MSNs identified 16,042 differentially histone-modified regions (DHRs) along the V–D axis in STR D1 MSNs, and 12,806 DHRs along the A–P axis in STR D2 MSNs. A considerable amount (2,183 and 4,489) of DHRs were linked to regional DEGs, indicating potential region-specific regulatory mechanisms based on histone modification, possibly underlying functional diversities. Specifically, by encoding a neuronal SNARE protein involved in the regulation of synaptic exocytosis, *STXBP6*^61^ can be a potential regulatory gene mediating the translation of regional epigenetic gradients into spatial functional distinctions. Together, these analyses provide an epigenetic framework for axial specialization and refine regional distinctions within shared BG cell types.

To increase the resolution of the spatial axes analysis, we leveraged the MERFISH dataset, which profiled the expression levels of ∼1,000 genes in brain slices from eight BG regions. We performed cross-modality integration between the Droplet Paired-Tag and MERFISH datasets using transcriptome as the anchor, which enabled imputation of spatial coordinates for 552,549 Droplet Paired-Tag nuclei. The spatial maps of the Droplet Paired-Tag closely matched the fine anatomical structures of MERFISH images. For example, the striosome MSNs displayed punctate distribution in CaH and CaB on Droplet Paired-Tag spatial maps, which reproduced the compartmental patterns at the group level on MERFISH images (Figure S4.1C). The gradient changes in transcriptome and histone modifications were also shown on the *STXBP6* gene in Droplet Paired-Tag nuclei spatial maps, which exhibited a ventral–to-dorsal increased expression in STR D2 MSNs, together with elevated H3K27ac signals at its promoter and gene body (Figure 4F). Along the A-P axis, *NPAS4* showed a spatial gradient decrease in expression in STR D1 MSNs from CaH to CaT, accompanied by reduced H3K27ac signals at the gene locus (Figure 4F). These results revealed a clear spatial pattern in histone modification that aligned with the spatial transcriptome, indicating that epigenetic regulation contributed to precise spatial specialization of the basal ganglia. By integrating Droplet Paired-Tag with MERFISH, we constructed spatially informed histone modification landscapes, enabling single-cell, spatial-resolved interpretation of epigenetic regulatory networks across BG structures.

In addition to the spatial gradients in transcriptome and histone modifications across regional axes, the developmental gradients associated with oligodendrocyte lineage were also observed within the basal ganglia. Pseudotime analysis based on transcriptomes reconstructed a trajectory from OPCs to oligodendrocytes (Figure S4.2A-B). In addition to genes exhibiting gradient up- and down-expression, subsets of genomic bins exhibited gradient increases or decreases in three histone marks along the OPC differentiation trajectory (Figure S4.2C). Consistent with prior reports that PRC2-mediated H3K27me3 is required for oligodendrocyte differentiation and myelination^62,63^, genome-browser tracks revealed that genes maintaining OPC undifferentiated and blocking oligodendrocyte differentiation gained H3K27me3 signals in oligodendrocytes compared with OPCs and COPs, including *HES5*, *BMP4*, and *ID2*. In contrast, the genes that related to oligodendrocyte myelination (*MAG*, *MAL* and *MOBP*) and differentiation (*NKX6-2*, *OLIG1/2*, and *MYRF*) showed increased H3K27ac signals and reduced H3K27me3 and H3K9me3 signals in oligodendrocytes (Figure S4.2D).

Together, these results highlight the utility of Droplet Paired-Tag for resolving epigenetic regulatory programs across both spatial axes and developmental trajectories in the human basal ganglia.

### Conservation of MSN groups and epigenome between human and mouse

To assess the conservation of transcriptomic and histone modification landscapes between human and mouse, we integrated the human BG MSN Droplet Paired-Tag dataset with mouse MSN Paired-Tag dataset (Wang *et al.*, co-submit). Using the transcriptome as the integration anchor, joint clustering identified nine conserved MSN groups with similar proportions in both species, supporting broad conservation of MSN cell states (Figure 5A-B). To quantify divergence in gene expression, we performed differential expression analysis on orthologous genes using edgeR for each group between human and mouse (Figure 5C). Across nine groups, hundreds of orthologous genes exhibited species-biased expression. Humans had more species-biased genes than mice in 7 of 8 groups; only the STRv D1 NUDAP MSN group had more mouse-biased genes (Figure S5.1A). In addition, each group contained a subset of conserved orthologous genes with similar expression levels between species, the numbers of which was similar to the counts of mouse-biased genes. (Figure S5.1A). To identify biological processes displaying high levels of conservation and divergence, we performed GO enrichment analysis. Human-biased genes were enriched for responsiveness to monoamine and catecholamine stimuli, and chemotaxis (Figure S5.1B). Conserved orthologous genes were enriched for foundational neuronal functions, including axon extension, synaptic regulation, and neuromodulatory GPCR signaling (Figure S5.1C). Thus, while cell-type identities are broadly preserved, each group had a specific repertoire of hundreds of orthologous genes with species-specific expression patterns.

**Figure 5.**
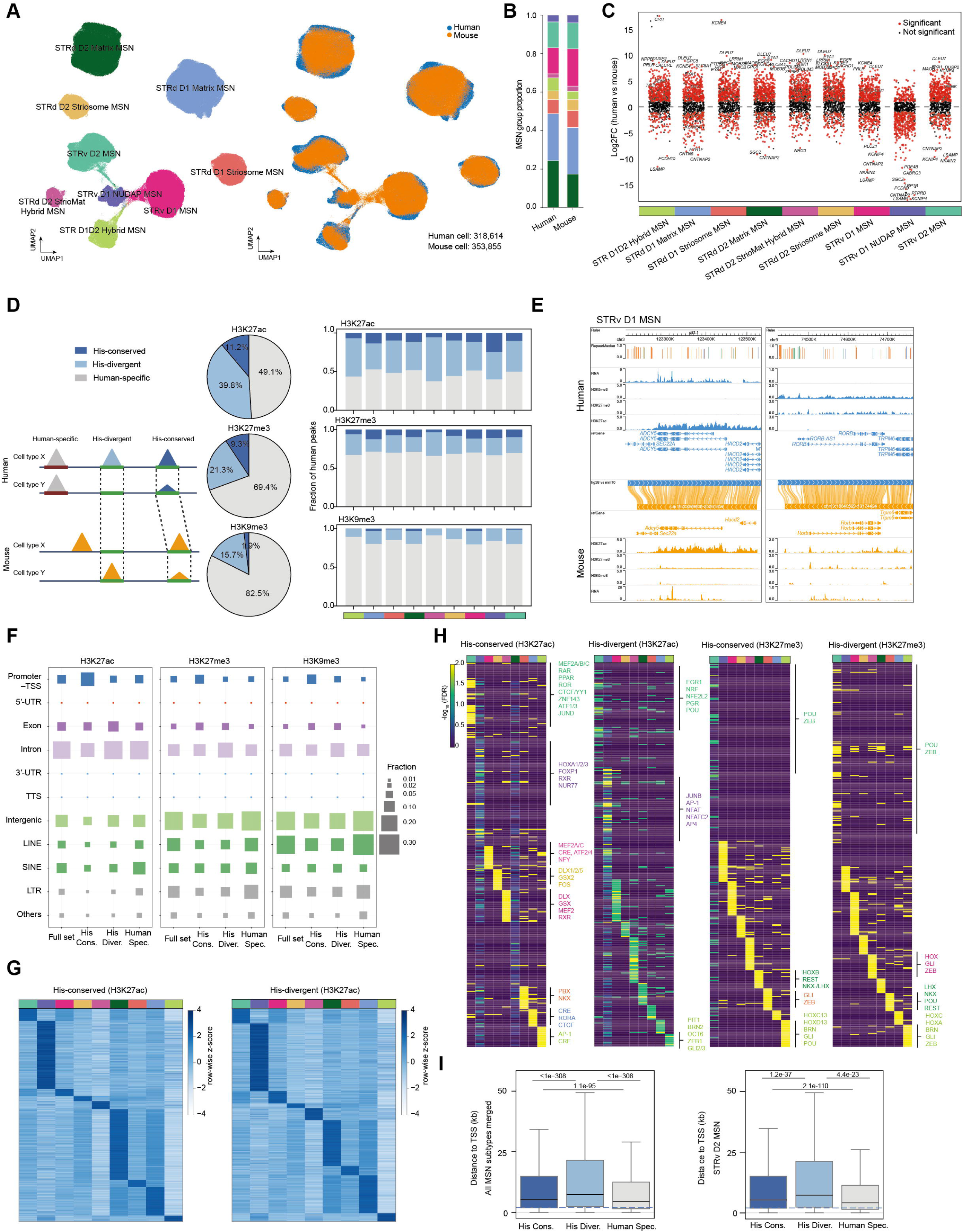
Comparative analyses between human and mouse MSNs. (A) UMAP co-embedding of 9 MSN groups from both human and mouse basal ganglia based on the transcriptome modality of Paired-Tag dataset (left). Cells were colored by species (right). (B) Stacked bar displaying MSN group proportions in human and mouse basal ganglia. Chi-square test, *P* value> 0.05. (C) Swarm plots showing the DEGs (Log_2_FC > 1 and P_adj_< 0.05) between human and mouse for each MSN group, where each dot represents a gene. (D) Left, schematic of three categories of histone modification peaks, including human-specific, His-divergent, and His-conserved peaks. The His-conserved peaks are conserved histone modification peaks in orthologous sequences. The His-divergent peaks are histone modification peaks only showing in one species in orthologous sequences. Human-specific peaks did not have orthologous regions in the mouse genome. Middle, pie charts showing fractions of three peak categories for each histone modification mark. Right, bar plots showing fractions of three peak categories in each MSN group. (E) Examples of His-conserved (left) and His-divergent peaks (right) at representative loci in WashU Comparative Epigenome Browser displaying alignment between human (hg38; top, blue) and mouse (mm10; bottom, orange) genomes with data tracks for RNA, H3K27ac, H3K27me3 and H3K9me3 in STRv D1 MSNs. (F) Dot plots showing the fractions of genomic features of three peak categories for each histone modification mark. (G) MSN group-biased conserved and divergent H3K27ac peaks. (H) TF motif enrichment in MSN group-specific conserved and divergent H3K27ac and H3K27me3 peaks. (I) Box plots showing the distance between three categories of H3K27ac peaks and TSSs in all MSNs and STRv D2 MSNs. Boxes indicate the interquartile range, center lines denote medians, and whiskers represent minimum and maximum values. Statistical significance was assessed using pairwise Wilcoxon rank-sum tests.

We further compared H3K27ac, H3K27me3, and H3K9me3 profiles between human and mouse within matched MSN groups defined by transcriptome integration. For this comparative epigenomic analysis, we defined conservation levels by integrating sequences and histone modification signals. For each histone mark, human peak genomic regions were mapped to syntenic loci in the mouse genome using liftOver, and conservation of histone modification signal was evaluated at the mapped intervals. For H3K27ac, we defined 11.2% of human peaks with both DNA sequence similarity and conserved modification signals as H3K27ac-conserved peaks, 39.8% of human peaks with only DNA sequence similarity as H3K27ac-divergent peaks, and 49.1% of human peaks with no orthologous mouse sequences as human-specific H3K27ac peaks (Figure 5D; Table S5.1). Similarly, we defined 9.3% H3K27me3-conserved peaks, 21.3% H3K27me3-divergent peaks, and 69.4% human-specific H3K27me3 peaks; and 1.9% H3K9me3-conserved peaks, 15.7% H3K9me3-divergent peaks, and 82.5% human-specific H3K9me3 peaks (Figure 5D; Table S5.2-5.3). The higher fraction of human-specific H3K27me3 and H3K9me3 peaks may be due to the broader peak structure of these repressive marks relative to H3K27ac, which increases the probability of spanning alignment gaps or poorly conserved regions and thereby reduces detection of continuous orthologous intervals in the mouse genome. To better visualize conserved and divergent histone modification tracks across MSN groups, we built an interactive comparative portal (https://wangcluster.wustl.edu/~wzhang/projects/MSN_epigenome/) (Figure 5E). We found that *ADCY5*, a key gene for the integration and control of dopaminergic signaling in MSNs^64^, exhibited conserved expression and histone modification patterns between human and mouse in STRv D1 MSNs. In contrast, *RORB*, a layer IV excitatory neuron marker^65^ and Alzheimer’s disease susceptibility gene^66^, showed mouse-specific active expression and H3K27ac peaks in STRv D1 MSNs, whereas the orthologous gene locus was silent in human STRv D1 MSNs and instead marked by repressive H3K27me3 and H3K9me3 peaks (Figure 5E).

We next annotated each category of histone modification peaks by standard genomic features. Across all three marks, histone-conserved peaks were preferentially enriched at promoter–TSS regions relative to divergent or human-specific peaks (Figure 5F), consistent with previous observations that conserved accessible cCREs are enriched at promoters^31^. In contrast, human-specific peaks were more enriched in TEs than histone-conserved or histone-divergent peaks, which likely contributes to the larger human-specific fractions observed for H3K27me3 and H3K9me3 relative to H3K27ac (Figure 5F). Previous studies reported that both human- and mouse-specific accessible cCREs are enriched in TEs^31^, suggesting that species may co-opt TEs to generate species-specific regulatory programs. Our results extended previous observations, showing that promoter-TSS enrichment of epigenomic conserved peaks and TE enrichment of species-specific epigenomic peaks is not limited to active elements but also applies to repressive elements.

Both histone-conserved and histone-divergent peaks displayed MSN group–specific patterns. Notably, STRv D1 and STRv D2 MSNs, as well as STRd D1 matrix and STRd D2 matrix MSNs, showed similar profiles for both conserved and divergent peaks, suggesting functional similarity despite belonging to different subclasses (Figure 5G and Figure S5.2A-B). To identify regulatory features associated with conservation levels, we performed TF binding motif enrichment analyses in histone-conserved and histone-divergent peaks and highlighted motifs significantly enriched in each MSN group with established roles in neuronal regulation. H3K27ac-conserved peaks were enriched for motifs associated with striatal identity modules and chromatin organization, including DLX/GSX, FOXP1, MEF2, ROR/RXR/Nur77, and CTCF/YY1/ZNF143, supporting deep conservation of core cell-identity regulatory elements (Figure 5H). In contrast, H3K27ac-divergent peaks showed stronger enrichment for stimulus-responsive motifs (EGR1, AP-1/JunB, and NFAT) and additional lineage-associated factors motifs, suggesting that evolutionary turnover is concentrated in inducible regulatory programs while foundational MSN identity circuitry remains conserved (Figure 5H). For H3K27me3, both conserved and divergent elements were dominated by motifs for developmental lineage TFs (POU, ZEB, HOX, NKX/LHX, GLI, and REST), consistent with a largely conserved repressive logic targeting fate and patterning programs (Figure 5H). Among enriched motifs, homeobox TF families (POU/HOX) were shared across H3K27ac and H3K27me3, conserved and divergent peaks, and multiple MSN groups, indicating their central roles in neuronal identity for both human and mouse (Figure 5H).

Finally, the distances from H3K27ac-divergent peaks to TSSs were larger than those from H3K27ac-conserved and human-specific H3K27ac peaks across all the MSN groups (Figure 5I and Figure S5.2C), indicating that H3K27ac-divergent peaks are more likely to represent distal enhancers, whereas H3K27ac-conserved and human-specific H3K27ac peaks tend to be proximal promoter. These results indicate that evolutionary changes in histone modification activity are less constrained at distal enhancers than at proximal promoters, whereas DNA sequence evolution may be more strongly constrained at distal enhancers than at promoters.

Collectively, we systematically assessed evolutionary conservation and divergence of MSNs between human and mouse at the levels of group composition, transcriptomes, and histone modification landscapes. These analyses indicate that MSN group identities are broadly conserved across species, particularly at promoter-proximal elements and core striatal identity modules spanning both active and repressive chromatin landscapes.

### Interpretation of risk variants of complex traits and diseases

Next, we sought to utilize BG cell-type-specific gene regulatory maps to interpret non-coding genetic risk variants associated with neuropsychiatric diseases and BG related traits. Using cell-type-resolved histone modification landscapes, we first prioritized BG cell types relevant to the different neuropsychiatric disorders. We collected genetic variants from 31 public GWAS studies (Table S6.1) and tested enrichment of trait heritability in histone modification peaks across BG subclasses using linkage disequilibrium score regression (LDSC), which identified functional enrichment in cell types and epigenomic regions from GWAS summary statistics using genome-wide information from all SNPs and explicitly models LD. We observed significant associations between neuropsychiatric disorders and intelligence-related traits^67–70^ with H3K27ac-marked genomic regions among multiple BG neuronal subclasses, whereas non–central nervous system traits showed little or no enrichment (Figure 6A and Figure S6.1A), consistent with previous observations that non-coding risk variants were enriched in accessible cCREs of disease-relevant cell types^33,71,72^. In contrast, a few enrichments of neuropsychiatric risk variants were observed within H3K27me3- and H3K9me3-modified peaks across BG neuronal cell types, while physical development traits^73–75^ showed enrichment in repressive-marked regions in neuronal cell types (Figure 6A and Figure S6.1A). Notably, genetic variants associated with neuropsychiatric disorders and intelligence also showed enrichment within H3K27me3 peaks across multiple non-neuronal cell types (Figure S6.1A).

**Figure 6.**
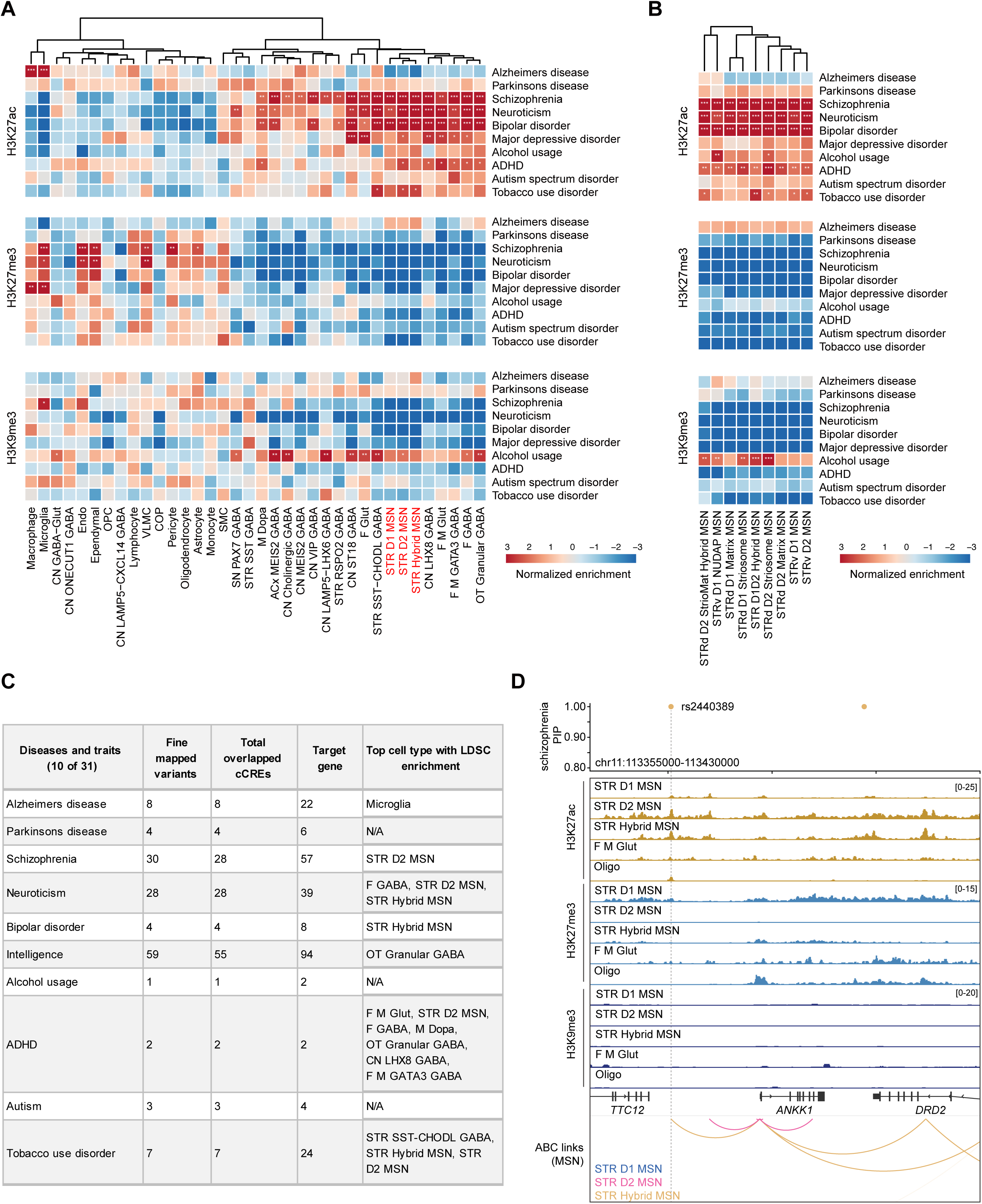
Interpreting noncoding risk variants of mental diseases and traits. (A) LDSC analysis results showing the risk variants associated with neuropsychiatric disorders and traits enrichment in three histone modification peaks from each major basal ganglia subclass. *FDR < 0.1; **FDR < 0.05; *** FDR < 0.01. (B) Heatmap showing enrichment of risk variants associated with neuropsychiatric disorders and traits in three histone modification peaks from each MSN group. *FDR < 0.1; **FDR < 0.05; *** FDR < 0.01. (C) Table showing the number of fine-mapped variants (PIP >0.1), number of cCREs overlapping with those variants, number of target genes linked with variants overlapping cCREs from ABC model, and cell types with top LDSC enrichment (FDR < 0.1) for selected diseases and traits. (D) Characterization of fine-mapped schizophrenia risk variant rs2440389 in a H3K27ac-marked enhancer. Genome browser tracks showing three histone modification profiles in five subclasses. Chromatin interaction tracks showing links between the variant-containing enhancer and putative target genes from the ABC model. All links showing with ABC score> 0.025. PIP, posterior inclusion probability.

The LDSC analysis results showed expected cell type–disease associations. For example, Alzheimer’s disease risk variants were strongly enriched in microglial H3K27ac peaks across two independent GWAS datasets^76–78^ (Figure S6.1A). We further found that the three MSN subclasses had strong heritability enrichment across multiple neurological and psychiatric disorders within H3K27ac peaks, including schizophrenia (SCZ)^79^, neuroticism^80^, bipolar disorder^81,82^, major depressive disorder^83^, attention deficit hyperactivity disorder (ADHD)^84^, and addiction-related tobacco use disorder^85^ (Figure 6A and Figure S6.1A). These neuropsychiatric traits also showed enrichment in H3K27ac peaks of other BG neuronal subclasses, including F M GATA3 GABA, M Dopa and F M Glut (Figure 6A). Interestingly, risk variants associated with alcohol usage^86^ were significantly enriched within H3K9me3 peaks in several neuronal subclasses (Figure 6A). For Parkinson’s disease^87^, which is linked to death of substantia nigra (SN) dopaminergic neurons and subsequent MSN dysfunction^88^, we observed no enrichment in M Dopa neurons or MSNs across the three histone modification landscapes (Figure 6A). Distinct BG regions are associated with specialized functions and differential disease vulnerability^89^. Thus, we reran the LDSC analysis at MSN groups level, which included regional information. Risk variants for several neuropsychiatric disorders were not uniformly enriched across different MSN groups (Figure 6B and Figure S6.1B). Notably, alcohol usage^86^ risk variants were preferentially enriched in H3K27ac peaks of STRv D1 NUDAP MSNs in the ventral striatum (Figure 6B), whereas tobacco use disorder^85^ variants showed strongest enrichment in STR D1D2 Hybrid MSNs (Figure 6B).

We next sought to interpret the regulatory mechanism underlying disease-associated non-coding risk variants in basal ganglia, based on the hypothesis that non-coding genetic variants acting within cCREs can alter the expression of disease-associated genes. First, we defined likely causal variants for neuropsychiatric diseases and traits from the 31 GWAS studies using the Bayesian fine-mapping method, FINEMAP (posterior inclusion probability [PIP]>0.1). Second, we used ABC model results to link these fine-mapping variants to their distal target genes based on cell-type-specific enhancer activity. Overall, we detected 1,251 likely causal variants residing within 1,208 ABC model-defined enhancers across different BG cell types, and linked to 2,526 putative target genes (Figure 6C; Table S6.2-6.3). For example, a schizophrenia-associated likely causal variant (rs2440389) overlapped an H3K27ac-marked enhancer, which was predicted to regulate *ANKK1* gene in STR Hybrid MSNs by the ABC model, nominating a potential cell-type-specific regulatory mechanism for schizophrenia susceptibility^90^ (Figure 6D).

Overall, these cell-type-resolved histone modification landscapes and catalogs of gene regulatory elements provide a framework for linking GWAS variants to specific BG cell types, regulatory elements, and putative target genes, thereby expanding mechanistic hypotheses and candidate therapeutic targets for neuropsychiatric disease.

### A sequence to function model predicts cell-type-specific gene regulation in the basal ganglia

Deep-learning models have become powerful tools for dissecting gene regulatory programs^91–94^. Sequence-based models (sequence to function, or S2F) that can predict gene expression and epigenomic states have been developed through training with large compendia of bulk tissue datasets. However, extending these approaches to individual brain cell types has been limited by the lack of large-scale single-cell epigenomic profiles, particularly for different histone modifications. Leveraging our Droplet Paired-Tag atlas, we developed a sequence-based deep-learning framework, EpiBRAIN (Epigenomics-based Brain Regulation Attention Inference Network), for predicting human brain cell-type-specific gene expression and multiple epigenomic profiles. EpiBRAIN was built upon the architecture of Borzoi^93^, a recently released S2F model that trained on bulk-tissue CAGE, DNase, ATAC-seq, ChIP-seq, RNA-seq datasets. EpiBRAIN processed a 524-kb input from the human reference genome (hg38) and predicted cell-type-specific ATAC-seq, H3K27ac, H3K27me3, H3K9me3, and RNA-seq profiles at 32-bp resolution (Fig. 7A, Fig. S7.1A, and Methods). In contrast to Borzoi, which treats each cell type and modality as an independent track, EpiBRAIN model employs a cell-type head architecture that predicts cell-type-specific chromatin embeddings, followed by shared modality heads. This design allows the model to efficiently capture correlations between modalities within each cell type. This architectural adaptation demonstrated superior performance compared to Borzoi’s default structure when evaluated on our datasets (Fig. S7.1B). We trained EpiBRAIN model on 37 distinct cell types (subclasses with ≥ 1000 cells) derived from the basal ganglia and cortex and evaluated its predictive accuracy on held-out testing data (Methods).

**Figure 7.**
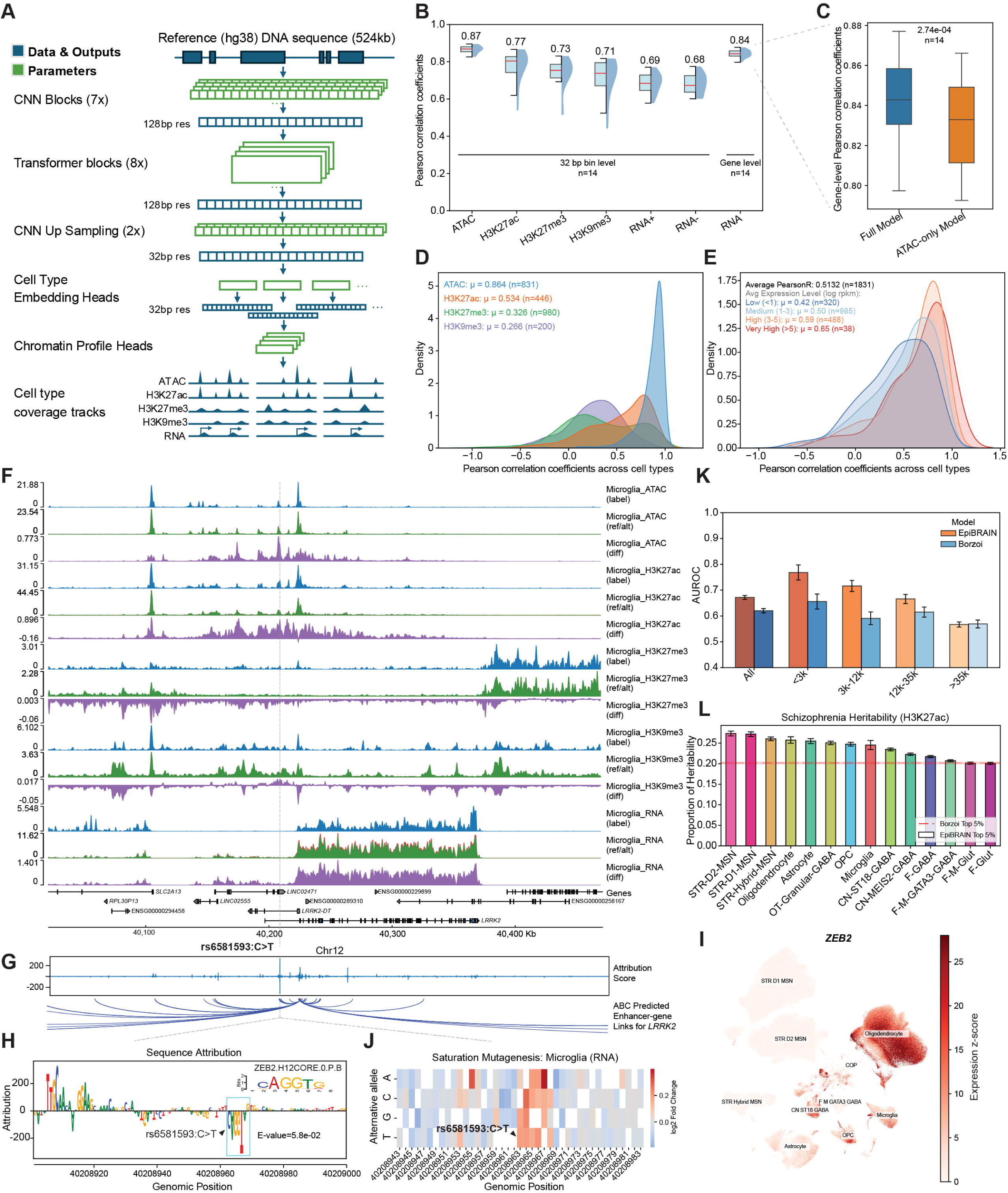
A deep learning model for predicting cell-type-specific epigenomic profiles, gene expression, and non-coding variant effects from DNA sequence. (A) Schematic of the EpiBRAIN deep learning model architecture. The model took a 524 kb genomic DNA sequence as input and processed it through CNN and Transformer blocks to predict multimodal outputs, including chromatin accessibility, histone modification profiles and gene expression, at 32-bp resolution. (B) Predictive performance of EpiBRAIN on a held-out test set. Box plots displaying the distribution of Pearson correlation coefficients between predicted and measured signals for five modalities across basal ganglia cell types (n=14). Performance is evaluated across chromatin accessibility (ATAC), histone modifications, and gene expression (RNA+ for forward strand, RNA- for reverse strand, RNA for reads per kilobase of transcript per million mapped reads (RPKM) aggregated by genes). The mean numbers were labeled on the top of the boxes. (C) Impact of training datasets on gene expression prediction. Box plots comparing the gene-level Pearson correlation coefficients of the EpiBRAIN model trained solely with ATAC-seq data (“ATAC-only Model”) versus the one trained with all the modalities (“Full Model”) across different cell types (n=14). Paired *t*-test. (D) Evaluation of cell-type specificity for epigenomic modalities predictions. Density plots showing the distribution of Pearson correlation coefficients calculated across cell types for cCREs with high variability (coefficient of variation > 1). Mean numbers were labeled for each group. (E) Evaluation of cell-type specificity for gene expression predictions. Density plots showing the distribution of Pearson correlation coefficients calculated across cell types for 1,831 genes, stratified by average expression levels. Mean numbers were labeled for each group. (F) In silico mutagenesis of the rs6581593 (C>T) variant in microglia. IGV tracks displaying the ground truth signals (“label”), model predictions of reference and alternative alleles (“ref/alt”), and the difference between ground truth and predictions (“diff”) for ATAC-seq, histone marks, and RNA-seq. (G) Sequence attribution analysis for *LRRK2* gene. Top, sequence attribution scores indicating the contribution of genomic regions to *LRRK2* expression (positive scores indicating promotion; negative scores indicating repression). Bottom, chromatin loops (enhancer-gene links predicted by ABC model) connecting distal regulatory elements to the *LRRK2* promoter. (H) High-resolution sequence attribution at the rs6581593 locus. The ZEB2 motif was predicted to be disrupted by rs6581593 (C>T) is highlighted by blue box, annotated with the motif matching score from HOCOMOCO v12. (I) UMAP visualization of basal ganglia cells colored by *ZEB2* gene expression level. (J) Saturation mutagenesis heatmap centered on the rs6581593 locus. Colors represent the predicted log2 fold change in *LRRK2* expression in microglia induced by each mutation. (K) Benchmarking variant prioritization performance. Bar plots comparing the Area Under the Receiver Operating Characteristic curve (AUROC) of EpiBRAIN and Borzoi in distinguishing causal eQTL variants (GTEx) in the basal ganglia. Variants are categorized by distance to the TSS. Mean ± SE (bootstrapping n=100). (L) Partitioned heritability of Schizophrenia GWAS variants estimation using stratified LD score regression (S-LDSC). Bar plots representing the proportion of heritability explained by the top 5% (59430) variants prioritized by EpiBRAIN model (colored bars) based on cell-type-specific H3K27ac attribution scores. The dashed line indicating the proportion of heritability explained by the top 5% of variants prioritized by the Borzoi model. Mean ± SE. For the box plots, hinges represent the 25th to 75th percentiles, the center line denotes the median, and whiskers represent the minimum and maximum values.

EpiBRAIN model achieved average Pearson correlation coefficients (PCCs) exceeding 0.80 between predicted and observed signals across cell types for chromatin accessibility and gene expression, and ∼0.7 for histone modifications across BG cell types (Fig. 7B). To test whether including histone modification profiles as training datasets can improve gene expression prediction beyond chromatin accessibility alone, we trained an ablation model using only single-cell ATAC-seq data, which reduced gene expression prediction accuracy across all cell types (Fig. 7C and Fig. S7.1C), indicating that histone modifications contained extended gene regulatory grammar. We further observed that the performance of EpiBRAIN model prediction generally scaled with the number of cells per cell type, approaching saturation for cell types with more than 1,000 cells (Fig. S7.1D–E). To confirm that EpiBRAIN model can capture cell-type-specific variability rather than constitutive genomic features, we evaluated performance on highly variable cCREs. The model maintained average PCCs of 0.5–0.8 for ATAC and H3K27ac signals (Fig. 7D) and ∼0.5 for gene expression across cell types (Fig. 7E). Notably, EpiBRAIN model showed stronger predictive performance for cCREs identified as regulatory by the ABC model compared with the other cCREs (Fig. S7.1F).

We next investigated the utility of EpiBRAIN model in elucidating the molecular mechanisms underlying neuropsychiatric disease variants. We focused on rs6581593 (C>T), a Parkinson’s disease–associated variant for which the mechanisms linking GWAS variants to pathogenicity remain unclear. EpiBRAIN model predicted that rs6581593, which resides within an cCRE, specifically altered the microglial chromatin landscape by increasing chromatin accessibility and H3K27ac signals while decreasing H3K27me3 signals (Fig. 7F). Sequence attribution analysis highlighted this element as a putative regulator of *LRRK2* gene (Fig. 7G), consistent with ABC model-predicted enhancer-target gene links as well as previous study’s CRISPRi experiment result^95^. Accordingly, the risk allele was predicted to drive microglia-specific upregulation of *LRRK2* (Fig. 7F and Fig. S7.2A), which aligns with prior reports of increased *LRRK2* expression in microglia in Parkinson’s disease^95^. Mechanistically, sequence attribution revealed that the variant disrupted a binding motif for the transcriptional repressor ZEB2 (Fig. 7H), which is expressed in microglia (Fig. 7I). In silico saturation mutagenesis further indicated that disruption of this motif increased predicted *LRRK2* expression (Fig. 7J), supporting a derepression model of rs6581593 (C>T). Although previous study did not directly perform CRISPR–Cas9 base editing of rs6581593 to test its causal role in *LRRK2* expression, iPSC-derived iMicroglia carrying the Parkinson’s disease risk haplotype rs76904798-CT and rs6581593-CT showed a statistically significant genotype-dependent increase in LRRK2 protein levels and activity^95^. In addition, EpiBRAIN model also correctly predicted that rs76904798 and rs1491942 did not alter *LRRK2* expression (Fig. S7.2B), consistent with their CRISPR–Cas editing experiment results showing that these two individual SNPs do not change *LRRK2* expression in iMicroglia^95^.

Finally, we assessed the performance of the EpiBRAIN model to prioritize causal variants in the context of eQTLs and GWAS. When distinguishing fine-mapped basal ganglia eQTL variants from negative controls, EpiBRAIN model outperformed Borzoi for proximal variants while achieving comparable performance for long-distance variants (Fig. 7K). Moreover, EpiBRAIN-predicted cell-type specific effect sizes correlated with the beta values of fine-mapped eQTL variants from independent brain studies^96,97^ (Fig. S7.2C–D). In the analysis of schizophrenia heritability, the top 5% of variants predicted by the EpiBRAIN model that had the largest effects on H3K27ac in MSNs explained ∼25% of disease heritability, representing an improvement over variants prioritized by Borzoi (Fig. 7L).

Collectively, these results demonstrate that the EpiBRAIN model can predict epigenomic and transcriptional landscapes, identify sequence motifs that drive regulatory element function, and predict the effects of non-coding variants implicated in neuropsychiatric disease in cell-type-specific manners.

## DISCUSSION

The basal ganglia are essential to the orchestration of reward, emotion, cognitive, and motor circuits, and implicate in a broad spectrum of neurological and psychiatric disorders, yet the cell-type-resolved regulatory programs that underlie BG function and vulnerability remain incompletely defined. Here, we present a single-cell atlas that jointly profiles histone modification (H3K27ac/H3K27me3/H3K9me3) together with the transcriptome in the same nuclei across eight BG regions, and provide access to these data through an interactive portal. Utilizing this multimodal dataset, we annotated chromatin states across ∼50% of the genome, and identified 356,380 candidate functional enhancers within E-P loops, 179,396 candidate silencers within S-P loops, and 54,847 super-enhancers. By integrating cCREs with TF motif information and target gene expression using SCENIC+ and refining enhancer–gene links with the ABC model, we further inferred cell-type-specific GRNs across BG subclasses. Together, this atlas provides a systematic annotation of chromatin states and epigenomic regulatory programs in the human basal ganglia. It expands upon previous accessibility-based cCREs annotation and provides a more comprehensive view of both active and repressive epigenomic regulation.

Despite the well-documented antagonistic roles of STR D1 and STR D2 MSNs in motor control, global comparative analyses revealed striking similarities in their transcriptomic profiles and active regulatory landscapes^11^, including H3K27ac-defined cCREs, E-P loops, and ABC model-predicted enhancer–gene links. Intriguingly, we observed that the distinctions between these two MSN subtypes resided in their silencer repertoires identified from H3K27me3 based S-P loops, accompanied by subtype-specific TF motif enrichment. This pattern supports a model in which repressive chromatin stabilizes neuronal identity and constrains regulatory programs relevant to neuronal function. Consistent with this model, loss of PRC2 in MSNs triggers inappropriate gene derepression and neurodegeneration^54^. This model provides a plausible route by which closely related MSN subtypes could adopt distinct, signal-tuned silencing programs, and indicates the necessity of investigating repressive chromatin states. By integrating DEGs with GRNs inferred from the SCENIC+ and the ABC model, we identified subtype-specific regulatory TFs and enhancers, with D1MSN highlighting dopamine-responsive immediate-early regulators (e.g. NR4A2, JUNB)^98,99^ whereas D2MSN shows higher activity for TFs associated with corepressor-mediated transcriptional constraint (e.g. NCOR1)^100^. These findings showed a framework for disentangling the fine-tuned regulatory programs that govern closely related cell types.

Utilizing the matched sample dissection between MERFISH and Droplet Paired-Tag, and single-cell spatial resolution of MERFISH images, we imputed spatial coordinates for individual Droplet Paired-Tag nuclei and reconstructed region-specific spatial epigenomic maps. This strategy revealed gradient changes in histone modifications at gene bodies and enhancers along the anatomical BG axes, which paralleled transcriptomic gradients and provided a mechanistic foundation for regional function specialization. Notably, although the caudate nucleus is often treated as a uniform structure, we observed robust A–P gradients in both gene expression and histone modifications within STR D2 MSNs from head to tail. Previous studies have observed a different pattern of functional connectivity gradient with other brain regions for the caudate nucleus’s A-P axis by fMRI^59^. Moreover, fMRI captured a gradual transition of performance-related activity from the head to tail of the caudate nucleus while participants learned to control a cursor by moving fingers^60^. The transcriptomic and epigenomic gradient changes along head to tail of the caudate nucleus may serve as the molecular foundation of its connectivity and functional gradients. For example, *NPAS4* performs activity-dependent regulation of inhibitory synapse development^101^ and its gradient expression along the A-P axis of the caudate nucleus is consistent with heightened activity-responsive transcriptional programs in the caudate head. In addition to resolving spatial heterogeneity, the Droplet Paired-Tag dataset also captured developmental dynamics of epigenome during OPC-to-oligodendrocyte differentiation.

Cross-species comparison of transcriptome and three histone modification landscapes across nine conserved MSN groups between human and mouse revealed both evolutionary conservation and divergence at the level of transcriptome and epigenomic regulation. In general, MSN group identities are broadly conserved between human and mouse in transcriptome, and conserved histone modification peaks are preferentially proximal to TSSs and enrich core striatal identity TF binding motifs. Conservation of chromatin regions that retain both sequence and cell-type-specific modification in mouse and human are likely under evolutionary constraint, indicating functional importance. Divergence is concentrated in distal regulatory elements, TEs and stimulus-responsive regulatory programs. H3K27ac landscapes showed higher conservation than H3K27me3 and H3K9me3, likely due to the broad peak structure of repressive marks and their enrichment in TEs and other rapidly evolving sequences.

Across conserved H3K27ac and H3K27me3 peaks, homeobox TF motifs were repeatedly enriched across multiple MSN groups, implicating homeobox TFs as central components of conserved neuronal identity. Independently, SCENIC+ eRegulons highlighted homeobox TFs as core regulators across BG cell-type-specific gene regulatory networks. We speculated that human had an evolutionarily conserved principle from *C. elegans* that distinct neuronal types were specified by combinatorial expression of homeobox TFs^51–53^. Indeed, we observed a unique combinatorial homeobox TF expression pattern for each BG subclass. These homeobox TF gene loci displayed high promoter-proximal H3K27ac modifications in the cell types where they were expressed, but were marked by H3K27me3 and H3K9me3 in non-expressing cell types. Notably, homeobox genes are canonical targets of the PRC2 and are often regulated by an epigenetic “on/off” switch during neuronal development. Recent study showed that genes encoding homeobox TFs were enriched among up-regulated genes in mouse PRC2-deficient MSNs in the striatum, and associated neurodegenerative changes in MSNs^54^. These observations suggest that homeobox TF programs in the basal ganglia are tightly constrained by histone modification states and may be linked to disease vulnerability. In addition, chromatin accessibility at specific combinations of homeobox TF motifs and co-TF motifs, shows strong cell-type specificity, which can regulate downstream gene expression. Together, these results suggest that combinatorial homeobox TF expression patterns shaped by chromatin states, and combinatorial motif accessibility grammar, help orchestrate neuronal identity programs.

A key motivation for building a cell-type-resolved gene regulatory atlas is to facilitate interpretation of disease-associated genetic variations, particularly non-coding variants whose mechanisms often remain unclear. By integrating cell-type-specific epigenomic and transcriptomic features, our study extended functional annotation beyond coding sequences and linked risk variants to regulatory elements and target genes at cell-type resolution. First, LDSC analysis integrating GWAS summary statistics with cell-type-specific histone modification landscapes identified trait enrichments across BG cell types and histone-marked genomic regions. As a positive control, Alzheimer’s disease risk variants were strongly enriched in microglial H3K27ac peaks. Besides, risk variants of schizophrenia, intelligence, bipolar, major depressive disorder, ADHD, neuroticism, insomnia, tobacco use disorder showed enrichment in neuronal H3K27ac peaks. In contrast, enrichments within neuronal H3K27me3- and H3K9me3-peaks were generally weak for neuropsychiatric diseases, although alcohol usage showed a notable enrichment that needs further investigation. Second, fine-mapped likely causal variants were overlapped with cell-type-specific cCREs and linked to putative target genes via ABC enhancer–gene connections, providing candidate effector genes and regulatory mechanisms for BG-relevant disease risk.

Deep learning models that extract regulatory context directly from DNA sequence have emerged as powerful tools for predicting functional genomic features, including chromatin accessibility, 3D interactions, and gene expression. Existing sequence-to-function models have limitations. Epiformer^31^, which is trained on cell-type-resolved sci-ATAC-seq data, did not directly predict gene expression. The Borzoi^93^ model was trained primarily on bulk datasets and therefore lacks cell-type resolution. Recently, Borzoi-prime^102^ model has incorporated single-cell ATAC-seq for cell-type-resolved training; however, it still lacks histone modification information, limiting their ability to capture chromatin-state complexity beyond accessibility. We developed a deep learning model EpiBRAIN based on the architecture of Borzoi and trained on cell-type-specific profiles of three histone marks together with chromatin accessibility to predict chromatin states and gene expression in cell-type-specific manners. We demonstrated that integrating histone modification profiles yields better performance compared to models trained solely on snATAC-seq data. Furthermore, our model improved the prediction of brain-specific causal eQTLs and the prioritization of neuropsychiatric disease-associated variants relative to the widely used Borzoi model. These improved performances likely reflect both the broader genomic coverage and the richer cell-type-specific chromatin-state information provided by multiple histone modifications. Notably, the model successfully predicted the regulatory impact of a Parkinson’s disease risk variant, forecasting an increase in *LRRK2* expression in microglia that was consistent with patient phenotypes. Beyond corroborating the microglia-specific effect observed in previous studies, our model provided a novel mechanistic hypothesis: the disruption of a binding motif for the transcriptional repressor ZEB2 by the risk allele leads to *LRRK2* up-regulation. Together, these results demonstrate that integrating histone modifications with sequence-based model enables robust *in silico* interpretation of non-coding disease variants and provides a scalable framework for prioritizing candidate regulatory mechanisms and therapeutic targets in neuropsychiatric disorders.

This study defined a cell-type-resolved catalog of active and repressive regulatory elements in the human basal ganglia, providing potential targets for next-generation precision genomic medicines. Recent success^103^ of enhancer-directed CRISPR therapy underscores that high-resolution annotation of the non-coding genome can be translated into therapeutic interventions. More broadly, because post-translational histone modifications are reversible, targeted manipulation of histone marks offers an additional therapeutic avenue to reset regulatory states at disease-associated loci and, in turn, reshape downstream molecular programs and disease-relevant phenotypes. The datasets generated in this study provide a useful resource for such cell-type-specific epigenome-editing strategies.

### Limitations of the study

This study has several limitations. Our cohort included only seven donors (five males and two females), limiting power to study sex-associated effects. Some rare cell types remain under-sampled and can’t provide saturation peaks. In addition, the dissection boundaries of GP were imperfect and introduced contamination by MSNs. Future studies with larger cohorts, improved microdissection, and expanded data scale will further refine basal ganglia regulatory maps and strengthen their utility for mechanistic and translational discovery.

## RESOURCE AVAILABILITY

### Lead contact

Bing Ren

### Materials availability

This study did not generate new unique reagents.

### Data and code availability

- Human Droplet Paired-Tag datasets have been deposited at NeMo and are publicly available as of the date of publication.
- MERFISH datasets have been deposited at BIL and are publicly available as of the date of publication.
- Any additional information required to reanalyze the data reported in this paper is available from the lead contact upon request.

## ACKNOWLEDGMENTS

This publication was supported by and coordinated through the Brain Initiative Cell Atlas Network (BICAN), and NIH grant UM1 MH130994 (to B.R., J.R.E., X.X., T.W. and M.M.B.). We thank QB3 MacroLab for purifying the proteins used in this study. This work was supported by the Flow Cytometry Core Facility of the Salk Institute (RRID:SCR 014839) with funding from NIH-NCI CCSG: P30 CA01495, and Shared Instrumentation Grants S10-OD023689 (Aria Fusion cell sorter) and S10 OD034268 (Thermo Fisher Bigfoot). We gratefully acknowledge the Broad Institute and NEMO for conducting the sequencing and data storage. Z.W. is a D.D.Brown Awardee of the Life Sciences Research Foundation.

## AUTHOR CONTRIBUTIONS

Brain collection: J.B., R.D.H., C.D.K., E.S.L., X.X.

Tissue dissection and nuclei preparation: J.A.R., C.T.B., A.S.B., J.K.W., K.W.K., K.G.R., J.L., S.C., J.A., C.K.Y., G.V.S., A.C.M., Y.S., A.B., C.O., M.L., M.V.M., C.R., S.N.A., N.S.

Droplet Paired-Tag data generation: L.C., A.L., H.S.I., K.D., T.L., Z.W., X.G., C.Z., N.R.Z.,

MERFISH data generation: J.C.K., Z.Z., J.F., A.M., J.O., Q.Z., C.T.B., A.S.B., K.W.K., K.G.R., J.L., C.K.Y., C.B., E.O., W.O.

Droplet Paired-Tag data analysis: K.L., Y.X.

MERFISH data analysis: Y.X., E.B., J.C.K.

Deep learning model construction: G.Z., D.L.

Data visualization: W.Z., D.L., T.W.

Data sharing: Z.W., S.Z., W.D., A.K., Y.F., N.J., T.E.B.

Data interpretation: L.C., K.L., Y.X., G.Z., J.A.R., C.T.B., W.D., M.M.B., B.R.

Manuscript writing: L.C., K.L., Y.X., G.Z., J.A.R., C.T.B., Q.Z., M.M.B., B.R.

Supervision: B.R., M.M.B., J.R.E.

## DECLARATION OF GENERATIVE AI AND AI-ASSISTED TECHNOLOGIES

During the preparation of this work, the authors used OpenAI ChatGPT and Google Gemini to assist with selected coding tasks and to refine the manuscript for readability. All AI-assisted outputs were reviewed and edited by the authors, who take full responsibility for the final content of the publication.

## STAR★METHODS

## METHOD DETAILS

### Human Subjects

For this study, basal ganglia samples from four neurotypical donors (two males aged 50 and 47 years, and two females aged 63 and 67 years) were obtained from the UC Irvine (UCI) biobank and the BioRepository and Integrated Neuropathology (BRaIN) Laboratory at the University of Washington. In addition, basal ganglia tissues from three additional human donors (males aged 29, 42, and 58) obtained from the Allen Institute were also included, which had previously been analyzed using snATAC-seq^31^ and snm3C-seq^58^. Postmortem human brain tissues were obtained under protocols approved by the University of California, Irvine Institutional Review Board (IRB #1621). Written informed consent for brain donation was obtained from the donor’s next of kin prior to recovery. Donors ranged in age from 2 to 65 years and had a postmortem interval (PMI) of less than 30 hours. Exclusion criteria included brain-related death, prolonged mechanical ventilation, fentanyl abuse, and major neurological or psychiatric disorders.

For each donor brain, tissue samples were taken from the frontal and occipital poles, the cerebellar cortex, and the midbrain for assessment of RNA quality. Total RNA was isolated using RNeasy Lipid Tissue Mini Kit (Qiagen 74804) and RNA Integrity Number (RIN) was determined using the Agilent 4150 Tapestation System (Agilent G2992AA). Only donor brains with RIN >6 for all regions assessed were used for the study, with the exception of UCI4723 cerebellar cortex (5.1) and UCI2424 occipital cortex (5.5) (Table S1.1).

### Brain Slabbing

Each brain was weighed, photographed, and bisected into two hemispheres. One hemisphere was fixed in 10% formalin for approximately five weeks, transferred to PBS containing 0.02% sodium azide, and then used for MRI and neuropathological assessment. From the remaining fresh hemisphere, the brainstem and cerebellum were first removed from the cerebrum. The cerebrum and cerebellum were then digitally scanned using a SHINING 3D EinScan Pro HD scanner, embedded in ice-cold alginate, and cut into 4 mm thick slabs, producing coronal slabs from the cerebrum and sagittal slabs from the cerebellum. The brainstem was cut transversely into 4 mm thick slabs. Each slab was photographed, labeled, flash frozen in 2-methylbutane cooled with dry ice (−40 °C), vacuum sealed and stored at −80 °C.

### Tissue Dissection

Regions of interest (ROIs) were identified and mapped on each brain slab by aligning each slab image to an interactive, digitally annotated human brain atlas^104^. Prior to dissection of ROIs, frozen slabs were transferred from −80 °C to −20 °C for several hours to allow for controlled thawing. Each mapped ROI was then dissected, minimizing white matter inclusion. Each dissected tissue sample was then vacuum sealed, stored at −80 °C.

### Anatomical criteria

For each brain, eight basal ganglia ROIs were anatomically mapped according to the following criteria: (1) The head of the caudate nucleus (CaH) was aligned to plates 16 through 18 of the Ding Atlas and microdissected to minimize contamination from the ventrally adjacent nucleus accumbens (NAC). (2) The body of the caudate nucleus (CaB) was aligned to plates 20 through 33 of the Ding Atlas and microdissected to minimize contamination from the anterior CaH and ventrally adjacent bed nucleus of stria terminalis (BNST). (3) The tail of the caudate nucleus (CaT) was aligned to plates 49 through 69 of the Ding Atlas and microdissected to minimize contamination from the medially adjacent hippocampus (HIP) and remove excess white matter. (4) The putamen (Pu) was aligned to plates 17 through 27 of the Ding Atlas and microdissected to minimize contamination from the medially adjacent globus pallidus (GP), laterally adjacent claustrum (Cla) and ventrally adjacent NAC. (5) The subthalamic nucleus (STH) was aligned to plates 37 through 49 of the Ding Atlas and microdissected to minimize contamination from the dorsally adjacent thalamus and ventrally adjacent midbrain. (6) The GP, including the external (GPe) and internal (GPi) segments and the ventral pallidus (VeP), was aligned to plates 22 through 42 of the Ding Atlas and microdissected to minimize contamination from the laterally adjacent Pu and ventrally adjacent structures of the basal forebrain (BF). (7) The NAC was aligned to plates 18 through 22 of the Ding Atlas and microdissected to minimize contamination from all adjacent regions: CaH, Pu, BNST, A25, BF, VeP, and septal nuclei (SEP). (8) The gray matter of midbrain 1 (MGM1) sample was collected from the two dorsal-most midbrain slabs, where the substantia nigra (SN), red nucleus (RN) and ventral tegmental region (VTR) were microdissected together.

### Microdissection

Using the anatomical criteria described above, tissue blocks were microdissected on an aluminum block surface chilled to -23°C inside a Leica cryostat (Leica CM1950). Tissue was cut with a nuclease-free razor blade (Genesee, 38-101) using steady, perpendicular see-saw motions to avoid frozen tissue fracturing. Distribution of microdissected tissue was as follows: a 7x7mm tissue block at full thickness for spatial transcriptomics (MERFISH 2.0), plus additional tissue for single-nucleus isolation to be used for both Droplet Paired-Tag and snm3C-seq in the companion study (Ding *et al.*, co-submit). Targeting one million neuronal nuclei per sample to be split between both modalities (750K for DPT, 250K for snm3C-seq), the tissue mass required for each region was determined to be the following: CaH (270 mg), CaB (320 mg), CaT (400 mg), Pu (248 mg), STH (300 mg), GP (1,200 mg), NAC (118 mg), and MGM1 (900 mg), where all available tissue was used for CaT, STH, GP, and MGM1.

### Cryosectioning for MERFISH 2.0 and NISSL staining

Each 7x7mm tissue block for spatial transcriptomics was embedded in Tissue-Plus™ O.C.T. Compound (Fisher Scientific, 23-730-571), sectioned at a thickness of 10 μm on a Leica Cryostat (Leica CM1950) using Epredia™ MX35 Premier™ low-profile blades (VWR, 89238-778), and mounted onto MERSCOPE Large Slide V 2.0 (Vizgen, 20400118). Sections were fixed in 4% paraformaldehyde (Electron Microscopy Science, 15714-S) diluted in 1X phosphate-buffered saline (Thermo Fisher Scientific, AM9625) at 47 °C for 30 min, permeabilized overnight in 70% ethanol and photobleached for 5 hours using a MERSCOPE photobleacher (Vizgen; 10100003). Additional sections were also collected on Fisherbrand™ Superfrost™ Disposable Microscope Slides (Fisher Scientific, 12-550-123), fixed in 4% paraformaldehyde (Electron Microscopy Science, 15714-S) diluted in 1X phosphate-buffered saline (Thermo Fisher Scientific, AM9625) at 4 °C for 15 min, stained with 0.04% cresyl violet acetate working solution (Fisher Scientific, cat. no. 50-318-82) for 20 min, rinsed in distilled water, dehydrated through graded ethanol solutions (50%, 95%, 100%), cleared in Xylene solution (Fisher Scientific, X5-4), and coverslipped for histological analysis.

### Tissue Grinding

Frozen tissue dissections containing the brain regions of interest were combined across slabs as needed and ground on dry ice using a porcelain mortar (VWR 470149-118) and pestle (VWR 470149-080). The resulting powder was portioned into 150-250 mg aliquots in 1.5 mL microcentrifuge tubes on dry ice and stored at -80°C until downstream processing.

### Nuclei Isolation and FACS

Recombinant RNase inhibitor (RNH1) was purified from the plasmid pmal_c5x_RNAse_Inhib (Addgene plasmid # 153314; RRID: Addgene_153314), a gift from Drew Endy & Philippa Marrack, by the QB3 MacroLab at UC Berkeley and reconstituted at a concentration of ∼10 mg/mL.

Frozen ground tissue was homogenized on ice in 9 mL of lysis buffer (0.32M sucrose, 0.1% Triton X-100, 5 mM MgCl2, 25 mM KCl, 10 mM Tris-HCl pH 8.0, 1 mM DTT, Roche Complete Mini Protease Inhibitor, 20 U/mL SUPERaseIN, and 20 μg/mL RNH1) using glass Dounce homogenizers. Sequential filtration of the resulting homogenate was performed using 70 μm (Corning, 352350) and 30 μm (Sysmex, 04-004-2326) cell strainers.

To remove myelin, homogenates were incubated with a sample-specific volume of myelin removal beads for 15 minutes at 4°C on a rotating mixer. Bead volume was adjusted for each sample based on estimated myelin content. Samples were then passed through LS columns (Miltenyi Biotec 130-042-401) mounted on a QuadroMACS Separator (Miltenyi Biotec, 130-091-051). LS columns were rinsed with 2 mL of wash buffer (0.1% Triton X-100, 5 mM MgCl2, 25 mM KCl, 10 mM Tris-HCl pH 8.0, 1 mM DTT, Roche COmplete Mini Protease Inhibitor, 20 U/mL SUPERaseIN, and 20 μg/mL RNH1) and the combined flow through was collected in a 15 mL conical tube.

Nuclei were then pelleted by centrifugation at 1,000g for 10 min at 4°C using a Thermo Scientific Sorvall Legend RT+ benchtop centrifuge (Thermo 75004377) with swinging bucket rotor (Sorvall 75006445). The supernatant was carefully aspirated and each pellet resuspended in 1 mL of sort buffer (DPBS, 1% BSA, 1 mM EDTA, 1mM DTT, Roche COmplete Mini Protease Inhibitor, 20 U/mL SUPERaseIN, and 20 μg/mL RNH1). Resuspended nuclei were then fluorescently labeled using Alexa Fluor 488-conjugated anti-NeuN antibody (Millipore MAB377X) and Hoechst 33342 (Invitrogen H3570) at a dilution of 1:1000 and incubated at 4°C for 30 minutes on a rotating mixer. Following staining, nuclei suspensions were diluted to ∼750K nuclei/mL with additional sort buffer and distributed into aliquots of ∼3 mL in 5 mL roundbottom tubes for FANS.

Single nucleus sorting was performed on a BD Influx cell sorter (BD Biosciences 646500) using the ‘1-drop pure’ sort mode with a 100 μm nozzle and sheath pressure of 20 PSI. 1X PBS was used as sheath fluid, and nuclei were maintained at 4°C during the sorting process. Hoechst-positive nuclei exhibiting 2N DNA content were first gated using a Side Scatter (SSC-Area) × Hoechst-Area plot. Debris was then excluded using a Forward Scatter (FSC-Area) × Side Scatter (SSC-Area) light-scatter plot, and remaining nuclei aggregates were excluded using FSC and SSC pulse Area– and Width–based parameters. Nuclei were lastly gated on NeuN signal (488), and NeuN-positive and NeuN-negative nuclei were sorted into separate 2 mL DNA LoBind tubes (Eppendorf 022431048) each containing 250 μL 5X collection buffer (DPBS, 5% BSA, Roche COmplete Mini Protease Inhibitor, 100 U/μL SUPERaseIN, and 100 μg/mL RNH1).

Collected nuclei were centrifuged at 300g for 5 min at 4°C followed by an additional centrifugation at 500g for 10 min using an Eppendorf microcentrifuge (Eppendorf 5417R) with swinging bucket rotor (Eppendorf A-8-11). The supernatant of each collection tube was removed down to ∼250 μL and all nuclei of like populations were combined and quantified.

### Droplet Paired-Tag

Droplet Paired-Tag was performed as previously described with minor optimization^37^. A step-by-step-protocol for library preparation is available here: https://www.protocols.io/view/droplet-based-single-cell-joint-profiling-of-histo-eq2ly6kergx9/v1.

NeuN^+^ and NeuN⁻ single nuclei were pelleted with the swinging bucket centrifuge (5804R, Eppendorf) and counted using the cell counter (RWD C100-Pro) with DAPI staining. To enrich for NeuN⁺ nuclei, NeuN⁺ and NeuN⁻ nuclei from the same tissue block were recombined, yielding approximately 1.2–1.5 million nuclei. The pooled nuclei were then divided equally into three separate tubes for profiling three distinct histone marks.

First, nuclei were permeabilized with OMNI buffer on ice for 4 minutes and washed once with MED#1 buffer. Then, the nuclei were incubated with 2 μL PA-Tn5 (0.4 mg/mL) and 2 μg antibody in MED#1 buffer overnight at 4 °C. PA-Tn5 and antibodies were then removed by washing with MED#2 buffer twice. Tagmentation was carried out in MED#2 buffer supplemented with 10 mM MgCl_2_ (Invitrogen, AM9530G) at 550 r.p.m. and 37 °C for 60 minutes in a ThermoMixer (Eppendorf), and the reaction was terminated by adding 2× stop solution. The nuclei were washed in 1× nuclei buffer once and counted again. From each tube, 18,000 nuclei were aliquoted and used for a single droplet generation reaction with the Chromium Next GEM Single Cell Multiome kit (10x Genomics, 1000283). Finally, DNA and RNA library amplification was performed according to the Chromium Next GEM Single Cell Multiome kit manual, with the following modifications: the starting amount of preamplification product for DNA library amplification was doubled, 13 amplification cycles were used for the DNA library, and a different SPRI bead size selection were applied for the DNA library (50 μL + 105 μL).

For this study, we used antibodies targeting three histone marks: H3K27ac (Abcam, ab4729), H3K27me3 (Abcam, ab192985), and H3K9me3 (Abcam, ab8898).

Libraries generated were sequenced on NovaSeq X (illumina) platform using standard illumina primers. For the DNA libraries, the read lengths are 100 + 8 + 24 + 100 (Read1 + Index1 + Index2 + Read2). For the RNA libraries, the read lengths are 28 + 10 + 10 + 90 (Read1 + Index1 + Index2 + Read2).

### Data preprocessing and quality control for Droplet Paired-Tag

Droplet Paired-Tag FASTQ files were demultiplexed using 10x Genomics cellranger-arc (v2.0.0) with the mkfastq command. DNA and RNA modalities were then processed separately using cellranger-atac (v2.0.0) and cellranger (v6.1.2), respectively. DNA and RNA barcodes were paired using a custom script based on the barcode correspondence generated by cellranger-arc, following the Droplet Paired-Tag protocol.

To identify high-quality nuclei, histone modification fragments from each sample were first aggregated and used for peak calling to support quality control. Narrow peaks were called for H3K27ac, and broad peaks were called for H3K27me3 and H3K9me3 using MACS2 (v2.1.2)^61^. For the histone modality, nuclei were filtered based on fragment counts and the fraction of reads in peaks (FRiP). Nuclei were retained only if they passed quality control in both the histone modification and transcriptome modalities, consistent with the Droplet Paired-Tag quality control workflow.

Before clustering, nuclei with high fractions of mitochondrial or ribosomal RNA reads were removed. Nuclei with extremely high read counts were excluded as likely doublets. Putative doublets in the chromatin modality were identified and removed using the Scrublet^105^ implementation in SnapATAC2^106^ (snap.pp.scrublet followed by snap.pp.filter_doublets) on a per-sample basis. Doublets in the RNA modality were identified per sample using DoubletFinder^107^, and nuclei flagged as doublets in either modality were excluded.

### Droplet Paired-Tag data analysis

#### Signal enrichment analysis

Signal enrichment over ChIP–seq peaks was calculated using deepTools^108^ (v3.5.5). Density plots and heat maps were generated to visualize signal levels, and peaks overlapping ENCODE blacklist^109^ (v2) regions were removed before enrichment calculation.

#### RNA preprocessing, quality control, and clustering

Single-cell RNA-seq data were processed using Scanpy^110^ (v1.11.1). Low-quality cells were removed based on standard quality control metrics, including total gene counts and the fraction of mitochondrial and ribosomal transcripts. Gene counts were normalized, and genes with very low expression were filtered out. The top 4,000 highly variable genes were selected and used for dimensionality reduction by principal-component analysis (PCA). For all datasets, the first 50 principal components were used for batch correction with Harmony^111^, followed by UMAP visualization and Leiden clustering. In addition, marker genes defined in the HMBA reference taxonomy were visualized to assess the correspondence between identified clusters and known cell types. Marker genes for each cluster were identified using Scanpy. For visualization, gene expression values were clipped to the 0–99.5% percentile range.

#### RNA Cell type annotation

Cell type annotation was performed using the hierarchical correlation mapping algorithm implemented in MapMyCell, with the consensus basal ganglia taxonomy (CCN20250428) as the reference. The reference taxonomy is organized as a hierarchical tree with neighborhood, class, subclass, and group levels. For each unlabeled cell, annotation started at the root of the tree. Marker genes for each neighborhood were used to compare the cell’s gene expression profile with the mean expression profile of each neighborhood, and the neighborhood with the highest correlation was selected. This step was repeated 1,000 times, each time using a random 90% subset of marker genes, and the neighborhood receiving the most votes was assigned. The same procedure was then applied step by step to assign class, subclass, and group identities within the selected branch. Using this approach, each cell was assigned a full hierarchical label, and a total of 61 cell types were identified.

#### Iterative clustering of single-nucleus DNA data

After quality control and doublet removal, we performed iterative clustering on the DNA (Histone modification) modality to define cell groups for downstream analysis. RNA-based cell type labels were transferred to the DNA dataset by matching paired barcodes. In brief, we converted RNA cell IDs to a common key (cell_id) and then merged RNA annotations into the DNA metadata table using this key. Donor, target, and region labels were parsed from sample names and stored in the DNA metadata for later batch correction and stratified checks.

#### Feature selection and dimensional reduction

We used SnapATAC2 (v2.50) to select informative genomic bins for clustering. We first selected candidate features (15Kbp genomic bins) and then removed bins with very low coverage across cells. We normalized the cell-by-bin matrix using TF–IDF and then applied truncated SVD for dimensional reduction. The first SVD component was removed to reduce the effect of global accessibility. The remaining components were used as the low-dimensional representation for clustering. Batch effects across donors were corrected using Harmony on the reduced space.

#### Graph-based clustering, iterative refinement, and entropy filtering

We built a k-nearest neighbor graph from the Harmony-corrected embedding and performed Leiden clustering. To improve cluster purity, we removed very small clusters (n < 10) and filtered out outlier cells with unusually large mean neighbor distances. We then performed a second round of clustering within each first-round cluster to further split heterogeneous groups. For cluster quality control, we computed Shannon entropy of the RNA-derived cell type labels within each cluster (or subcluster). Clusters with high entropy, which indicated mixed cell-type composition, were removed, and the neighbor graph and UMAP were recomputed on the remaining cells. UMAP was used for visualization and manual sanity checks using donor, region, and RNA-transferred labels.

### Identification of reproducible peaks

Peak calling was performed for each cell type following the ENCODE ChIP–seq analysis pipeline (https://www.encodeproject.org/chip-seq/histone/). Prior to peak calling, reads and cells were filtered using the same criteria as in previous studies to remove low-quality data at both bulk and single-cell levels. For each cell cluster, all properly paired reads were aggregated to generate a pseudo-bulk dataset, which was treated as one biological replicate. Two pseudo-replicates were then generated by randomly splitting the reads from the pseudo-bulk dataset into two equal halves. Peak calling was performed independently on each pseudo-replicate using MACS2. Reproducible peaks were identified by IDR analysis, and only peaks detected in both pseudo-replicates were retained for downstream analyses. Peaks overlapping ENCODE hg38 blacklist regions were removed. Peak sets from all cell types were then merged using BEDtools^112^ (v2.27.1) to generate a union peak list for the full dataset. Super-enhancers were identified using ROSE based on H3K27ac peaks for each cell type, with default parameters, and super-enhancers were defined separately for each cell type.

### ChromHMM analysis

Chromatin state annotation was performed using ChromHMM^41^ (v1.27) based on Paired-Tag profiles of H3K27ac, H3K27me3, H3K9me3, and HMBA snATAC-seq^11^. All chromatin state annotations reported here were generated specifically for this study and were not intended to replace or reproduce large-scale consortium-level annotations. Analyses were conducted at the subclass level. For each subclass, single-cell fragments from the three histone modification datasets and snATAC-seq were aggregated to generate pseudo-bulk datasets. Two pseudo-replicates were generated per subclass by randomly splitting fragments into two equal parts, each containing 50% of the reads.

Before model training, chromatin signals were binarized at 500-bp resolution using ChromHMM based on fragment-derived signal tracks, following the standard ChromHMM binarization procedure with default parameters. Only genomic bins supported by sufficient signal were retained. ChromHMM models were trained using the LearnModel function, testing models with 2 to 16 chromatin states. Model training was performed independently on each pseudo-replicate.

To determine the optimal number of chromatin states, we compared models using the CompareModels function in ChromHMM. Emission probabilities from models with fewer states were compared to those from higher-state models using Pearson correlation. This analysis was performed separately for the two pseudo-replicates. We observed that model similarity increased with state number and reached a plateau at nine states, indicating that additional states did not capture new combinations of chromatin signals. The 9-state model also showed high consistency between pseudo-replicates and clear, interpretable combinations of histone modifications and chromatin accessibility.

Based on these results, a 9-state model was selected for downstream analyses. Genome segmentation was performed separately for each pseudo-replicate using the MakeSegmentation function with default parameters. Final chromatin state assignments required concordant state calls between the two pseudo-replicates for each subclass. Genomic regions without reproducible state assignments were labeled as non-reproducible signal. Chromatin state tracks across subclasses were merged and summarized using BEDTools.

### Enhancer and silencer annotation

Hi-C loop annotations (Ding *et al.*, co-submit) were used as supporting information to assist the classification of distal regulatory elements linked to promoters, following previously described strategy^42^ for integrating 3D genome contact information with regulatory element annotation. All analyses were performed at the BG subclass level. For each subclass, loop coordinates were provided in BEDPE format and used only to define pairs of interacting genomic regions. Each loop was split into two anchors and assigned a unique loop identifier for downstream annotation.

Promoters were defined as ±2 kb windows centered on transcription start sites (TSS). TSS coordinates were obtained from a gene annotation table and inferred from transcript boundaries and strand information when not explicitly provided. These promoter intervals were compiled into a BED file and used as the reference promoter set. For each subclass, loop anchors were annotated by overlap with three feature sets: promoter windows, subclass-specific H3K27ac peaks, and subclass-specific H3K27me3 peaks, using BEDtools for overlap analysis.

Anchor labels were assigned using a priority-based rule. Anchors overlapping promoter windows were labeled Promoter. Remaining anchors were labeled Enhancer if they overlapped H3K27ac peaks but not H3K27me3 peaks and labeled Silencer if they overlapped H3K27me3 peaks but not H3K27ac peaks. Anchors overlapping both histone marks were assigned based on the larger base-pair overlap, while anchors without overlap to any feature were labeled Other. Loop-level classes were then derived from the labels of the two anchors and categorized as P–P, P–E, P–S, or Other. Annotated loops were exported for summary and visualization, and candidate enhancer and silencer sets were defined by collecting anchors labeled Enhancer or Silencer for downstream analyses.

### ABC model

We applied the ABC model^45,46^ as we described in the companion study (Ding *et al.*, co-submit) to predict cell-type-specific enhancer–promoter interactions using pseudobulk Hi-C contacts, gene expression, ATAC-seq consensus peaks and H3K27ac signal as input. Hi-C matrices were power-law normalized, and top 150,000 strongest ATAC-seq peaks were retained. ABC scores were computed for enhancers within 5 Mb of each promoter (threshold 0.025 per official guidance).

### Gene regulatory network analysis and integration with the ABC model

To limit bias from very large cell groups, the multiomic dataset was downsampled either across the full dataset or within each subclass. At most 20,000 nuclei were retained per subclass. This strategy preserved major regulatory signals while reducing over-representation of abundant populations.

Enhancer activity was represented by H3K27ac, which marks active enhancers and supports enhancer-driven regulatory interactions in the basal ganglia. Downsampled H3K27ac fragments were processed with pycisTopic to quantify accessibility-like signal and to generate binarized topic matrices. Gene expression count matrices, H3K27ac peak regions, and H3K27ac-derived topic matrices were used as input to SCENIC+^47^ (v1.0a1) to infer transcription factor (TF)–target gene relationships and regulon activity. SCENIC+ was run with default settings unless otherwise stated.

For TF prioritization and visualization, SCENIC+ outputs were integrated with enhancer–gene links from the ABC model. Regulon target sets were defined using SCENIC+ TF–target relationships. An ABC support score was calculated for each TF and cell type by averaging ABC scores across the TF’s target genes. Regulon specificity was computed by column-normalizing regulon activity across cell types (RSS-like scaling). TFs were ranked within each cell type using a combined score derived from regulon specificity and ABC support, and top-ranked TFs were selected for downstream visualization.

For interpretation, inferred regulatory networks were examined across annotated cell types. GRN outputs were also visualized by highlighting D1- and D2-enriched neuronal populations to illustrate axis-related regulatory programs. Core regulatory modules were selected based on SCENIC+ enrichment results and visualized using Cytoscape^113^.

### Homeobox motif activity, co-accessibility, and network analysis

We analyzed homeobox-related regulatory programs using HMBA snATAC-seq pseudo-bulk profiles at the BG subclass level. Peak signals were represented as genomic intervals (hg38) in an AnnData object, with one pseudo-bulk profile generated for each subclass.

To define the homeobox transcription factor (TF) set, we used a published curated list of human homeobox genes^114^ and matched these TFs to motif annotations. Motif scanning and deviation analysis were performed using pychromVAR, following the chromVAR framework. Peak sequences were extracted from the hg38 reference genome. GC bias was computed for each peak, and matched background peak sets were generated to control for sequence bias. Position weight matrices were obtained from the JASPAR2024^115^ CORE collection. Motif matches were called using a fixed stringency threshold (P value cutoff = 5 × 10⁻⁵). Motif deviation z-scores were then computed for each subclass pseudo-bulk profile and used as a measure of TF motif activity.

For visualization, motifs with high variance across subclasses were selected and displayed as clustered heat maps of deviation z-scores. Focused heat maps were also generated for major homeobox-related motif groups. To assess homeobox-centered co-accessibility, we identified co-active TFs by comparing motif deviation patterns across subclasses and cell types, and retained strong homeobox–cofactor pairs for downstream analysis. Regulatory networks were constructed with TF motifs as nodes and homeobox–cofactor relationships as edges, with edge attributes indicating the supporting cell type. Network visualizations were generated either in Cytoscape for selected subnetworks or using a custom concentric layout for global views, in which node color represents TF family and per-cell-type activity is shown as an activation ring around each node.

### Basal ganglia regional axes from RNA and chromatin profiles

Regional axes within basal ganglia STR D1 MSN and STR D2 MSN populations were inferred using integrated RNA and chromatin profiles. Cells were first grouped by cell identity and anatomical region. For each group, normalized gene expression matrices were reduced by principal component analysis, and the top components were used to construct a k-nearest neighbor graph. To reduce noise, regional information was smoothed by averaging region labels across each cell and its neighbors.

A one-dimensional regional index was then assigned to each cell by projecting cells along the inferred regional continuum and rescaling values to the range [0,1], following the known anatomical order of regions. Cells were ordered by this index and grouped into equally sized bins. Mean RNA expression and histone modification signals were computed for each bin.

Axis-associated genes and histone modification regions were identified by testing for monotonic signal changes along the regional index, with multiple-testing correction. Histone regions were linked to genes using transcription start site windows (±2 kb). For interpretation, activating H3K27ac signals were required to change in the same direction as RNA expression, whereas repressive H3K27me3 and H3K9me3 signals were required to change in the opposite direction. Genes supported by both RNA and chromatin trends were used for downstream analyses. To visualize regional signal patterns, STR D1 and D2 cells were aggregated into region-specific pseudo-bulk profiles, and representative loci were displayed in the Integrative Genomics Viewer (IGV)^116^.

### Trajectory inference of the oligodendrocyte lineage

To infer the oligodendrocyte maturation trajectory, we first performed trajectory learning on a downsampled subset of the RNA data using Monocle3^117^. For downsampling, at most 2,000 cells were retained for each cell type, which limited bias from highly abundant populations while preserving the major lineage structure. Cells were selected to represent all oligodendrocyte stages, including OPC, COP, ImOligo, and mature oligodendrocyte populations.

Monocle3 was applied to this reduced dataset to learn the global lineage structure and to determine the overall ordering of oligodendrocyte stages. The inferred trajectory was consistent with the known biological progression of the oligodendrocyte lineage and was used to define a stage-ordered backbone.

Based on this backbone, we then projected all oligodendrocyte cells from the full dataset onto the trajectory in UMAP space. Briefly, a k-nearest neighbor graph was constructed using UMAP coordinates, and representative anchor cells were selected for each stage. Adjacent stage anchors were connected by shortest paths on the graph to form a continuous backbone. Each cell was assigned a pseudotime value by projecting its UMAP coordinate onto the backbone and computing its relative arc-length position, which was rescaled to the range [0, 1]. This pseudotime was used for downstream analyses of gene expression and chromatin dynamics along the oligodendrocyte maturation axis.

### TF motif enrichment and GO analysis

TF motif enrichment was performed for each cCRE module using HOMER^118^ (v5.1). The findMotifsGenome.pl function was applied with default parameters to identify enriched motifs relative to genomic background. TF motif enrichment results from the known motif database were used to generate the motif heat. Gene Ontology (GO) enrichment analysis was performed using gseapy^119^, a Python interface to the Enrichr platform, using differentially expressed genes from one-versus-rest comparisons as input. Enrichment was assessed against the GO Biological Process 2023^120^, KEGG^121^,and DisGeNET^122^ database, and significance was determined by Fisher’s exact test with Benjamini–Hochberg correction (adjusted *P* < 0.05).

### Cross-species integration of human and mouse single-nucleus RNA-seq

Human and mouse single-nucleus RNA-seq data were integrated using scVI-tools^123^ (scVI v0.20.3). Raw UMI counts were used as model input and stored in the counts layer. Cells from both species were combined into a single AnnData object, and species was provided as the batch covariate. An scVI model was trained on the combined dataset to learn a shared latent space that captures biological variation while reducing technical and species-associated effects. The trained scVI model was then used to initialize a semi-supervised SCANVI model. Human cells with known cell-type labels were treated as labeled data, and mouse cells were treated as unlabeled.

After training, a latent representation was computed for all cells and used for downstream graph construction and visualization. To further reduce residual species effects in the latent space, Harmony was applied to the SCANVI latent embedding using species as the batch variable. A k-nearest neighbor graph was constructed from the Harmony-corrected embedding (cosine distance; n_neighbors = 25), and UMAP coordinates were computed for visualization (min_dist = 0.25). Mouse cell-type labels were predicted using SCANVI, and prediction confidence was defined as the maximum class probability from the soft prediction output. Mouse predictions with confidence < 0.60 were assigned as “Unassigned”. The final integrated dataset was used for downstream clustering, visualization, and cross-species comparative analyses.

### Cross-species analysis of conserved and divergent chromatin peaks

To link mouse Paired-Tag chromatin profiles with RNA-defined cell groups, Paired-Tag nuclei were first matched to RNA cell identities using shared cell IDs. Nuclei were grouped by RNA-predicted cell type, and reproducible peak sets for each histone mark were obtained from IDR-filtered peak calls. For cross-species comparison, human peaks (hg38) were mapped to the mouse genome (mm10) using a liftOver-based approach. Each human peak was represented by its genomic center, and the mapped center position in mm10 was used to reconstruct a peak of the same length as the original human peak. Human peaks that could not be mapped were classified as human-specific peak.

Mapped human peaks were then compared with the corresponding mouse peak sets for the same cell type and histone mark. A mapped human peak was defined as His-conserved if it overlapped at least one mouse peak in mm10 coordinates. In contrast, a mapped human peak was defined as His-divergent if it showed no overlap with any mouse peak. Overlap tests were performed using BEDtools^112^. For each cell type and histone mark, human-specific, His-divergent, and His-conserved peaks were counted and summarized. These classifications were used to quantify the extent of chromatin conservation and divergence across species and to support downstream comparative analyses and visualization.

### Sample preparation for MERFISH 2.0

Fixed, photobleached, and permeabilized sections were washed twice in 15 mL Conditioning Buffer (Vizgen 20300116) in a 10 cm petri dish. Sections were then incubated in 10 mL Conditioning Buffer with 10 μL RNase inhibitor (New England BioLabs M0314L) at 37°C for 15 min. Anchoring pretreatment was performed by adding 200 μL Pre-Anchoring Reaction Buffer directly to the sample, covering with parafilm, and incubating at room temperature for 16 hours. After pretreatment, sections were washed with 10 mL Sample Prep Wash Buffer (Vizgen 20300001) and incubated in Formamide Wash Buffer (Vizgen 20300002) at 37°C for 15 min. RNA anchoring was carried out in Anchoring Buffer (Vizgen 20300117) at 37°C for 2 hours. Sections were then gel-embedded using Gel Embedding Premix (Vizgen 20300004) with 10% Ammonium Persulfate (Sigma 09913) and TEMED (Sigma T7024). 200 μL of gel embedding solution was applied, and Gel Slick (Lonza 50640)-treated coverslips (Corning 2855-25) were placed onto the sections with excess gel aspirated. Gel was allowed to polymerize for 1.5 hours. Following embedding, cell clearing was performed by incubating sections in Clearing Solution (Vizgen 20300114) and Proteinase K (New England Biolabs P8107S) 1:100 for 20 hours at 37°C in a humidified chamber. Sections were then washed 3x5 minutes in Sample Prep Wash Buffer and incubated in Formamide Wash Buffer at 37°C for 30 min. For probe hybridization, 200 μL of MERSCOPE 1000 Gene Panel, V 2.0 was applied directly on the sample and hybridized for 24 hours at 47°C in a humidified chamber. Post-hybridization washes were performed twice in Formamide Wash Buffer at 47°C for 30 min each. 200 μL of Enhancer Probes (Vizgen 20300194) were then applied to sections and covered with parafilm for a 20-hour hybridization at 37°C. The sample was washed twice with Enhancer Wash Buffer (Vizgen 20300192) at 37°C for 20 min, stained with DAPI and Poly T Reagent V2.0 for 15 min at room temperature, washed for 10 min with 5 mL of Formamide Wash Buffer, and imaged on the MERSCOPE Ultra system (Vizgen 10000008) using MERSCOPE Large 1000-Gene Imaging Kit V 2.0 (Vizgen 10400174). After image acquisition, MERFISH data were processed by MERSCOPE, and cell segmentation was performed using the Cellpose algorithm based on DAPI and PolyT staining. A fully detailed, step-by-step instruction on the MERFISH sample prep can be found in Vizgen’s MERFISH 2.0 Sample Preparation User Guide for Sectioned Tissue Samples (Vizgen 91600132). Full Instrumentation protocol can be found in MERSCOPE Instrument Guide (Vizgen 9160001). The MERFISH dataset was generated as two replicates by independent groups in the Bing Ren lab and the Joseph R. Ecker lab on adjacent tissue slices, and both replicates were included in downstream analyses.

### MERFISH gene panel selection

We selected 920 genes for MERFISH imaging on the Vizgen MERSCOPE Ultra platform. Two initial subsets of genes were selected based on the snRNA-seq DEGs using NS Forest^124^ and snm3C-seq DMGs using Spapros (https://github.com/theislab/spapros) separately, then combined those two gene lists with the top 3 DEGs and DMGs. After removing the duplicated genes, adding additional cell type marker genes and genes of interest, and 20 internal control genes with predictable expression across cell types, we obtained 920 genes. We selected 20 internal control genes to facilitate trouble shooting and quality comparison across experiments. These genes were selected based on the Siletti et al. scRNA-seq dataset^23^ by identifying the top 20 genes with the least variance of gene expression across all brain cell types and were expressed in at least 10,000 brain cells. Finally, Vizgen eliminated genes that were unsuitable for MERFISH, and we added replacement genes for those eliminated. In order to confirm the viability of the gene list for cell type identification, we re-clustered our Droplet Paired-tag data-subset to the genes in our panel-to confirm that the original clusters could still be distinguished. Code library and probe design were optimized by Vizgen; see Table S1.2 for the complete codebook and gene list.

### MERSCOPE data analysis and quality control

MERSCOPE’s built in pipeline was used to call transcripts^23^, and starting cell boundaries were identified using Cellpose 1^125^ based on the DAPI staining. Transcripts were assigned to original cells identified by Cellpose to create a cell-by-gene table. Proseg^126^ was used to assign additional transcripts that were excluded due to conservative DAPI boundaries. Cells were filtered to remove MERFISH cells with counts less than 10 counts per gene, as well as cells with volume greater than three times the median, similar to methods described previously^127,128^. We evaluated the overall MERFISH data by quantifying the detection efficiency of each dataset, the reproducibility among four donors and between technical replicates, and the expression correlation with bulk RNA-seq. We observed a generally high correlation between technical replicates and donors, measured by mean counts of mRNA molecules detected per imaging area and per cell. We also correlated the gene expression from MERFISH data with bulk RNA-seq data and got reasonable correlations.

### Clustering and reference-based annotation of MERFISH data

MERFISH datasets from multiple preprocessed imaging slides were pooled for joint clustering and cell-type annotation using Seurat (v5.2.0)^129^. Spatial information was incorporated by constructing field-of-view (FOV) objects based on cell centroid coordinates for each imaging slide independently. Imaging slides that failed to capture expected laminar tissue organization, as assessed by biochemical staining and tissue integrity metrics, were excluded from downstream analyses.

To assign MERFISH cells to transcriptomic cell types, MERFISH gene expression profiles were integrated with a reference single-nucleus RNA-seq dataset generated from the same brain regions. Prior to integration, both MERFISH and snRNA-seq datasets were normalized using SCTransform (v0.4.2)^130^ to mitigate differences in transcript detection efficiency across assays. Integration was performed using canonical correlation analysis (CCA) to generate a shared low-dimensional embedding.

Cell-type annotation of MERFISH cells was carried out using a neighborhood-based probabilistic classifier operating in the joint CCA embedding space. For each MERFISH cell, the k = 50 nearest reference snRNA-seq cells were identified based on Euclidean distance. Distances between a MERFISH cell and its nearest neighbors were converted into normalized weights that sum to one^40^. A weighted probability score was computed for each candidate cell type, and each MERFISH cell was assigned to the cell type with the highest weighted probability. Assignments with a maximum probability below 0.4 were considered ambiguous and were labeled as unassigned.

### Spatial position imputation for Droplet Paired-Tag cells

To infer spatial positions for Droplet Paired-Tag cells, we performed reference-based spatial imputation using MERFISH data as a spatial anchor. Imputation was conducted separately for each imaging slide and cell subclass to preserve region- and subclass-specific spatial information.

First, Droplet Paired-Tag cells were first projected into a shared low-dimensional space with matched MERFISH cells using canonical correlation analysis (CCA) embeddings. For each Droplet Paired-Tag cell, a nearest-neighbor graph was constructed in the CCA space, restricting candidate reference cells to those originating from the same dissection region and subclass. The k = 10 nearest MERFISH reference cells were identified based on Euclidean distance. Next, spatial coordinates for each Droplet Paired-Tag cell were imputed by assigning the centroid coordinates of its closest MERFISH neighbor in the embedding space. This nearest-neighbor assignment preserves fine-scale spatial structure while minimizing over-smoothing across anatomical boundaries. To assess confidence in spatial imputation, we computed an unweighted neighborhood purity score for each Droplet Paired-Tag cell, which measures the label consistency across its top 10 nearest neighbors. Only Droplet Paired-Tag cells with a purity score ≥0.5, indicating that at least half of the nearest reference neighbors shared the same subclass identity, were retained for downstream spatial visualization.

### GWAS heritability enrichment analysis

To assess enrichment of GWAS heritability in cell-type-specific chromatin features, we performed partitioned linkage disequilibrium score regression (LDSC) (v1.0.1) using histone modification peaks^131^. For each cell type (subclass and group) and histone mark, foreground annotations were defined by peaks called in that cell type. To control for state-specific genomic properties, matched background annotations were generated by merging all peaks called using the same histone mark across all cell types.

All foreground and background peak sets were converted from hg38 to hg19 using a reciprocal liftOver strategy. Briefly, cCREs were lifted from hg38 to hg19 and then back to hg38 using a minimum match threshold of 0.95, and only regions that successfully mapped in both directions were retained. To further minimize mapping artifacts, cCRE widths were capped at 1 kb in hg19 prior to liftOver. This procedure ensured that only high-confidence regulatory regions were included in downstream analyses.

GWAS summary statistics for neurological and control traits were obtained from publicly available sources and converted into LDSC-compatible annotation files using make_annot.py. LD scores were computed with a 1 cM LD window and restricted to HapMap3 SNPs to ensure robust LD estimation^132^. LD score computation was performed independently for each cell-type- and histone modification-specific annotation, as well as for the corresponding matched background annotation, using 1000 Genomes European reference panels^133^. Partitioned LDSC regression was performed using the ‘--h2-cts’ framework. For each histone modification, a partition definition file was constructed to jointly model all cell-type-specific annotations alongside the matched background annotation. LDSC regression was run for each GWAS trait using baseline LD annotations and standard regression weights. For each trait, statistical significance was assessed after correcting for multiple hypothesis testing using the Benjamini-Hochberg (BH) procedure.

### Fine-mapping

We conducted fine-mapping using FINEMAP with a similar pipeline adapted from the FinnGen study^134,135^. Specifically, for each genome-wide significant locus (default configuration: P < 5e-8), we define a fine-mapping region by taking a 3 Mb window around a lead variant. We merge the regions if they overlap. If a merged window exceeds 6MB, we iteratively shrink the window by 10%, until the merged window fits into 6MB or is split into merged windows that each fit into 6MB. We then calculate the LD using plink for each fine-mapping region with reference from 1000 Genome Project^133,136^. With the inputs of summary statistics and LD from the steps 1-2, we conduct fine-mapping using FINEMAP with the maximum number of causal variants in a locus L = 10.

### Identify putative target genes for fine-mapped non-coding variants

We intersected activity-by-contact (ABC) enhancer-gene links with fine-mapped neuropsychiatric disease variants to identify putative downstream target genes of non-coding risk loci. ABC links were filtered to retain only subclass-resolved interactions involving promoter-distal enhancers, thereby focusing the analysis on cell-type-specific regulatory elements. Fine-mapped variants were converted from hg19 to hg38 coordinates using UCSC liftOver^137^, and genomic intersections between variants and ABC-defined enhancer regions were performed using the GenomicRanges package (v1.58.0) in R^138^. Variants overlapping ABC enhancers were assigned to their linked target genes based on the corresponding links. The resulting variant-enhancer-gene pairs were subsequently used to prioritize loci for downstream visualization and interpretation.

### Deep learning model architecture

Our deep learning model utilizes a hybrid architecture integrating Convolutional Neural Networks (CNNs) and Transformer networks. The backbone follows the Borzoi^93^ architecture, comprising convolutional and transformer blocks, followed by a task-specific prediction head optimized for Droplet Paired-Tag data. Specifically, we introduced a subset of cell-type-specific embedding heads designed to project the sequence embeddings (dimension: 1024), generated by the CNN up-sampling layers, into an 8-dimensional cell-type-specific chromatin embedding space. Subsequently, a subset of shared chromatin profile heads decodes these embeddings into corresponding genomic tracks, including chromatin accessibility (ATAC), three histone modifications (H3K27ac, H3K27me3, H3K9me3), and two gene expression tracks (forward and reverse strand RNA).

Unlike the default Borzoi configuration, this architecture decouples cell-type specificity from track decoding. This design improves efficiency for Droplet Paired-Tag data by enabling the model to explicitly learn correlations between tracks within a single cell type. Each cell-type-specific embedding head consists of a linear layer (1024x8) without activation. We observed that adding non-linearity to this stage yielded comparable performance, favoring the simpler linear approach. Each chromatin profile head consists of a linear layer (8x6) followed by a Softplus activation function. We ablated various hidden dimension sizes and determined that an 8-dimensional embedding yields optimal performance.

### Deep learning model training and testing datasets

Training data were preprocessed from BAM files. We utilized bamCoverage to convert BAM files into BigWig format. To enhance signal stability, we applied data smoothing to the histone modification tracks. Specifically, we used a smoothing window of 300 bp for H3K27ac and H3K27me3, aligning with the approximate DNA wrapping length of a nucleosome (∼150 bp). For H3K9me3, we applied a smoothing window of 1000 bp to capture its broader signal domains. RNA data were processed from BAM to BigWig format following the ENCODE protocol^139,140^. We processed data from 70 distinct cell types across the basal ganglia (n=36) and cortex (n=34). For the final dataset, we retained 37 cell types with >1000 cells (14 basal ganglia and 23 cortex cell types), the cell type names and modalities are available in the (Table S7.1)

Genome-wide train/validation/test splits were conducted following the Borzoi protocol, including identical data augmentation strategies (reverse complement and random shifting by ±3 bp). We applied optimal clipping and soft clipping to the Droplet Paired-Tag signals to mitigate noise-induced artifacts; specific thresholds are detailed in the data configuration file available on (Table S7.1).

### Deep learning model training

We initialized the model backbone with pre-trained weights from Borzoi^93^, while the cell-type embedding heads and chromatin profile heads were randomly initialized. The model was trained using the Adam^141^ optimizer with an initial learning rate of 1x10^(-4). The schedule included a warmup phase of 625 steps, followed by a linear decay over 25,000 steps to a minimum learning rate of 1x10^(-5). Training was distributed across 8 NVIDIA A40 GPUs with a batch size of 1 per GPU. Training ran for a maximum of 200 epochs, utilizing early stopping based on validation loss convergence. A complete training run converged at around 25 epochs and required approximately 10 hours.

We trained three distinct model variants for comparison:

1. Main Model (Full Model): Trained on the filtered set of 37 cell types (>1000 cells).
2. Ablation Model (ATAC-only): Trained on the same cell types but restricted to ATAC and RNA modalities.
3. Full Cell-Type Model: Trained on all 70 cell types to evaluate performance scaling relative to cell population size.

### Deep learning model evaluation

We evaluated model performance using two distinct validation schemes to assess both local signal fidelity and cell-type specificity. In the first scheme, we measured the intra-cell type Pearson correlation between observed and predicted signals within the testing dataset. This assessment was conducted at two resolutions: a high-resolution 32 bp bin level, where read counts were aggregated into 32 bp genomic windows, and a gene-level resolution, where signals were aggregated across all exons associated with each gene using GENCODE v48 annotations^142^. In the second scheme, we evaluated the inter-cell type Pearson correlation to determine the model’s ability to distinguish signal variation across different cell types. This analysis was performed at the level of cCREs, where read counts were aggregated within 500 bp consensus regulatory elements defined by ATAC-seq peaks, and at the gene level, where read counts were aggregated across all exons for each gene.

### Motif Discovery

To identify sequence motifs that drive cell-type-specific gene expression, we computed sequence attribution scores using the Gradient × Input method^93^. For each gene of interest, we defined the prediction target as the aggregated read coverage across all exons associated with the gene in a specific cell type. We then calculated the gradient of this target with respect to the input sequence at each position *j* for all four nucleotides, *S_i,j_*. For visualization purposes, we extracted the attribution scores corresponding solely to the reference nucleotides, *S_ref,j_*. To generate a Position Weight Matrix (PWM), we applied a SoftMax transformation to the attribution scores across all four nucleotides at each position (Equation X). Finally, we utilized TomTom to match the resulting PWMs against the HOCOMOCO v12 motif database^143^.

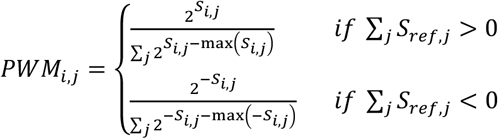

### Variant effect prediction

We applied the model to predict variant effects using a cell-type and modality-specific approach. For a given locus, inputs for both the wild-type allele and its reverse complement were averaged to generate the wild-type prediction. This process was repeated for the mutant allele. We quantified the variant effect by calculating the difference between wild-type and mutant predictions using the L2 score metric described in the Borzoi^93^ paper.

To benchmark performance, we calculated the L2 sum of L2 scores across all tracks for both Borzoi and our model as the average of both models. We generated a dataset of causal eQTL variants (finemap posterior inclusion probability (PIP) > 0.9) together with negative controls (finemap PIP < 0.0001) in GTEx^144^ basal ganglia tissues to evaluate both scores. For both scores, we computed the Area Under the Receiver Operating Characteristic (AUROC) and did bootstrapping 100 times to estimate the standard error. Finally, to benchmark the performance of prioritizing causal variants in schizophrenia GWAS results, we employed stratified Linkage Disequilibrium Score Regression (S-LDSC)^131^. We calculated the heritability enrichment of the top 5% of variants ranked by our model per cell type and compared this against the top 5% of variants ranked by average score of Borzoi.

To benchmark the variant effect sizes on gene expression change, we curated two datasubsets of eQTL variants. The first dataset is generated from Jang *et al.*^97^, where we took the finemapped variants of cell types that overlapped with our basal ganglia data. For each finemapped variant, we calculated the log fold change of aggregated read counts across all exons for the corresponding gene and cell type RNA track. The second dataset is generated by N. de Klein *et al.*^96^, where we took the finemapped variants of basal ganglia region. For each fine-mapped variant, we calculated the log fold change of aggregated read counts across all exons and all cell type RNA tracks for the corresponding gene.

