## Supplementary material for "Single-cell Multiome Analysis of Chromatin State and Transcriptome in the Human Basal Ganglia": Sup figures: Sup figures-2601131.pdf

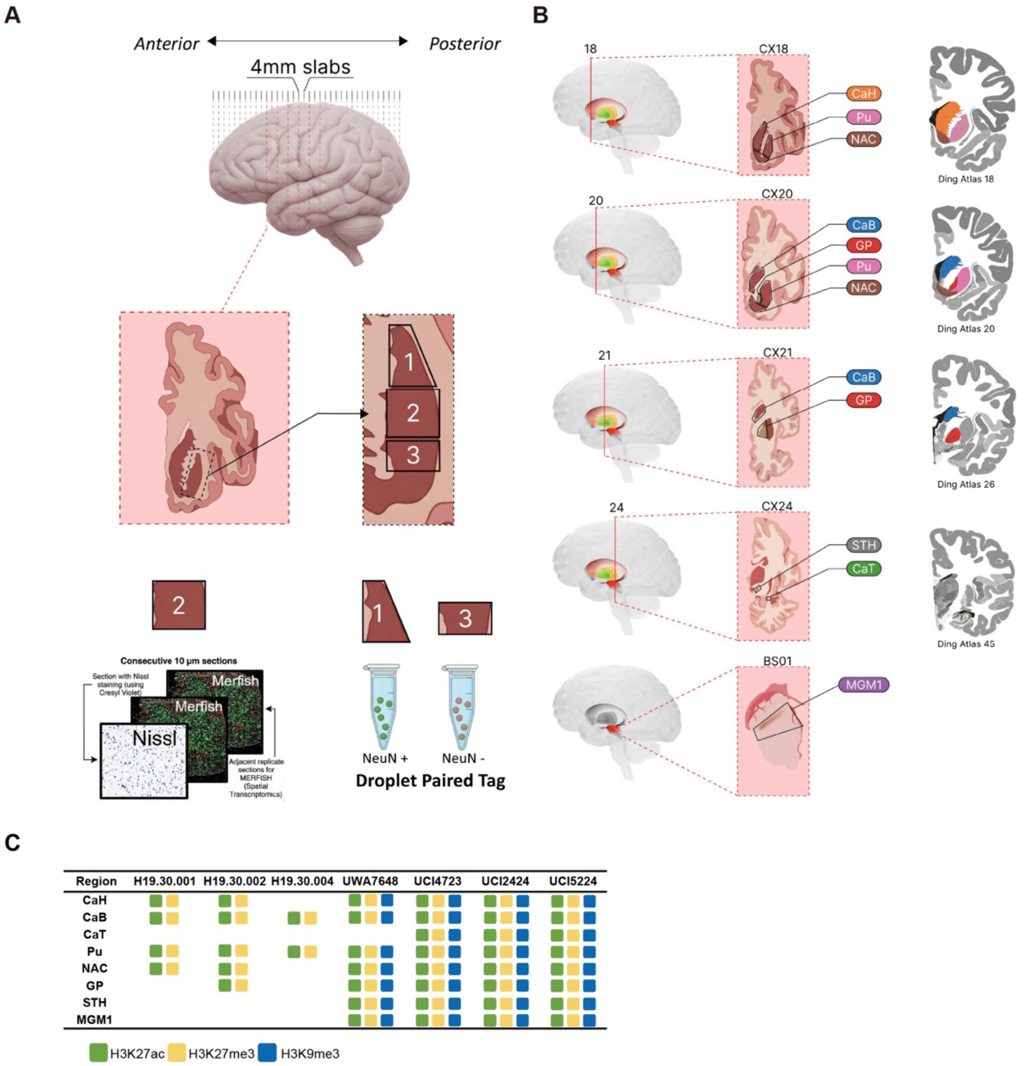


**Figure S1.1 Tissue dissection and multimodal profiling strategy for each human basal ganglia region.**

(A) Schematic of tissue dissections strategy for Droplet Paired-Tag and MERFISH.

(B) Schematic illustrating the dissection boundaries for each basal ganglia region.

(C) Table showing Droplet Paired-Tag data coverage for each donor and each region.

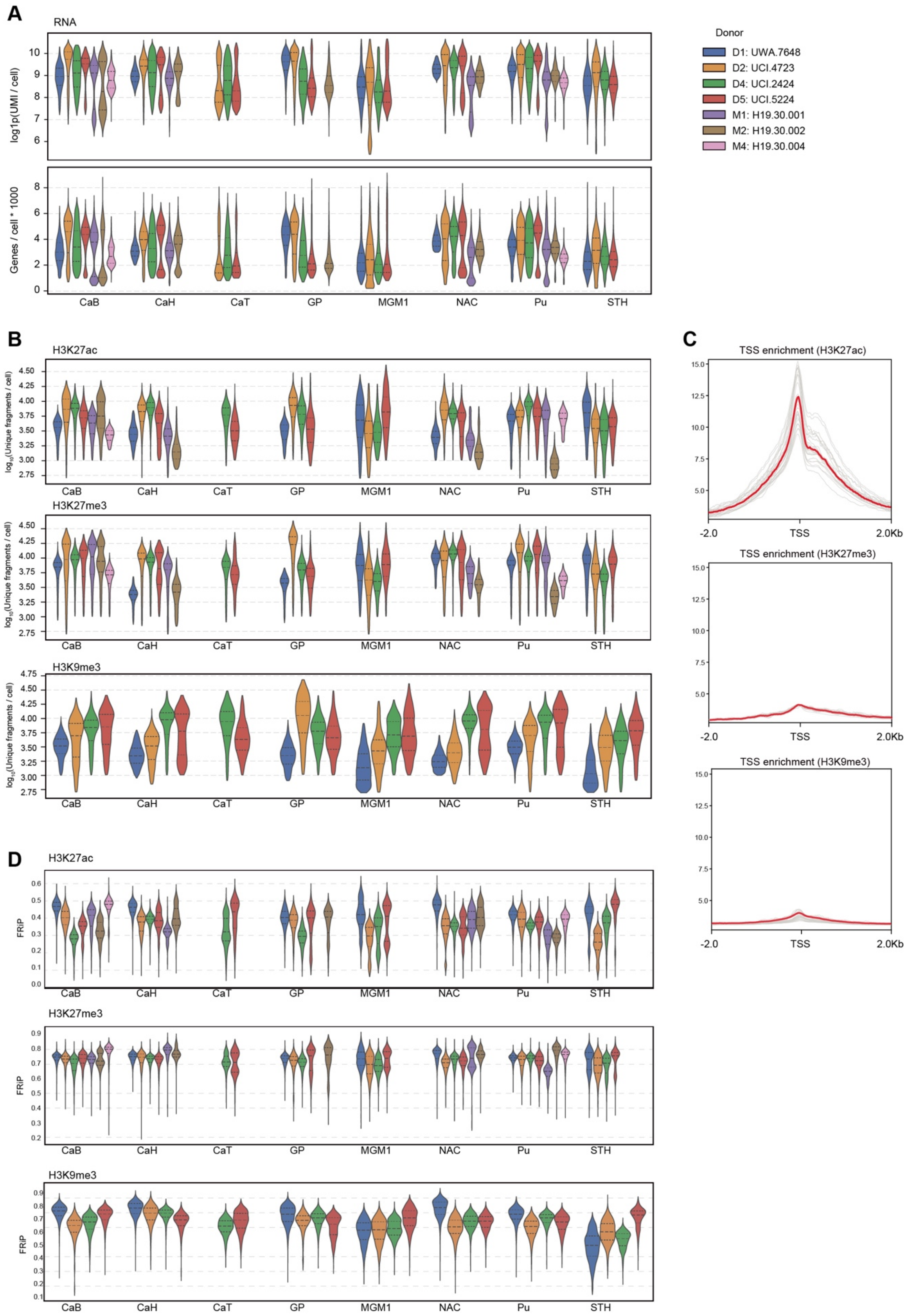


**Figure S1.2 Droplet Paired-Tag quality control metrics.**

(A) Box plots showing the distributions of the UMI and genes number per cell in Droplet Paired-Tag RNA modality separately for each donor and region.
(B) Box plots showing the distributions of the unique chromatin fragments number per cell detected by H3K27ac, H3K27me3 and H3K9me3 antibodies in Droplet Paired-Tag DNA modality separately for each donor and region.
(C) Box plots showing the distributions of FRiP for H3K27ac, H3K27me3 and H3K9me3 profiles for each donor and region.
(D) Distribution of TSS enrichment for H3K27ac, H3K27me3 and H3K9me3 profiles. TSSe curves for each donor and region are shown in grey; average TSSe curves are shown in red.

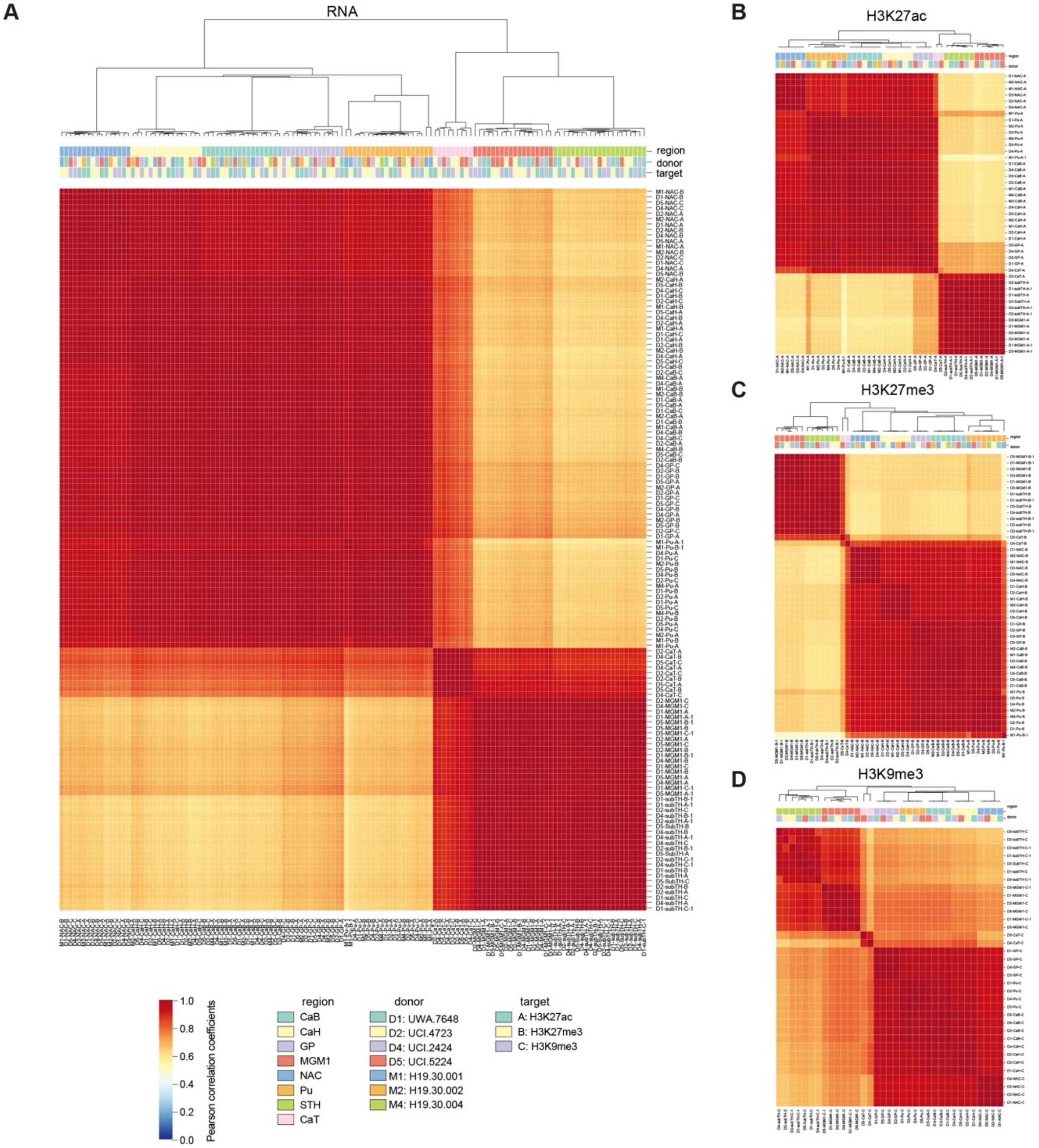


**Figure S1.3 Droplet Paired-Tag library correlation matrix.**
(A) Heat map showing the pairwise Pearson correlation coefficients between Droplet Paired-Tag RNA libraries. The row names consist of three parts: donor label, basal ganglia region and histone modification mark.

(B-C) Heat map showing the pairwise Pearson correlation coefficients between Droplet Paired-Tag DNA libraries, including H3K27ac (B), H3K27me3 (C), and H3K9me3 (D).

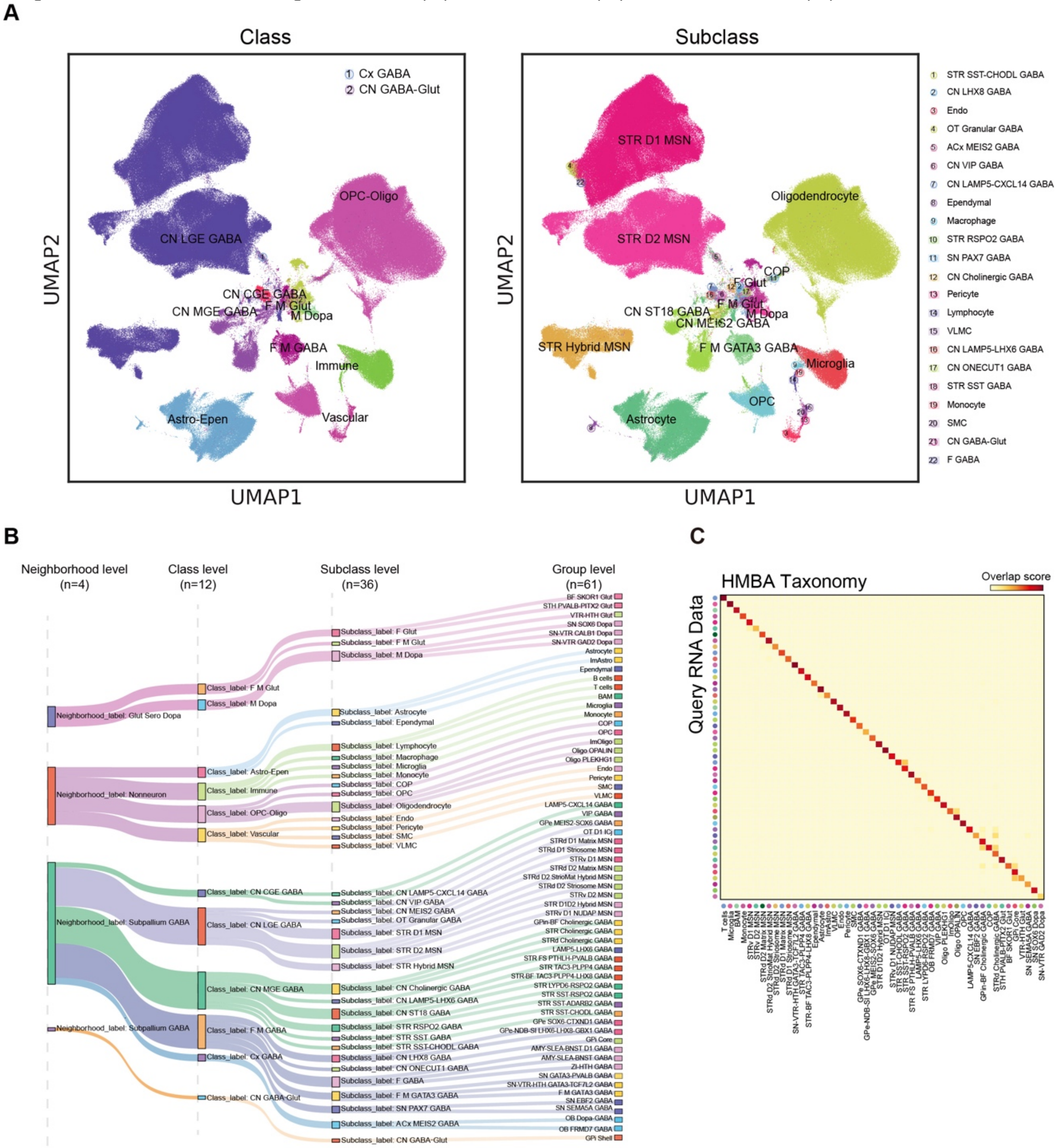


**Figure S1.4 Cell type annotation for Droplet Paired-Tag.**

(A) UMAP embedding and clustering analysis of the transcriptome modality from Droplet Paired-Tag dataset, colored by class types (left) and subclass types (right).

(B) Sankey diagram showing the hierarchical cell-type taxonomy derived from Droplet Paired-Tag data. Cells were first assigned to neighborhoods (n = 4), and then progressively refined into classes (n = 12), subclasses (n = 36), and groups (n = 61). Box heights are proportional to the number of cells at the neighborhood, class, and subclass levels, and ribbons trace how each cell population at a coarser level subdivides into finer annotations.

(C) Confusion matrix showing Droplet Paired-Tag query cells and HMBA cells overlap scores at each group (cell number>50) after integration. Droplet Paired-Tag query cells were annotated by MapMyCell hierarchical correlation mapping algorithm with the HMBA taxonomy as reference.

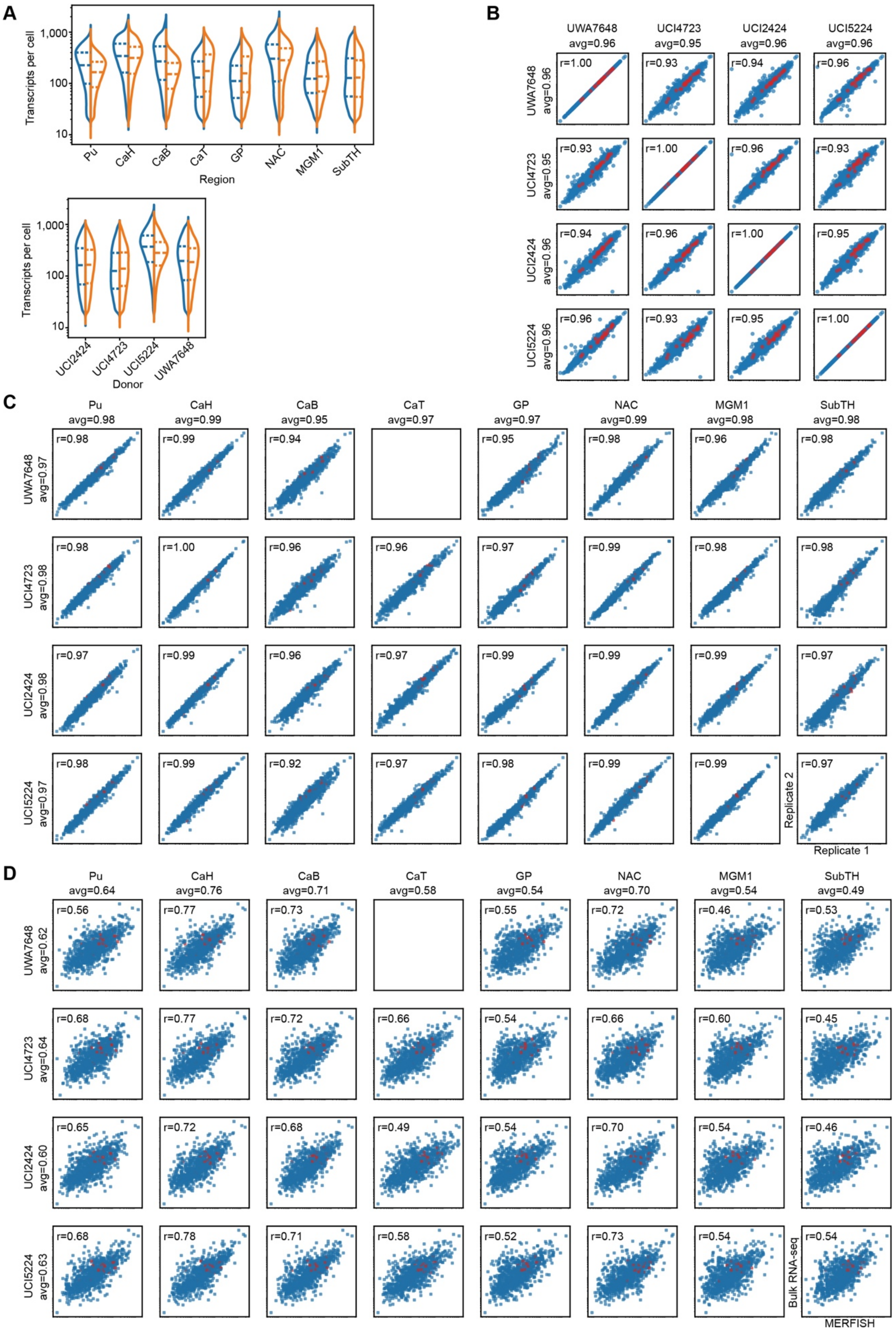
A
Transcripts per cell
1,000
100
10
Pu
CaH
CaB
CaT
GP
NAC
MGM1
SubTH
Region
Transcripts per cell
1,000
100
10
UCI2424
UCI4723
UCI5224
UWA7648
Donor
B
UWA7648 avg=0.96
UCI4723 avg=0.95
UCI2424 avg=0.96
UCI5224 avg=0.96
UWA7648 avg=0.96
r=1.00
r=0.93
r=0.94
r=0.96
UCI4723 avg=0.95
r=0.93
r=1.00
r=0.96
r=0.93
UCI2424 avg=0.96
r=0.94
r=0.96
r=1.00
r=0.95
UCI5224 avg=0.96
r=0.96
r=0.93
r=0.95
r=1.00
C
Pu avg=0.98
CaH avg=0.99
CaB avg=0.95
CaT avg=0.97
GP avg=0.97
NAC avg=0.99
MGM1 avg=0.98
SubTH avg=0.98
UWA7648 avg=0.97
r=0.98
r=0.99
r=0.94
r=0.95
r=0.98
r=0.96
r=0.98
UCI4723 avg=0.98
r=0.98
r=1.00
r=0.96
r=0.96
r=0.97
r=0.99
r=0.98
r=0.98
UCI2424 avg=0.98
r=0.97
r=0.99
r=0.96
r=0.97
r=0.99
r=0.99
r=0.99
r=0.97
UCI5224 avg=0.97
r=0.98
r=0.99
r=0.92
r=0.97
r=0.98
r=0.99
r=0.99
r=0.97
Replicate 2
Replicate 1
D
Pu avg=0.64
CaH avg=0.76
CaB avg=0.71
CaT avg=0.58
GP avg=0.54
NAC avg=0.70
MGM1 avg=0.54
SubTH avg=0.49
UWA7648 avg=0.62
r=0.56
r=0.77
r=0.73
r=0.55
r=0.72
r=0.46
r=0.53
UCI4723 avg=0.64
r=0.68
r=0.77
r=0.72
r=0.66
r=0.54
r=0.66
r=0.60
r=0.45
UCI2424 avg=0.60
r=0.65
r=0.72
r=0.68
r=0.49
r=0.54
r=0.70
r=0.54
r=0.46
UCI5224 avg=0.63
r=0.68
r=0.78
r=0.71
r=0.58
r=0.52
r=0.73
r=0.54
r=0.54
Bulk RNA-seq
MERFISH

**Figure S1.5 MERFISH quality control metrics.**
(A) Violin plots showing the distribution of total transcripts per cell for each brain region (top) and for each donor (bottom). Two replicates from two different labs are shown in blue and orange, separately.
(B) Correlation of average gene expression of 920 MERFISH panel genes across donors. Red points represent internal control genes selected for their predictable expression profiles across cell types and samples.
(C) Correlation between two MEFISH replicates for each donor and brain region, computed using average transcripts per cell per gene. Pearson correlation coefficients and average values are shown for each replicate pair.
(D) Correlation between MERFISH average transcripts per cell per gene and GTEx bulk RNA seq raw counts for the Putamen. Pearson correlation coefficients are shown for each donor and region.

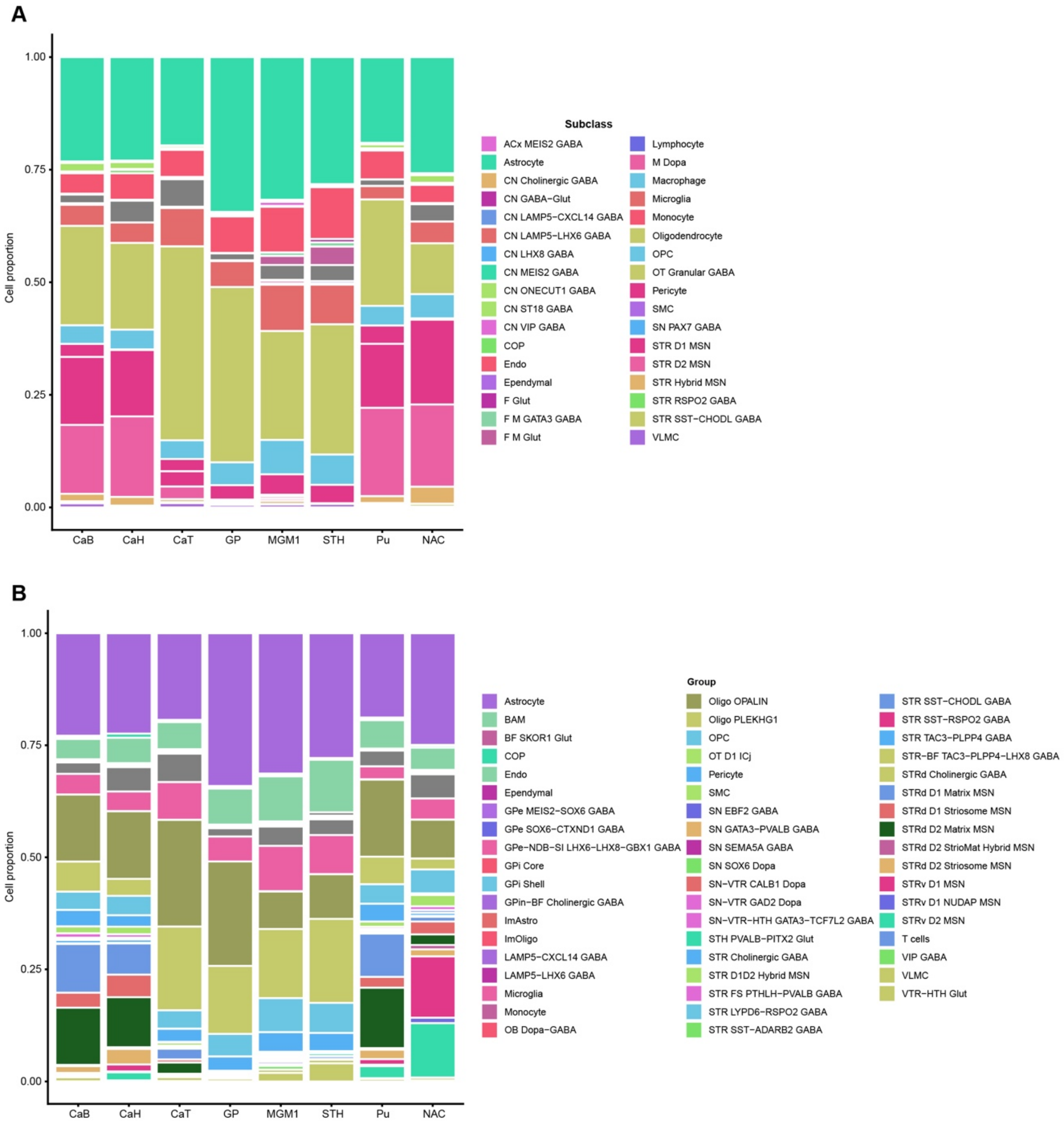


**Figure S1.6 Cellular composition for each basal ganglia region from MERFISH dataset.**
(A-B) Stacked bar plot showing the percentage of each cell subclass (A) and group (B) defined in MERFISH images from each basal ganglia region.

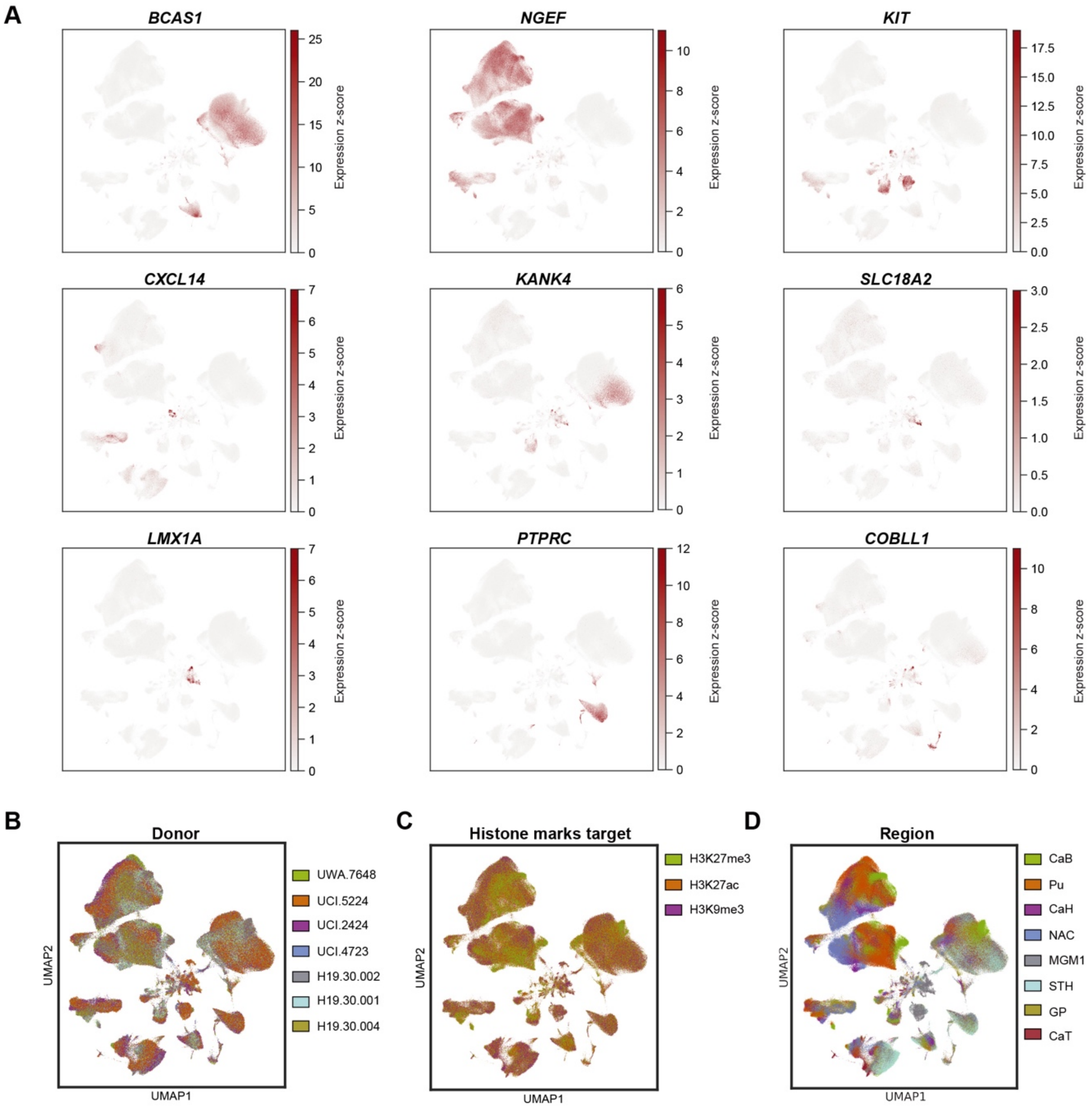


**Figure S1.7 UMAP visualization of Droplet Paired-Tag profiles.**
(A) UMAP of the Droplet Paired-Tag RNA modality colored with gene expression level of marker genes for each group.
(B-C) UMAP visualization of the Droplet Paired-Tag RNA modality showing donor (B), three histone modification marks (C), and dissected region distribution (D).

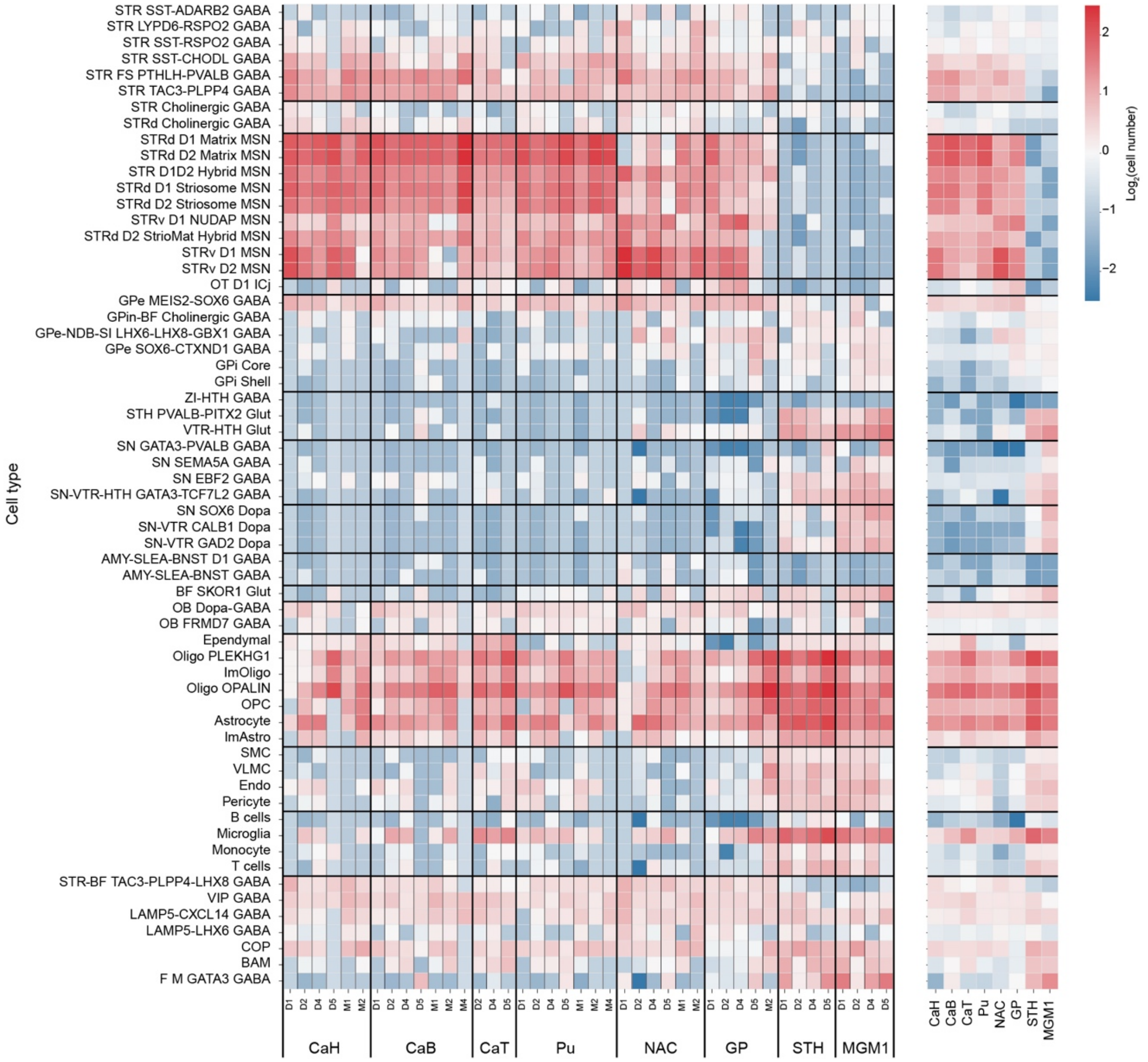


**Figure S1.8 Cellular composition for each basal ganglia region from Droplet Paired-Tag dataset.**

Heatmap showing the number of cells recovered in the Droplet Paired-Tag dataset for each cell group, divided by basal ganglia region and donor.

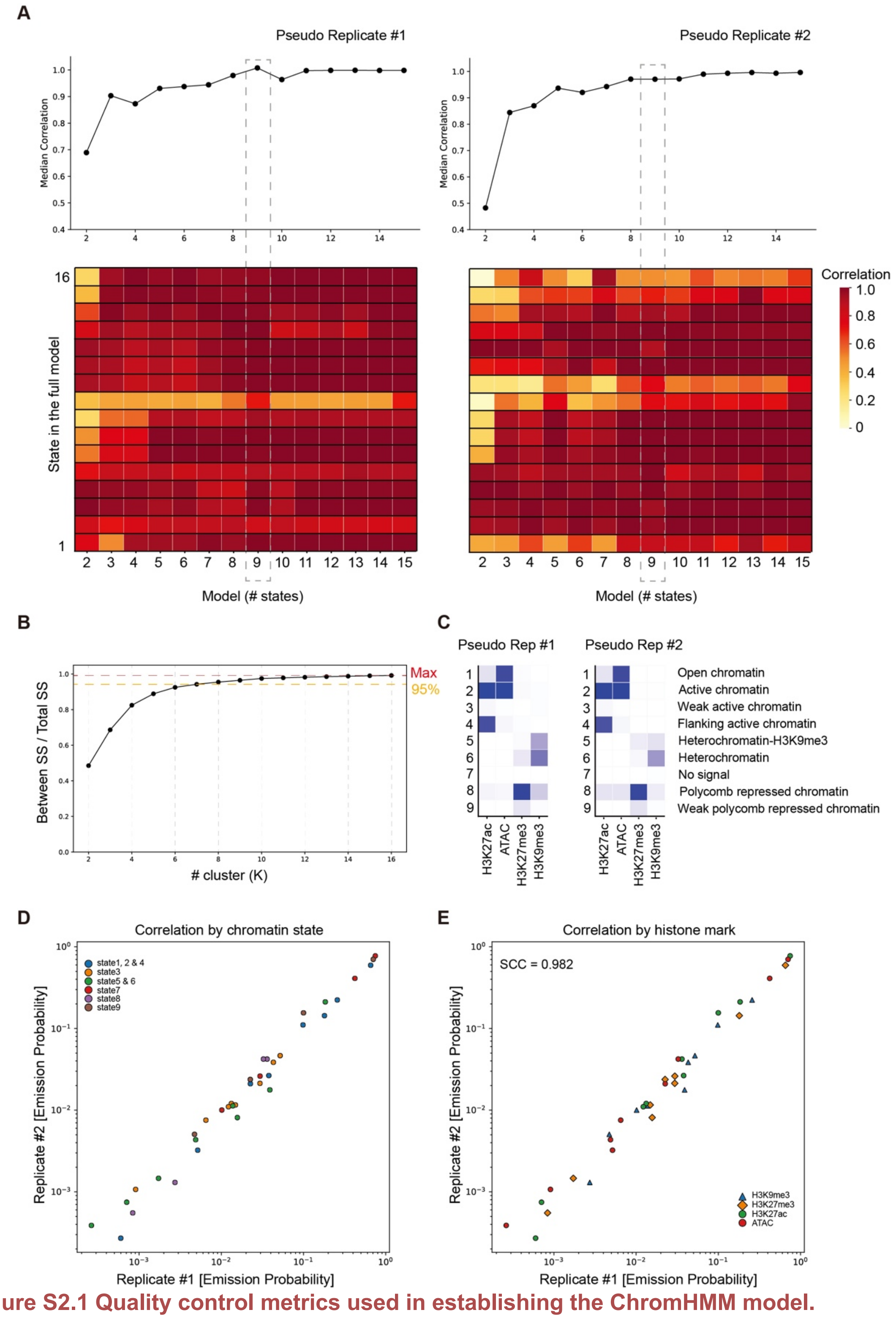


**Figure S2.1 Quality control metrics used in establishing the ChromHMM model.**

(A) Heatmaps showing the maximum Pearson's correlation between each state in the full model and its best-matching state in each simpler model in two pseudo-replicates. The top panels report the median correlation across all 15 states, for both replicates.

(B) K-means clustering of emission probabilities across all models. The optimal number of states was defined as the smallest k achieving at least 95% of the maximum cluster separation. SS, sum

of squares.
(C) ChromHMM-defined emission probabilities for three histone modification marks and chromatin accessibility across states, shown for both replicates.
(D and E) Spearman's correlation of ChromHMM emission probabilities between the two biological replicates, colored by states (D) and by chromatin mark (E).

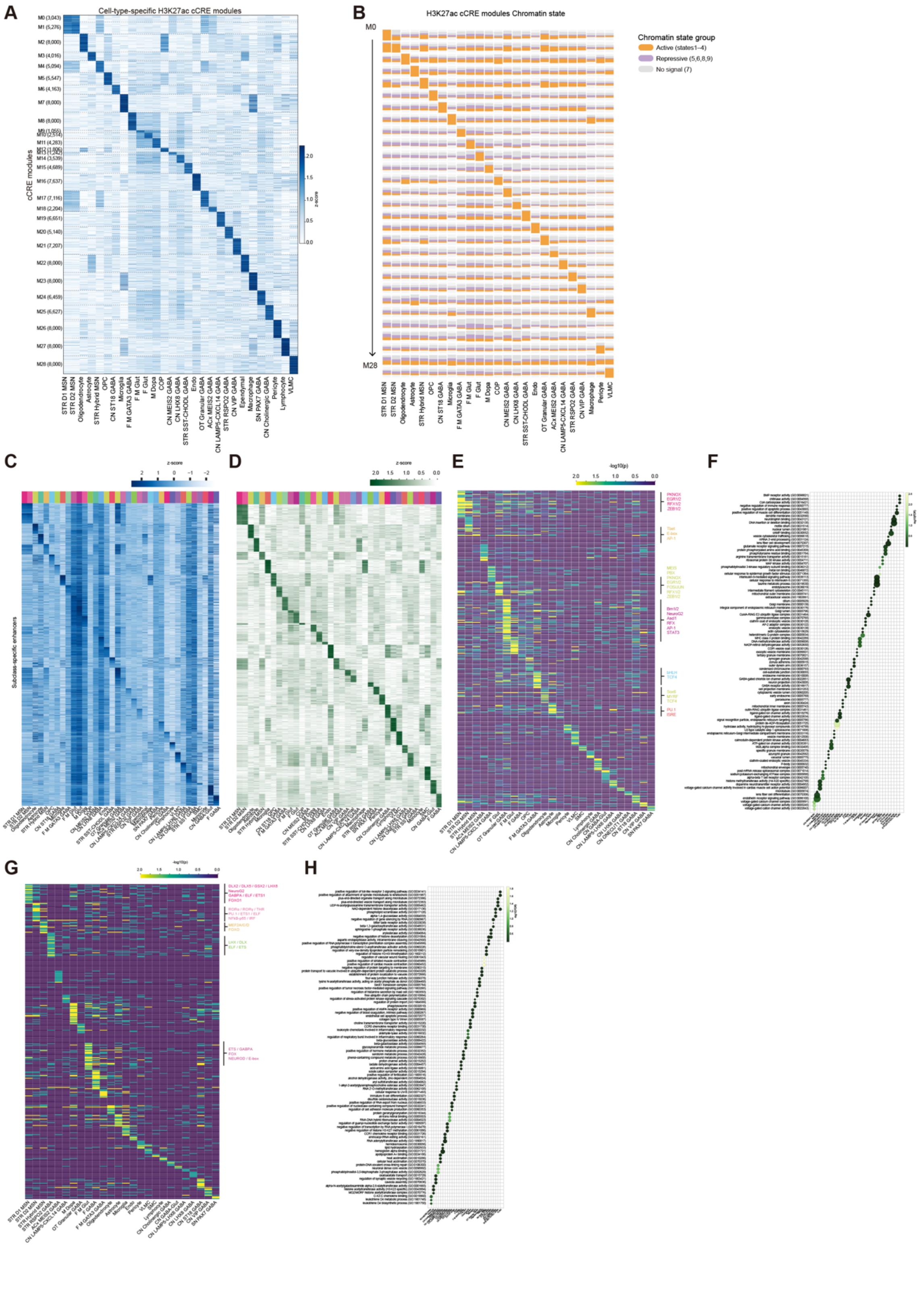

**Figure S2.2 Identification and characterization of cCREs across basal ganglia subclasses.**
(A) Clustering of subclass-specific cCREs into 29 modules (M0-28) based on H3K27ac profiles across different subclasses.
(B) Chromatin state of H3K27ac-marked cCREs in each module across cell subclasses.
(C) The H3K27ac levels of 36 subclass-specific enhancer modules identified from E-P loops.
(D) The RNA expression of enhancer linked genes within E-P loops across 36 subclasses.
(E) TF motif enrichment in enhancers identified from E-P loops across different subclasses.
(F) Biological pathways enriched in enhancer linked genes within E-P loops across 36 subclasses.
(G) TF motif enrichment in silencers identified from S-P loops across different subclasses.
(H) Biological pathways enriched in silencers linked genes within S-P loops across 36 subclasses.

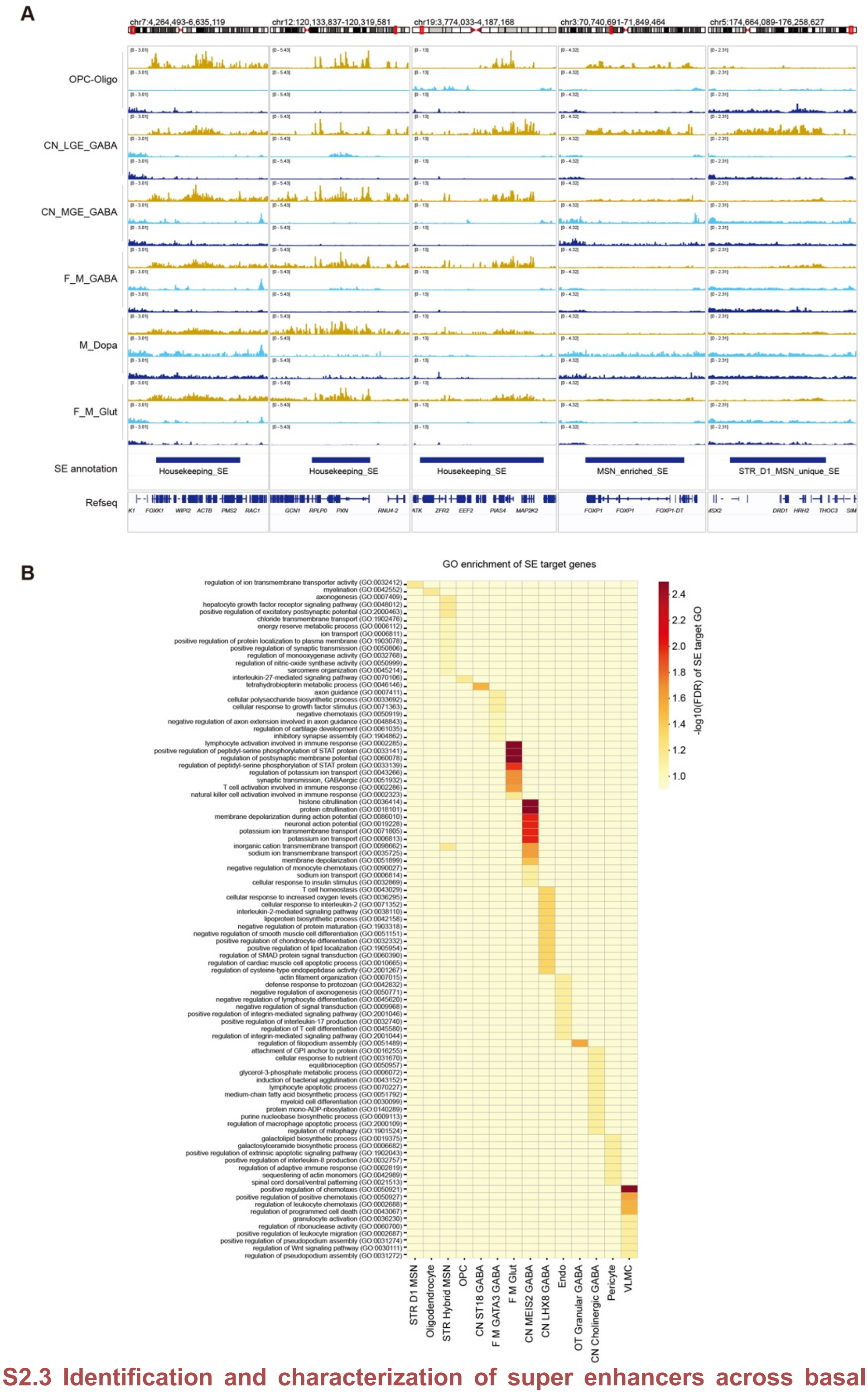


**Figure S2.3 Identification and characterization of super enhancers across basal ganglia subclasses.**

(A) Genome browser tracks of histone modification profiles for each class at housekeeping super enhancers shared across classes and at class- or subclass-specific super enhancers.

(B) GO enrichment analysis for target genes of super enhancers. Color represents enrichment significance (-$\log_{10}$ FDR).

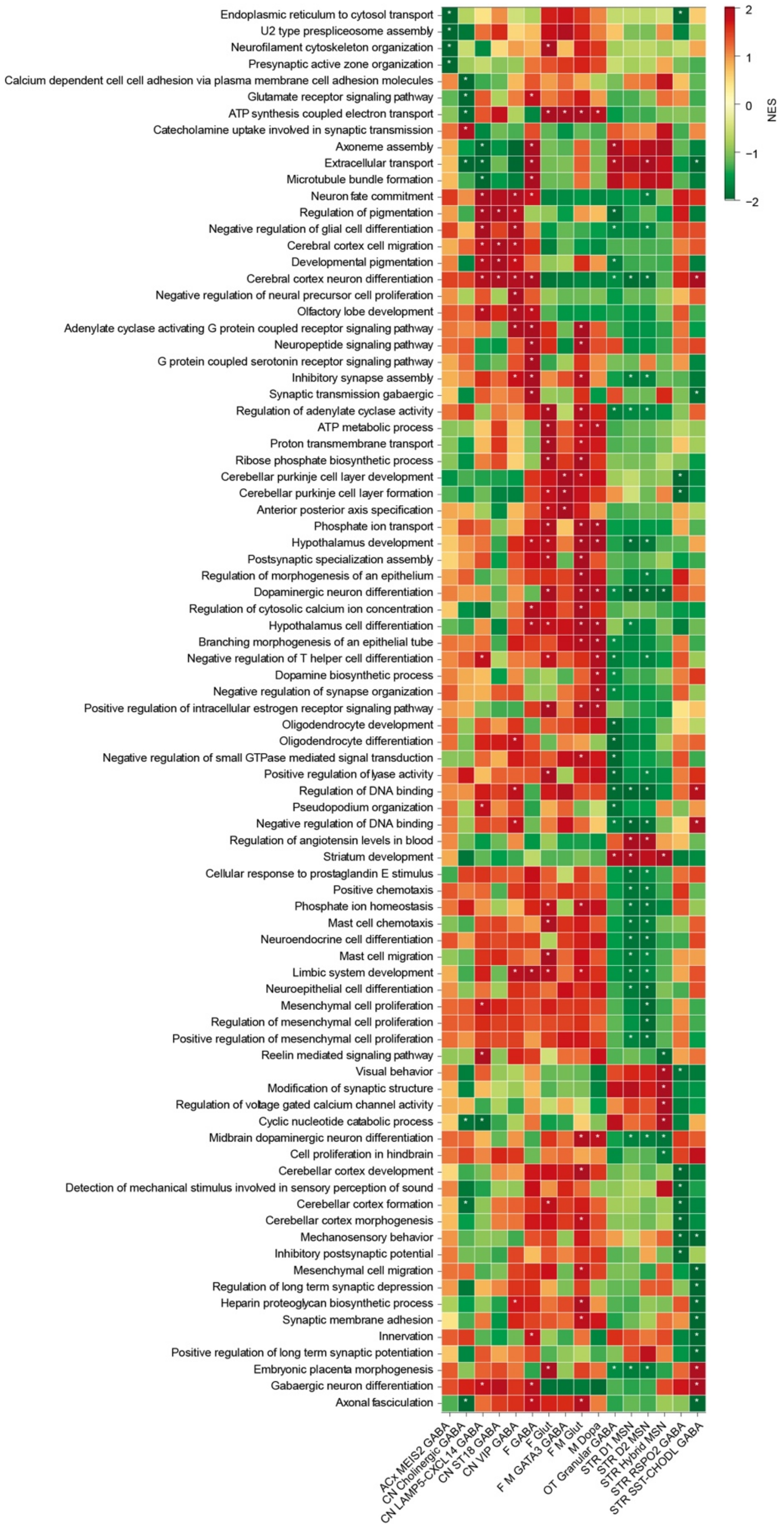


**Figure S3.1 Biological pathways enrichment analysis of DEGs across basal ganglia subclasses.**

Heatmap showing biological pathways enrichment for genes with significant up and down changes in each subclass. Color shown in the heatmap are normalized enrichment scores (NES). Asterisks present pathways significant at FDR < 0.1.

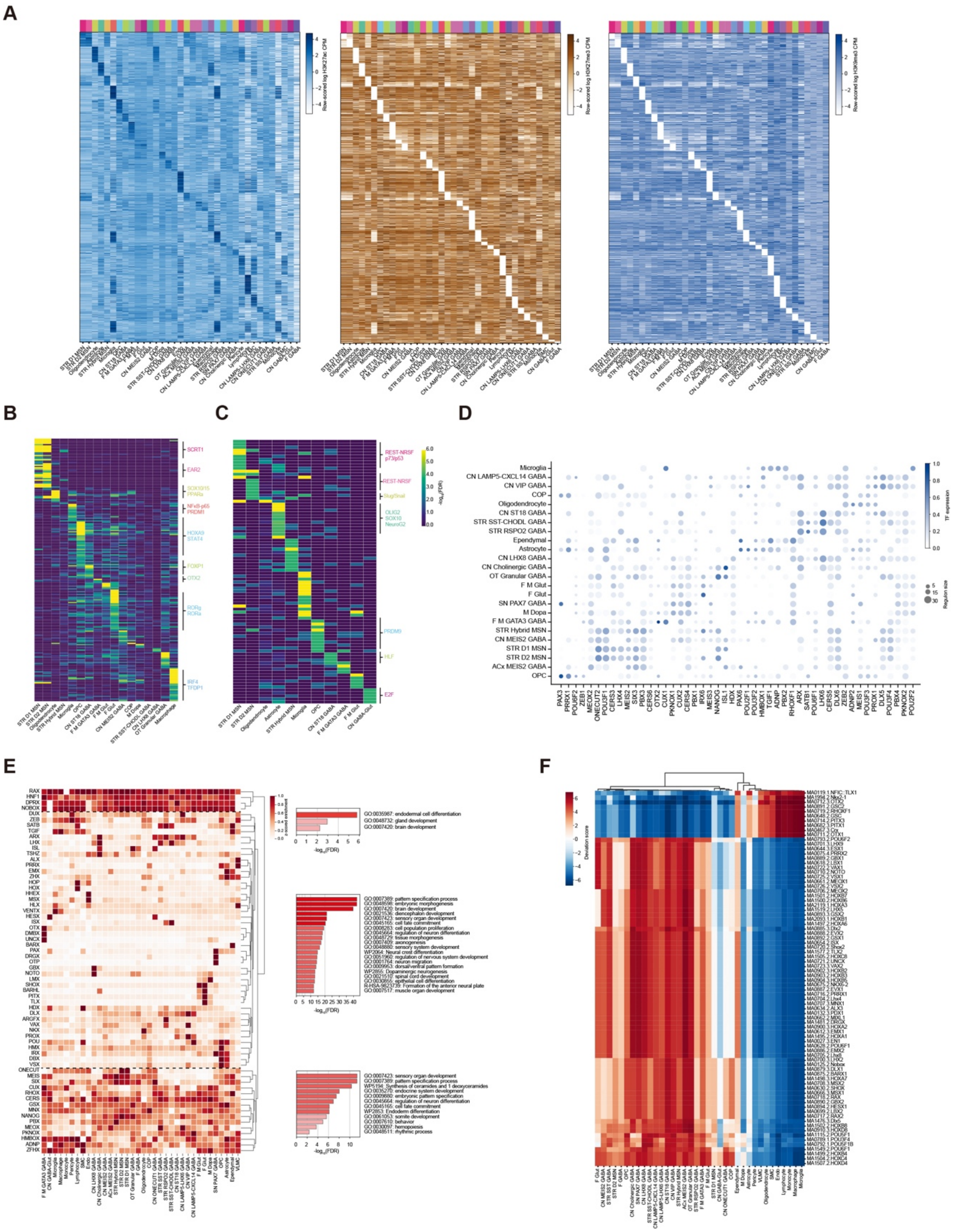


**Figure S3.2 Cell type-specific gene regulatory programs in the human basal ganglia.**
(A) Heatmaps showing three histone modification levels surrounding the TSS (± 2 kb) of cell-type-specific upregulated genes across 36 subclasses.
(B) TF motif enrichment in the top three ABC model-predicted enhancers linked to subclass-specific upregulated genes, with the corresponding H3K27ac signal shown as a heatmap in Figure 3B.
(C) TF motif enrichment in the putative cCREs linked to subclass-specific upregulated genes from the ABC model, selecting regions with relative depletion of H3K27me3 compared with other

subclasses. H3K27me3 signals at these cCREs are shown in Figure 3B.
(D) Clustermap showing homeobox TF eRegulons in basal ganglia. Dot size showing genes within each homeobox TF eRegulon in each subclass. Dot color showing homeobox TF gene expression level in each subclass.
(E) Homeobox TF gene expression patterns across 36 subclasses and hierarchical clustering that classified them into three subpopulations (left panels). GO enrichment analysis of homeobox TF genes from each subpopulation (right panels).
(F) The chromatin accessibility of the binding motifs of each homeobox TF across 36 basal ganglia subclasses.

A

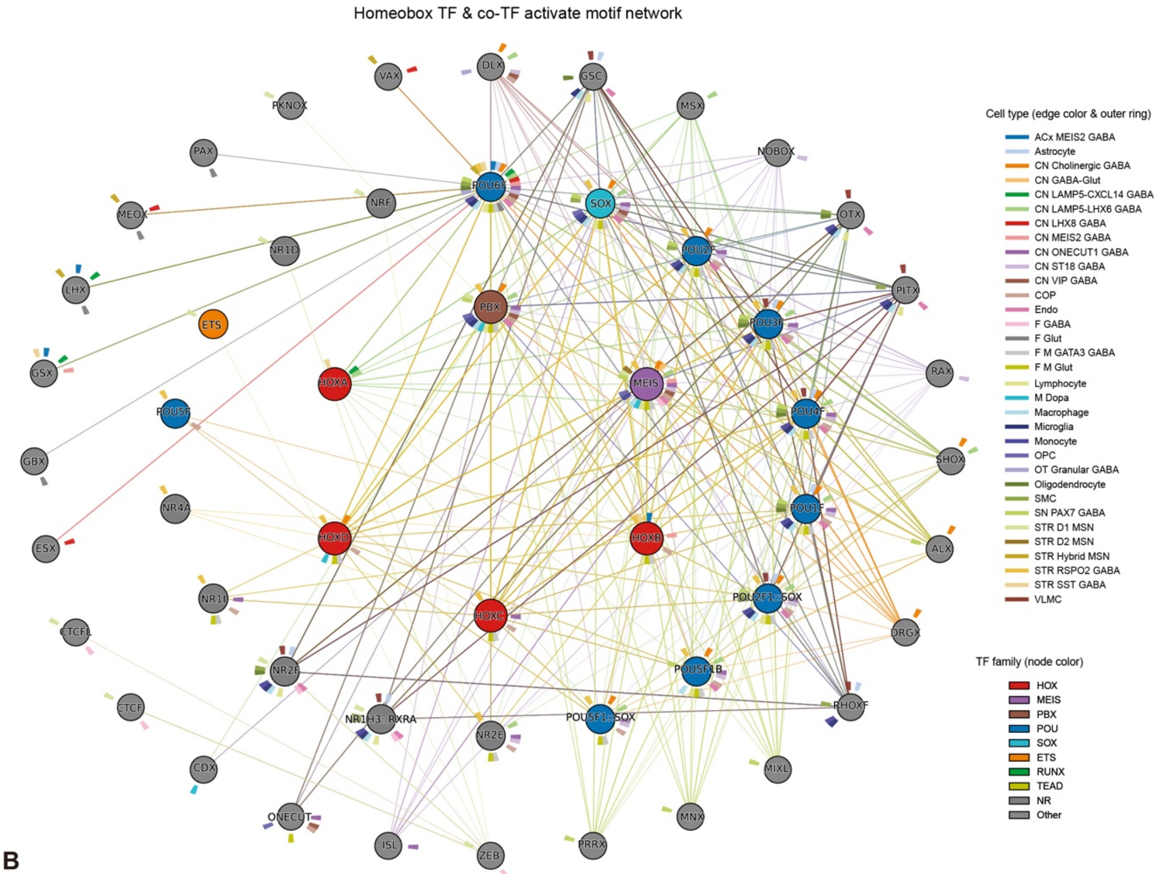


B

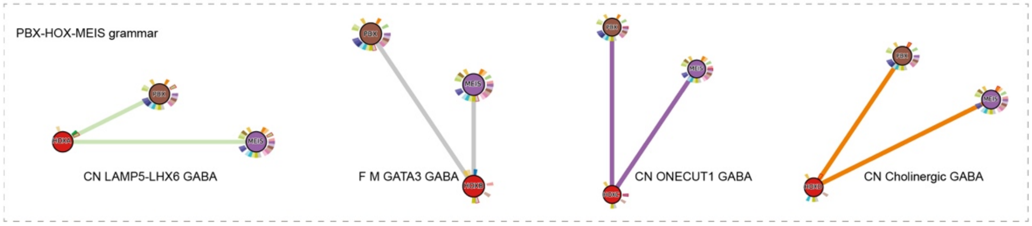


**Figure S3.3 The TF motif co-accessible networks of homeobox TF and co-TF across basal ganglia subclasses.**
(A) Network plots showing homeobox TF and co-TF motifs co-accessibility across 36 subclasses. Circles representing homeobox TFs and co-TFs, colored by subordinate TF families. Edges representing the subclass where homeobox TF and co-TF motifs are co-accessible.
(B) Co-accessibility grammar of PBX-HOX-MEIS motifs across four GABAergic neuron subclasses.

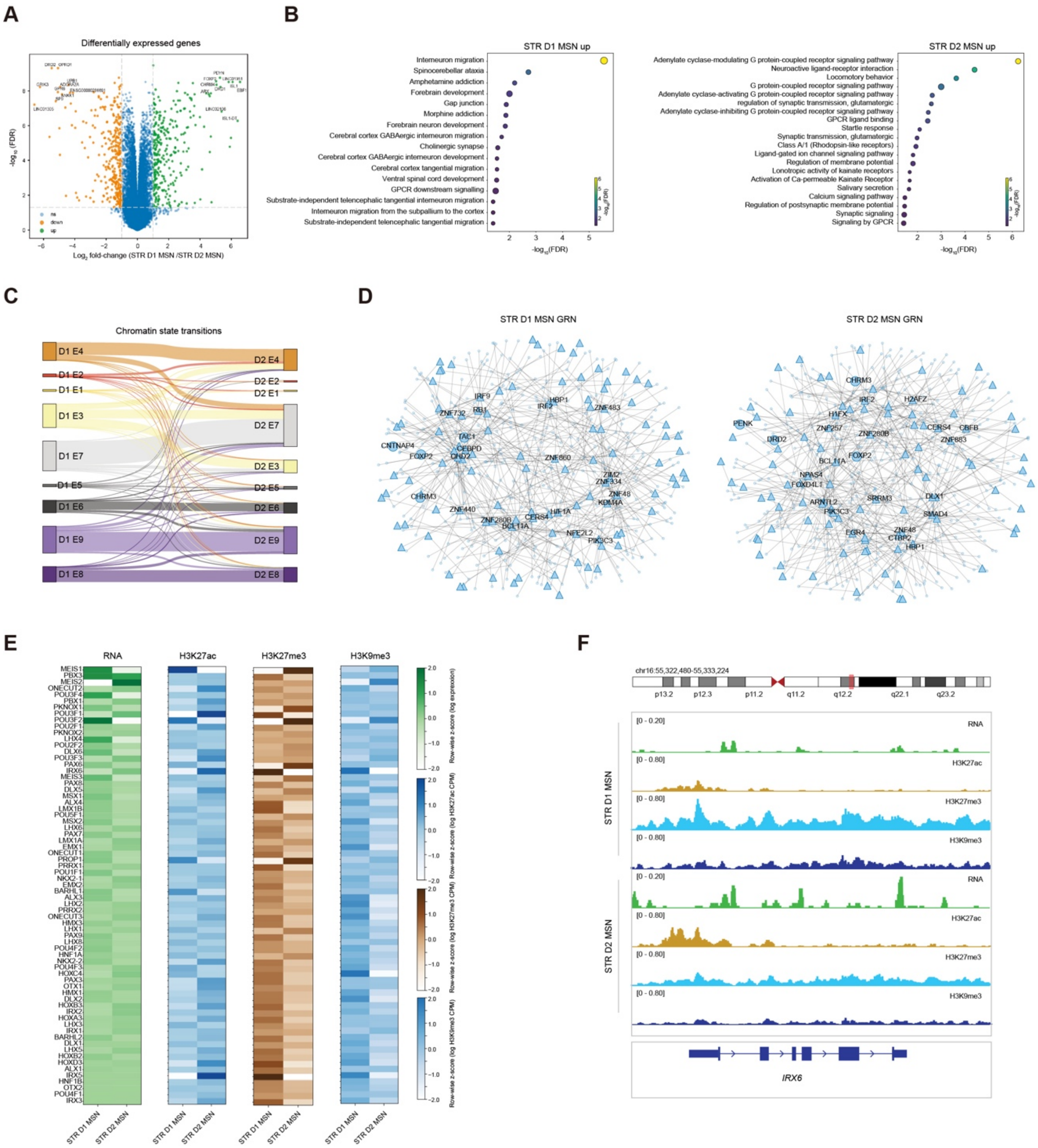


### Figure S3.4 Distinctive gene expression patterns and gene regulatory networks between STR D1 and D2 MSNs.

(A) Volcano plot representing the DEGs (|Log$_2$fold change (FC)| > 1.0 and false discover rate (FDR) < 0.01) between STR D1 and D2 MSNs with orange (upregulated in STR D2 MSNs) and green (upregulated in STR D1 MSNs) colors, where each point represents a gene.
(B) Dot plot showing top enriched GO terms from GSEA analysis of DEGs between STR D1 and STR D2 MSNs. Dot size reflects gene set size of GO terms.
(C) Sankey plot showing the chromatin state transitions between STR D1 and STR D2 MSNs.
(D) Network plots showing GRNs inferred for STR D1 MSNs (left) and STR D2 MSNs (right). Triangles represent TFs, and circles represent target genes.
(E) Heatmaps showing expression levels of homeobox TF genes and three histone modification levels surrounding the TSS (± 2 kb) between STR D1 and D2 MSNs.
(F) Genome browser track view at the *IRX6* locus as an example to show expression and epigenomic distinction in homeobox TF genes between STR D1 and D2 MSNs.

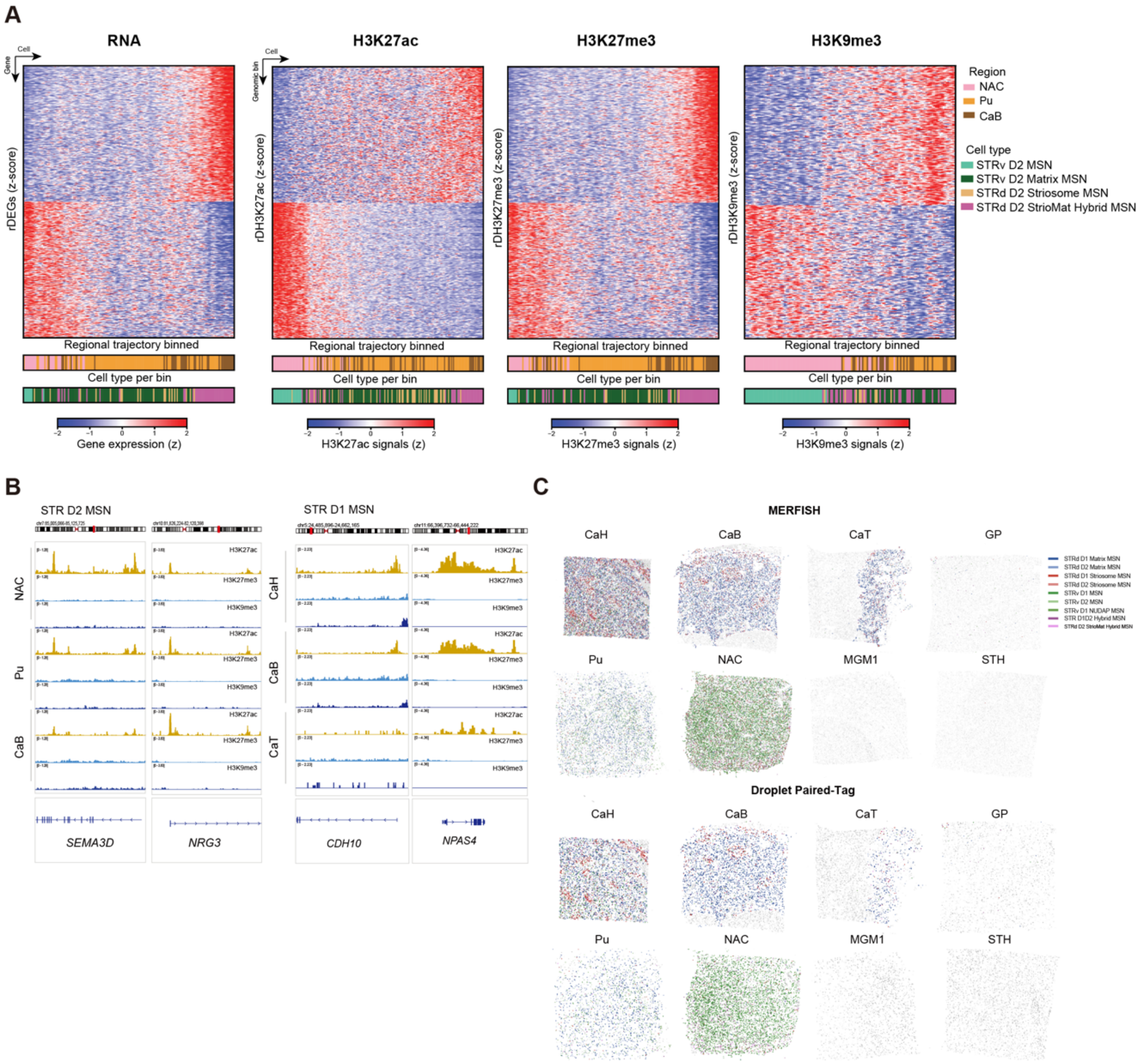


**Figure S4.1 Epigenomic gradients along basal ganglia regional axes.**

(A) V–D axis gradients in gene expression and histone modification signals of H3K27ac, H3K27me3 and H3K9me3 in 15-kb genomic bins for groups within STR D2 MSN subclass.

(B) Genome browser tracks showing histone modification gradient changes along the V-D and A-P axis.

(C) Top, the spatial maps of MERFISH identified MSN groups in each basal ganglia region. Bottom, the spatially mapped Droplet Paired-Tag cell atlas in eight basal ganglia regions. Colors represent imputed spatial locations of nuclei within each MSN group.

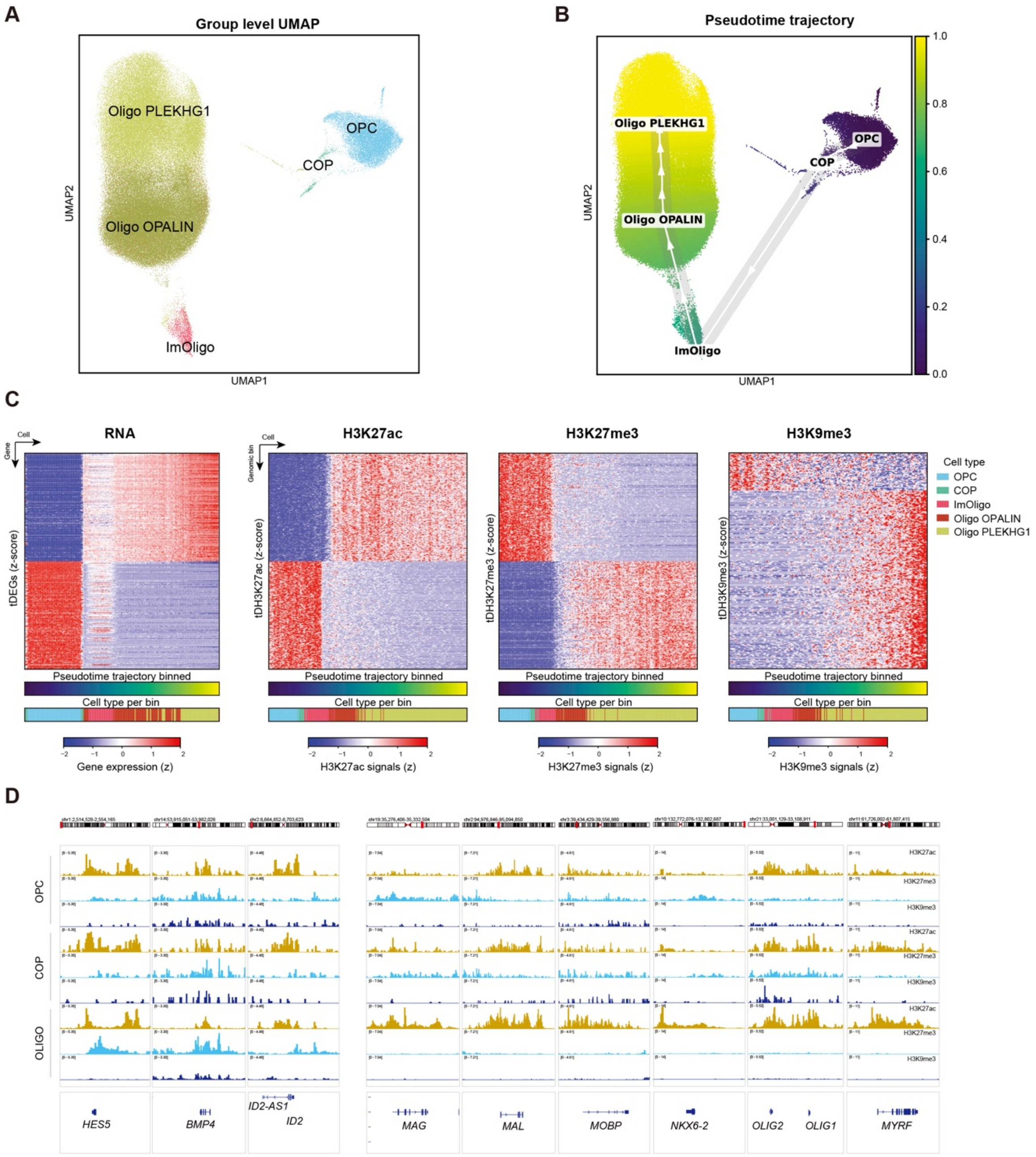


**Figure S4.2 Epigenomic gradient changes during OPC to oligodendrocyte differentiation**

(A) UMAP showing groups in oligodendrocyte lineages.

(B) UMAP embedding the trajectory of oligodendrocyte maturation. Each dot represents a single nucleus and is colored by pseudotime.

(C) Heatmaps showing the gene expression levels of differentially expressed genes, and histone modification levels of differentially modified genomic bins along the pseudotime trajectory.

(D) Genome browser tracks showing gradient changes of histone modification levels on oligodendrocyte myelination and differentiation relating genes.

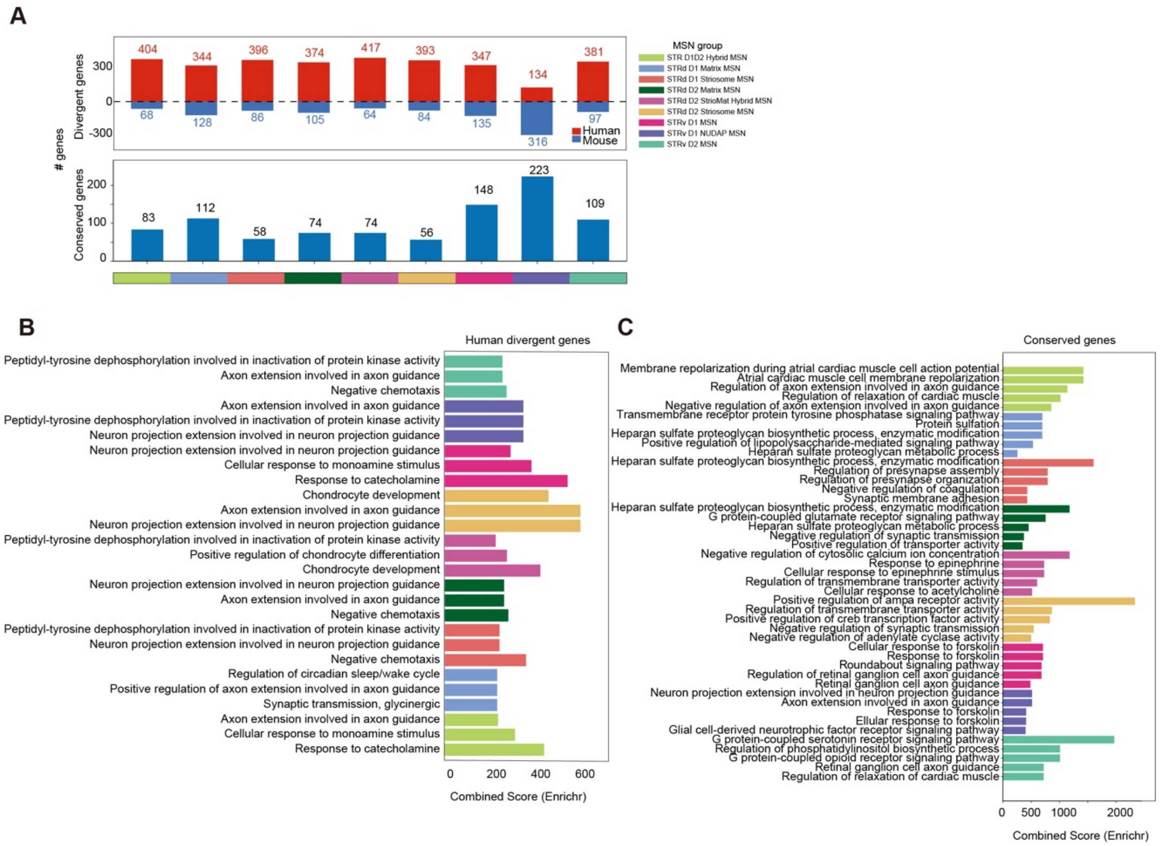


**Figure S5.1 Comparative analyses of transcriptome between human and mouse MSNs.**
(A) Bar plot showing number of conserved and divergent genes between human and mouse in each MSN group.
(B and C) Top three enriched GO terms from GSEA analysis for human up-regulated genes for each MSN group (B). Top five enriched GO terms from GSEA analysis for conserved genes between human and mouse for each MSN group (C). Bar plot showing combined score for each GO term.

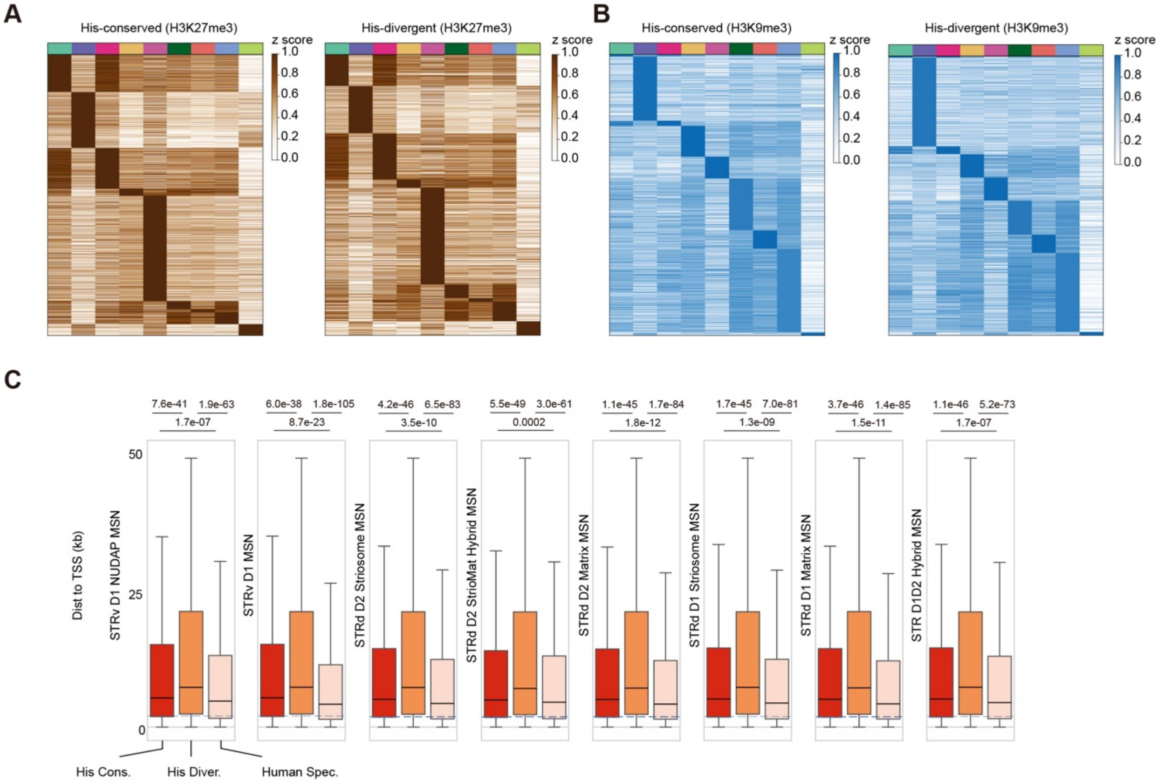


**Figure S5.2 Comparative analyses of histone modification between human and mouse MSNs.** (A and B) MSN group-biased conserved and divergent H3K27me3 (A) and H3K9me3 (B) peaks. (C) Box plots showing the distance of three categories of H3K27ac peaks to TSS in each MSN group. Hinges represent the 25th to 75th percentiles, the center line denotes the median, and whiskers represent the minimum and maximum values. Wilcoxon rank-sum Test.

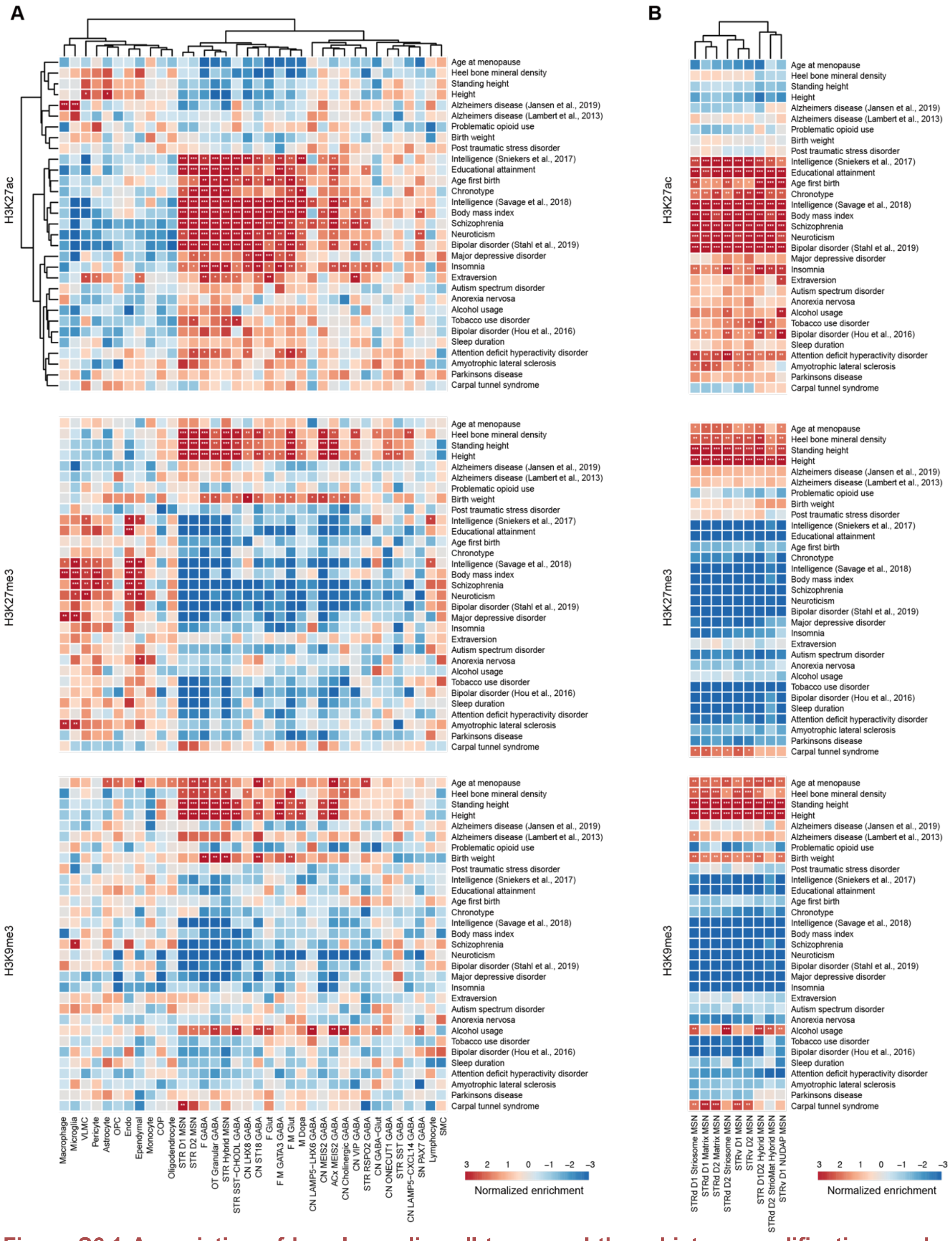


**Figure S6.1 Association of basal ganglia cell types and three histone modification marks with complex traits and diseases.**

(A) Heatmap showing LDSC analysis results of the risk variants associated with traits and diseases enrichment in three histone modification peaks from each major basal ganglia subclass. *FDR < 0.1; **FDR < 0.05; *** FDR < 0.01.

(B) Heatmap showing enrichment of risk variants associated with traits and diseases in three histone modification peaks from each MSN group. *FDR < 0.1; **FDR < 0.05; *** FDR < 0.01.

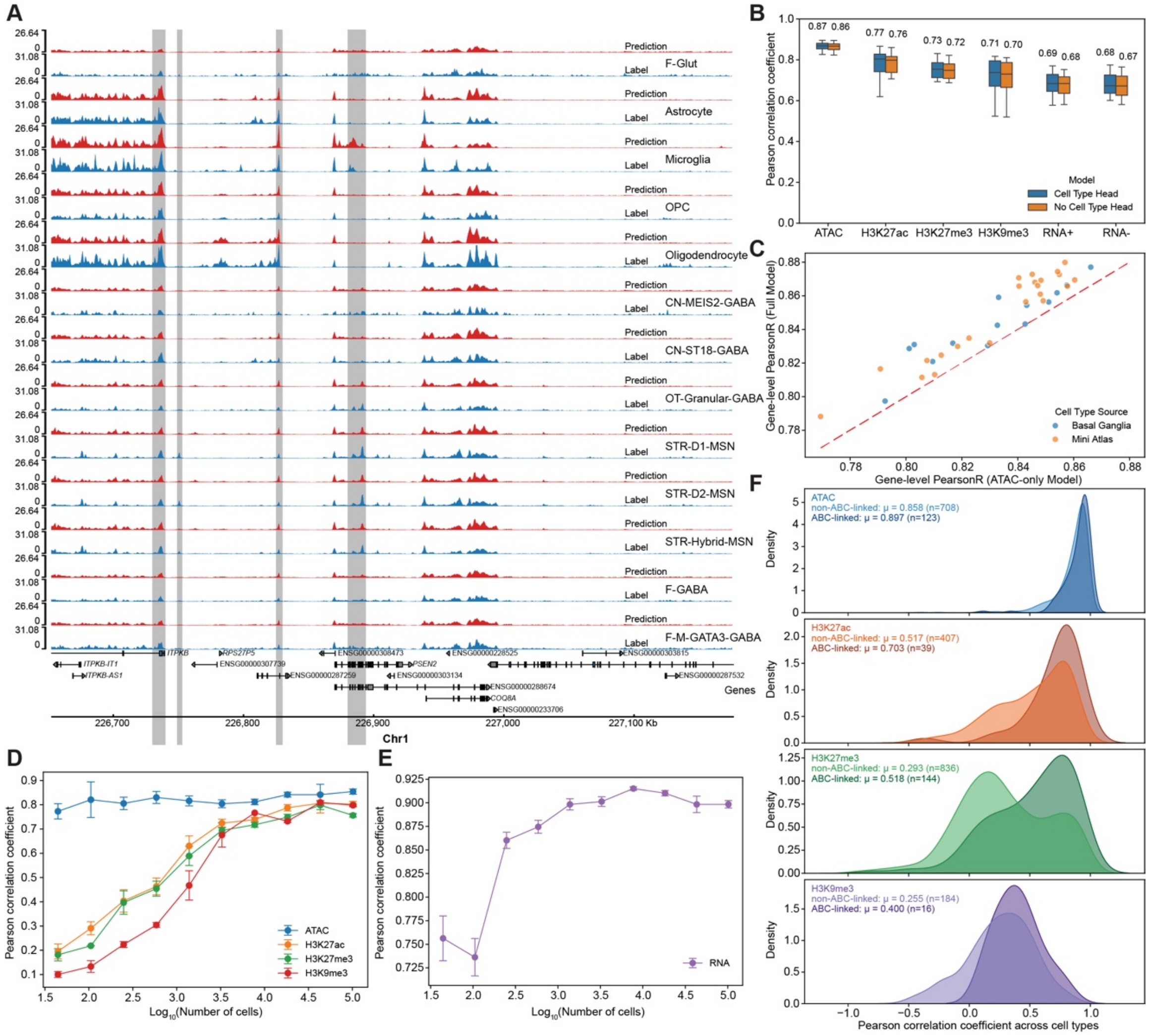

**Figure S7.1 Evaluation of deep learning model performance in predicting cell-type-specific epigenomic profiles and transcriptome.**

(A) Genome browser tracks showing observed (blue) and predicted (red) H3K27ac signals around the *PSEN2* locus. Cell-type-specific H3K27ac peaks are highlighted by a grey background.

(B) Performance comparison between different model architectures. Boxplots displaying Pearson correlation coefficients for the model utilizing the proposed cell-type-specific prediction head (blue) versus the standard cell-type-by-track architecture (orange). Performance is evaluated across chromatin accessibility (ATAC), different histone modification profiles, and gene expression (RNA+, forward strand; RNA-, reverse strand).

(C) Scatter plot comparing gene expression-level prediction accuracy (Pearson correlation) between the full model, trained on histone modification profiles, chromatin accessibility profiles and transcriptome, and the ATAC-only model, trained on chromatin accessibility profiles and transcriptome. Points colored by cell type source.

(D) Model prediction performance (Pearson correlation) for chromatin accessibility and histone modifications plotting as a function of the cell number per cell type.

(E) Gene expression prediction performance plotting as a function of the cell number per cell type.

(F) Distributions of Pearson correlation coefficients for predicted chromatin accessibility and three histone modification profiles across cell types. Performance is stratified by cCREs predicted to be linked to TSSs via the ABC model (dark shades) versus non-linked cCREs (light shades).

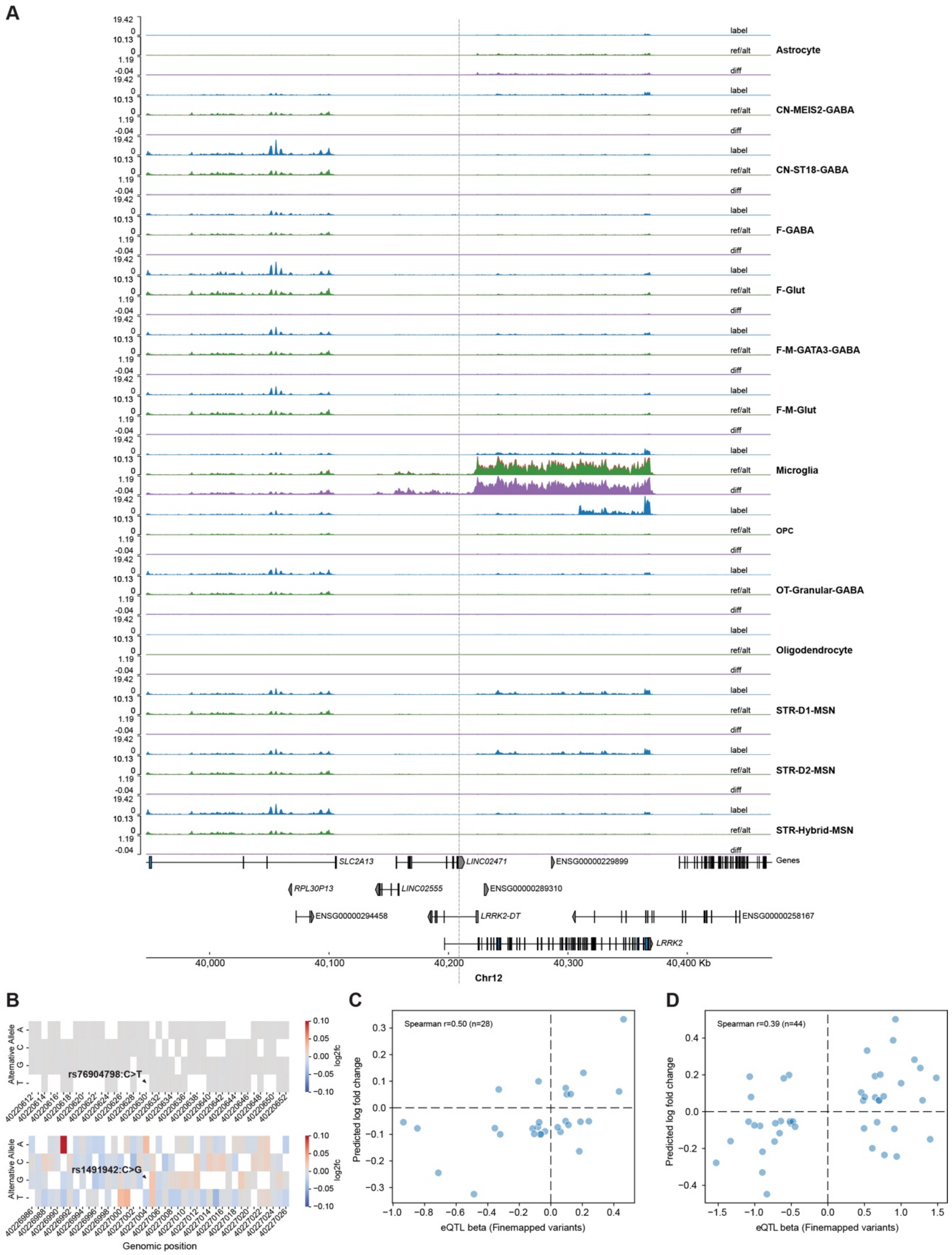


**Figure S7.2 Application of the deep learning model for predicting cell-type-specific non-coding variant effects.**

(A) Visualization of model predictions for the rs6581593 (C>T) variant across cell types in basal ganglia. Tracks displaying the predicted signal for the reference allele (Ref), the alternative allele (Alt), and the difference (Diff) between them.

(B) In silico saturation mutagenesis analysis of the *LRRK2* locus. The plots displaying the predicted

effect of all possible mutations ($log_2$ fold change in gene expression) within the genomic regions surrounding risk variants rs76904798 (C>T) and rs1491942 (C>G).
(C) Scatter plot showing comparison between model-predicted variant effect sizes (log fold change) and observed eQTL beta values for fine-mapped variants from the Single Brain database. The Spearman's correlation coefficient is shown.
(D) Scatter plot showing comparison between model-predicted variant effect sizes (log fold change) and observed eQTL beta values for fine-mapped variants in the basal ganglia (MetaBrain database). The Spearman's correlation coefficient is shown.
